# Integrated metabolomic and genomic insights into amino acid incorporation within the hybrid polyketide-alkaloid antibiotic TLN-05220

**DOI:** 10.1101/2025.10.30.685158

**Authors:** Jennifer L. Cordoza, Rachel A. Johnson, Andrew E. Whiteley, Calista A. Horta, Jayna C. Nicholas, Diana M. Owen, Laura Rodriguez-Velandia, Gordon T. Luu, Valentina Z. Petukhova, Jackson T. Calhoun, Laura M. Sanchez, Shaun M. K. McKinnie, Katharine R. Watts

**Affiliations:** University of California, Santa Cruz, Santa Cruz, CA, USA; California Polytechnic State University, San Luis Obispo, San Luis Obispo, CA, USA; University of Illinois Chicago, Chicago, IL, USA

## Abstract

Actinobacteria are a rich source of bioactive compounds and unique biosynthetic chemistry. *Micromonospora echinospora* subsp. *challisensis* NRRL 12255 produces the aromatic polyketide TLN-05220, which exhibits nanomolar activity against antibiotic-resistant human pathogens including vancomycin-resistant *Enterococcus faecalis* and methicillin-resistant *Staphylococcus aureus*. The pentangular polyphenol core of TLN-05220 is decorated with a piperazinone moiety, yet the enzymes responsible for the construction of this uncommon modification from amino acid precursors are unknown. Synthetic piperazinone-containing molecules have diverse antimicrobial, antiviral, anticancer, and anti-inflammatory bioactivity profiles, and determining biosynthetic routes for the assembly of this heterocycle may enhance drug discovery and medicinal chemistry efforts. We identified a putative TLN-05220 biosynthetic gene cluster (BGC) in the commercially available strain *M. echinospora* ATCC 15837 containing both type-I and type-II polyketide synthases, two predicted asparagine synthetase-like enzymes, and two genes (*tln*1 and *tln*5) that putatively encode pyridoxal 5’-phosphate (PLP)-dependent amino acid synthases. Stable isotopic feeding studies coupled with liquid chromatography–tandem mass spectrometry (LC–MS/MS), identified L-alanine, L-serine, and glycine as metabolic precursors of TLN-05220. Subsequent in vitro enzymology established that Tln1 is a PLP-dependent alanine racemase, while Tln5 performs a stereoselective β-substitution reaction of *O*-phospho-L-serine with a preferential D-alanine nucleophile. Alanine racemization and Tln5 pseudodipeptide L-serine-Cβ-*N*-D-alanine (D,L-PDP) incorporation into TLN-05220 were further supported using deuterated intermediates and mass spectrometry techniques. Establishing the enzymes that catalyze amino acid installation within TLN-05220 inspires further biosynthetic discovery and engineering while enabling the biocatalytic syntheses of novel amino acid-containing polyketide antibiotics.

**Graphical Abstract:** 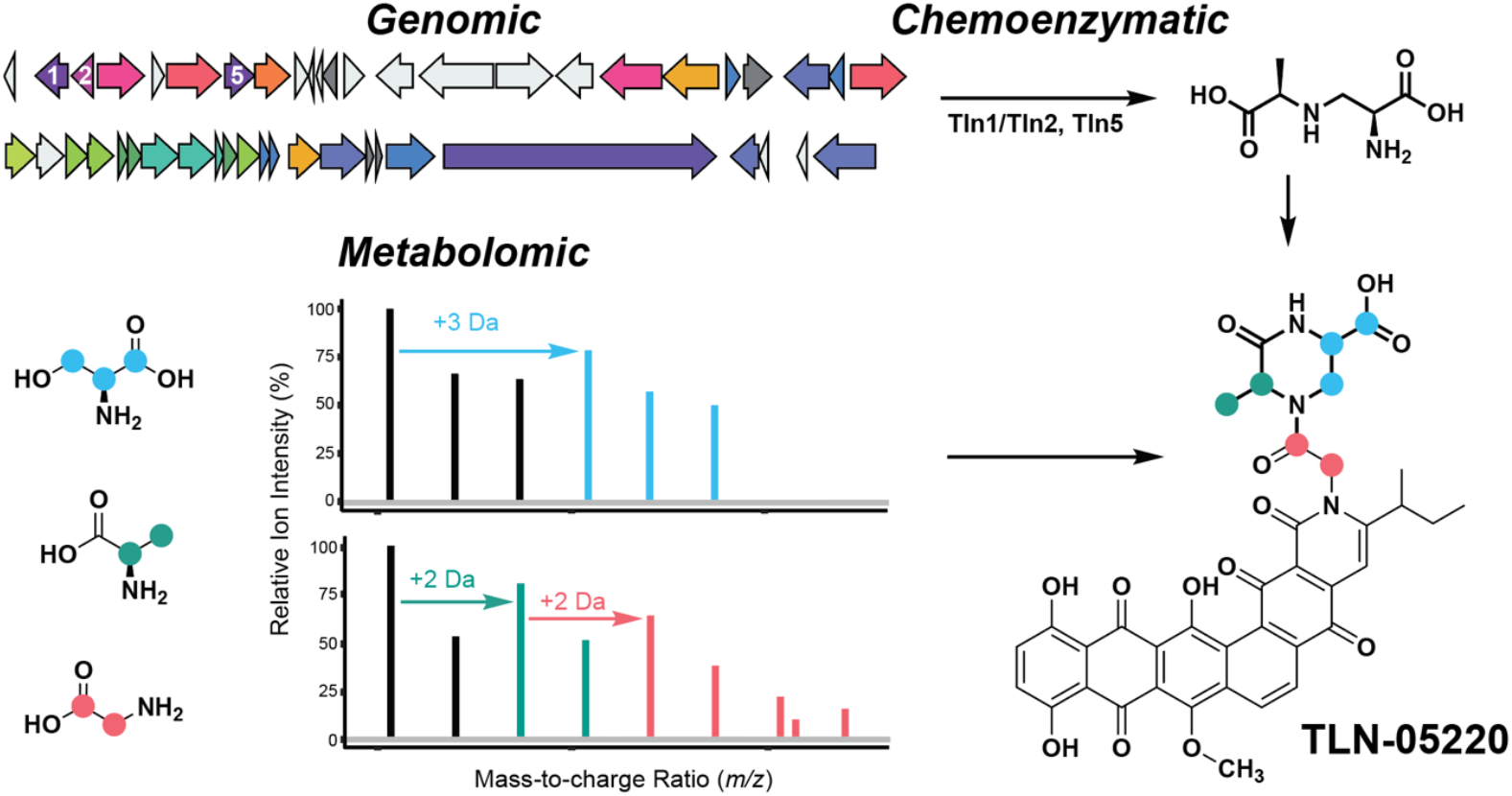

## Introduction

In bacteria, aromatic polyketides are often biosynthesized by type II polyketide synthases (PKSs). Type II PKS systems utilize dissociable domains, including the essential acyl carrier protein (ACP), acyltransferase (AT), and characteristic ketosynthase–chain length factor (KS–CLF) heterodimer, to iteratively catalyze controlled chain elongation by Claisen condensations.^1^ The echinosporamicin-type compounds, including echinosporamicin 1–5,^2^ bravomicin A,^3,4^ and TLN-05220 and TLN-05223,^4^ are a class of pentangular polyphenol aromatic polyketides (**Figure 1**). Discovered from various Micromonosporaceae, this compound class shows antimicrobial activity against Gram-positive bacteria in the nanomolar range, maintaining activity against strains that are vancomycin- and methicillin-resistant (**Table S1**).^2–4^

**Figure 1.**
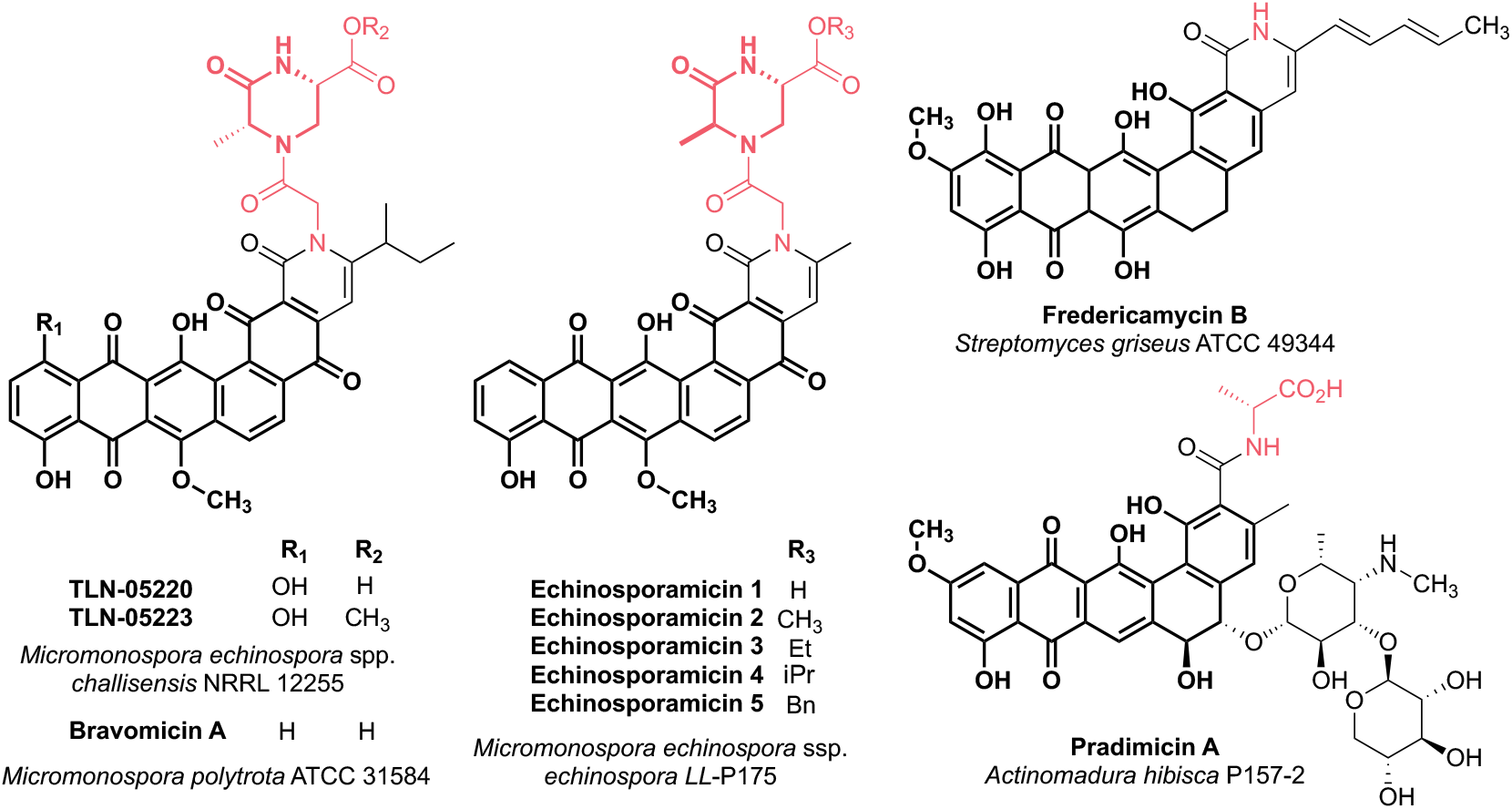
Chemical structures of the echinosporamicin-type molecules and related nitrogen-containing type II polyketides. The pentangular polyketide (bolded in black) is modified by the addition of one or more atoms derived from amino acid(s) (pink). The piperazinone moiety bolded in pink for TLN-05220 and TLN-05223,^4^ bravomicin A,^3,4^ and echinosporamicin 1–5.^2^ Fredericamycin B and pradimicin A lack a piperazinone but contain amino acid-derived features (pink).^5–7^ Strains from which the compounds were originally discovered are listed below the names of each compound series. Note that bravomicin A is depicted as the revised structure as suggested by Banskota et al,^4^ and echinosporamicin stereochemistry is shown as suggested by He et al.^2^

The echinosporamicin-type molecules are decorated with a piperazinone ring, which is hypothesized to be composed of amino acid precursors (**Figure 1**).^4^ The piperazinone (ketopiperazine) group is characterized as a 6-membered heterocycle containing a single endocyclic amide and a secondary amine (**Figures 1, S1**). Diketopiperazine cyclic dipeptides are biosynthesized using either a cyclic dipeptide synthetase (CDPS) or by a non-ribosomal peptide synthetase (NRPS);^8,9^ a desaturated pyrimidinone heterocycle has been characterized within oxepine-pyrimidinone-ketopiperazine (OPK) natural products following NRPS cyclization and subsequent oxidative tailoring.^10,11^ Despite the structural similarities, the biosynthetic steps yielding a piperazinone moiety have been underexplored due to the relative rarity of this structural moiety (**Figure S1**).

Through genome scanning in the TLN-05220 producer *Micromonospora echinospora* ssp. *challisensis* NRRL 12255, Thallion Pharmaceuticals identified a putative 39.8 kb TLN-05220 biosynthetic gene cluster (BGC) (**Figure S2**).^4^ The reported cluster (MIBiG BGC0001062)^12^ contains genes encoding: a minimal type II PKS (ACP, AT, and KS– CLF); two ketoreductases; five monooxygenases; a multicopper oxidase; three cyclases; a 3-hydroxyisobutyrate dehydrogenase; an *O*-methyltransferase; an amidotransferase; a ketoacyl-ACP synthase III and acyltransferase domain-containing protein; in addition to a type I PKS module.^4^ Experimental evidence that links this BGC to the TLN-05220 metabolite have yet to be reported,^12^ however, bioinformatic studies reveal high sequence similarity between the CLF within the cluster to those within other pentangular polyketide biosynthetic pathways,^13,14^ strongly supporting the assignment to biosynthesize the polyketide structure within TLN-05220. Rationally, Banskota et al. hypothesized that a NRPS-encoding gene elsewhere in the genome was responsible for the biosynthesis of the piperazinone.^4^ To our knowledge, a hybrid type II PKS-NRPS biosynthetic pathway has never been reported, thus we were motivated to further investigate TLN-05220 biosynthesis.

Here, we identify a new TLN-05220 producer, *Micromonospora echinospora* ATCC 15837. Stable isotope labeling (SIL) experiments on semi-solid agar, analyzed by liquid chromatography–tandem mass spectrometry (LC–MS/MS) and matrix-assisted laser desorption/ionization–imaging mass spectrometry (MALDI–IMS), show that L-serine, L-alanine, and glycine are biosynthetic precursors. In vitro assays reveal that two pyridoxal 5’-phosphate (PLP)-dependent enzymes, Tln1 and Tln5, encoded by two genes that lie upstream of the originally proposed TLN-05220 gene cluster (MIBiG BGC0001062; GenBank accession number FJ915123)^4,12^ participate in the biosynthesis of the piperazinone. We identify Tln1 as an alanine-selective racemase and establish Tln5 as an unusual PLP-dependent enzyme that catalyzes the β-substitution of *O*-phospho-L-serine (L-OPS) with D-alanine to form L-serine-Cβ-*N*-D-alanine, a novel pseudo-dipeptide (D,L-PDP). Additional in vivo SIL experiments with deuterated D,L-PDP further supports the involvement of this intermediate in TLN-05220 biosynthesis. Together, these findings redefine the biosynthetic boundaries of this unusual polyketide-alkaloid antibiotic and open new opportunities for further genome mining, genetic engineering, and biocatalytic application of this unique amino acid manipulation enzymology.

## Results

Neither the full genome sequences nor the identified *Micromonospora* producer strains of any of the echinosporomicin-type polyketides are publicly available. Thus, we used the critical KS–CLF gene sequences from the TLN-05220 cluster (MIBiG BGC0001062)^4,12^ as a query to search for potential TLN-05220 producers from other Micromonosporaceae. From the full genome sequences available at Joint Genome Institute (JGI), we identified *M. echinospora* ATCC 15837 to have a 96–97% identical KS–CLF sequence (**Table S2**). Using antiSMASH,^15,16^ we analyzed the ATCC 15837 genome for candidate BGCs and identified a 74.5 kb cluster containing the KS–CLF gene we used for our initial query. Twenty-six of the thirty genes in the previously reported TLN-05220 cluster (MIBiG BGC0001062) encode for protein homologs with greater than 30% sequence identity in the cluster of interest found in ATCC 15837 (**Figure S2**). In contrast to MIBiG BGC0001062, the putative TLN-05220 BGC from *M. echinospora* ATCC 15837 includes 19.9 kb that encodes 17 additional upstream genes and 18.6 kb that encodes 19 additional downstream genes flanking the original boundaries. Using biosynthetic rationale, we propose a 60.0 kb TLN-05220 BGC, which extends the bounds of the cluster to include all 17 upstream genes and 2 downstream genes (**Figure 2** and **Table S3**). The majority of the remaining downstream genes (12 out of 17) annotated as hypothetical proteins/proteins of unknown function and were excluded from the redefined TLN-05220 BGC (**Figure S2**).

**Figure 2.**
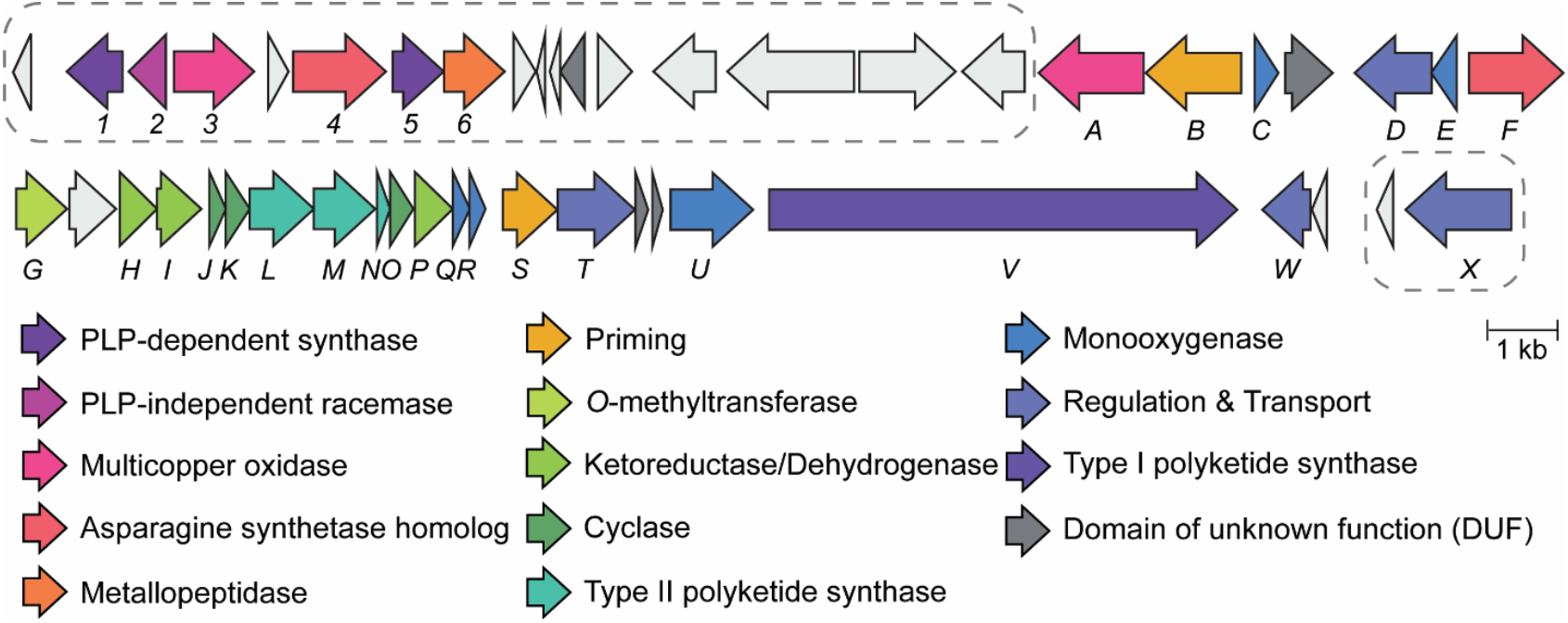
Proposed extension of boundaries for the *M. echinospora* ATCC 15837 TLN-05520 biosynthetic gene cluster. Genes within the range of *tlnA*–*tlnW* aligned well with MIBiG BGC0001062, the previously reported for TLN-05220 cluster (see **Figure S2**). The dashed boxes encompass 17 upstream genes and 2 downstream genes that we hypothesize are additionally important in TLN-05220 biosynthesis. Putative functions of encoded enzymes are color coded.

To validate *M. echinospora* ATCC 15837 as a TLN-05220 producer, we cultured it on N-Z-Amine agar plates and analyzed the metabolic extract using targeted LC–MS/MS. The TLN-05220 molecule was observed at 738.1923 *m*/*z* (calc. [M+H]^+^ *m/z* 738.1930; **Figure 3**). By tandem MS, the protonated molecule fragmented and presented the characteristic fragment ions at a *m*/*z* of 580.1226 and 552.1307 (expected [M+H]^+^ *m/z* 580.1238 and 552.1289, respectively; **Figure S3**), indicating dissociation of the piperazinone ring. The proposed carbon monoxide eliminations are well documented from multiple natural product classes.^17^ To further probe production of TLN-05220 in ATCC 15837, MALDI– IMS analyses were conducted on semi-solid agar plates. Ion imaging showed diffusion of TLN-05220 into the agar after 16 days by detection of the sodium- and potassium-adducts using this methodology ([M+Na]^+^ and [M+K]^+^ of 760 and 776, respectively; **Figure S4**). With successful production and detection of the metabolite of interest, we sought to test if glycine, L-alanine, and L-serine are incorporated into TLN-05220, pointing to their significance as biosynthetic precursors. For this bacterium, we did not obtain sufficient growth on minimal media, as is traditionally utilized for stable isotope labeling (SIL) studies. Instead, we supplemented N-Z-Amine agar plates with ^13^C-labeled amino acids in three distinct conditions: 1,2,3-^13^C_3_-L-serine; 2,3-^13^C_2_-L-alanine; and both 2,3-^13^C_2_-L-alanine and 1,2-^3^C_2_-glycine. *M. echinospora* ATCC 15837 was also grown on N-Z-Amine agar control plates containing the respective unlabeled amino acid and all plates were incubated for 5 days. LC–MS analyses of the extracted bacteria show distinct isotopic shifts compared to the media control (**Figure 3A**). When adding 1,2,3-^13^C_3_-L-serine to the media, a new TLN-05220 isotopologue was observed at 741.2026 *m*/*z* (**Figure 3B**). This ion is 3 Da heavier than the TLN-05220 [M+H]^+^ ion in the media only control (738.1926 *m*/*z*), indicating incorporation of three carbon-13 atoms from 1,2,3-^13^C_3_-L-serine into the TLN-05220 backbone. A significant increase in the abundance of the 740.2002 *m*/*z* TLN-05220 isotopologue is observed when feeding ATCC 15837 2,3-^13^C_2_-L-alanine, coupled with a decrease in the 738.1926 *m*/*z* abundance compared to the media control (**Figure 3C**). The 2 Da shift of the observed TLN-05220 [M+H]^+^ ion supports 2,3-^13^C_2_-L-alanine incorporation into the molecule. When feeding in both 2,3-^13^C_2_-L-alanine and 1,2-^13^C_2_-glycine, the +2 Da TLN-05220 isotopologue (740.1993 *m*/*z*) that is observed with just 2,3-^13^C_2_-L-alanine alone decreases, and the abundance of a 742.2068 ion increases supporting two additional carbon-13 atoms were incorporated into TLN-05220 from 1,2-^13^C_2_-glycine (**Figure 3D**). Taken together, these results provide strong evidence that L-alanine, L-serine, and glycine are all amino acid precursors of TLN-05220 production within the ATCC 15837 strain (**Figure 3E**).

**Figure 3.**
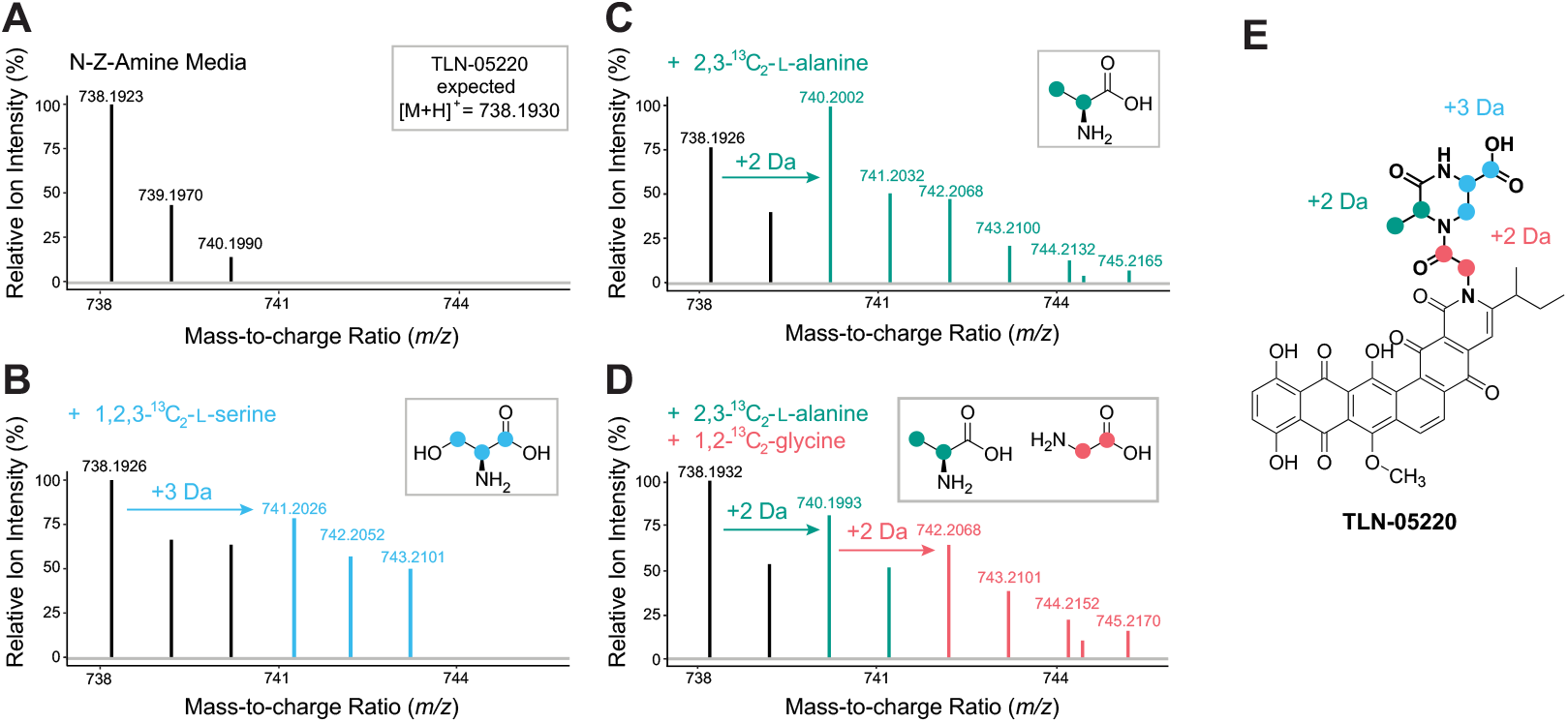
Stable isotope labeling analyses of TLN-05220 via LC–MS. Extract of *M. echinospora* ATCC 15837 grown on (**A**) N-Z-Amine A media control compared to media with carbon-13 labeled amino acids: **(B)** 1,2,3-^13^C_3_-L-serine (blue), (**C**) 2,3-^13^C_2_-L-alanine (teal), and (**D**) both 2,3-^13^C_2_-L-alanine (teal) and 1,2-^3^C_2_-glycine (pink). The TLN-05220 [M+H]^+^ ion is expected at a mass-to-charge (*m*/*z*) ratio of 738.1930 (<1 ppm error of observed *m*/*z* values). Peaks observed due to labeling are color coded and observed Dalton (Da) shifts due to ^13^C incorporation are indicated with arrows. Ion intensities are relative to the most abundant *m*/*z* ratio observed in each mass spectra window. The positions of each carbon-13 label are shown within the structure of each amino acid as respectively colored circles (panels **B–D**), and colored labels from each amino acid correspond to (**E**) the proposed isotopically labeled positions within the TLN-05220 metabolite.

Adenylation (A) domains within NRPSs select for amino acid substrates, suggesting that a NRPS may be responsible for incorporation of glycine, L-alanine, and L-serine into TLN-05520 as previously predicted.^4^ However, analyses of the 46 A-domains within NRPS and NRPS-like enzymes in the ATCC 15837 genome revealed no enzyme that is bioinformatically predicted to accept all three of the proposed amino acid TLN-052250 precursors (**Table S4**). The lack of NRPS candidate(s) initiated our search for alternative candidate piperazinone biosynthetic genes that utilize amino acid precursors. We took notice of several genes upstream within the newly expanded TLN-05220 BGC region, namely *tln1, tln4*, and *tln5*, due to their bioinformatic annotations associated with amino acid metabolism (**Figure 2, Tables S3** and **S5**).

Both Tln1 and Tln5 are classified as members of the pyridoxal 5’-phosphate (PLP)-dependent cysteine synthase/cystathionine β-synthase protein family (InterPro IPR050214,^18^ **Table S5**); Tln4 bioinformatically annotates as an asparagine synthetase involved in ligation reactions and exhibits high sequence identity to another amino acid ligase homolog in the proposed cluster, TlnF (42% pairwise identity). With these insights, we revised our TLN-05220 biosynthetic hypothesis to rationalize a non-NRPS incorporation of the amino acid precursors identified from stable isotope feeding (**Figure S5**). Retrobiosynthetic disconnection of the three amide bonds liberates glycine and a novel Ala-Ser derived pseudo-dipeptide (PDP) from the pentangular polyphenol polyketide precursor (**Figure 4**). Subsequent cleavage of the PDP β-carbon–nitrogen bond justified the remaining two primary metabolites, L-alanine and L-serine. In the forward biosynthetic direction, enzymes within the IPR050214 family catalyze important primary metabolic β-substitution reactions using native or *O*-functionalized serine.^19,20^ This reactivity is also precedented in microbial secondary metabolism,^21,22^ leading us to propose that Tln1 and Tln5 catalyze β-substitution reactions analogous to those of known PLP-dependent enzymes such as SbnA,^23^ CmnB,^24^ and AMA synthase,^25^ mediating incorporation of amino acids into the piperazinone ring of TLN-05220 via an analogous PDP. Furthermore, Tln4 and TlnF, both bearing *N*-terminal serines characteristic of ligase-type asparagine synthetases, may catalyze amino acid ligation steps similar to PdmN in pradimicin biosynthesis (**Figures 1, S6**, and **S7**).^6,7^ By contrast, the fredericamycin amidotransferase FdmV^5^ contains an *N*-terminal cysteine associated with amine transfer rather than ligation, underscoring the likely ligase function of Tln4 and TlnF. Together, we reasoned that Tln1, Tln4, Tln5, and TlnF may orchestrate the non-NRPS incorporation of glycine and an Ala-Ser-derived PDP during TLN-05220 assembly (**Figure S5**).

**Figure 4.**
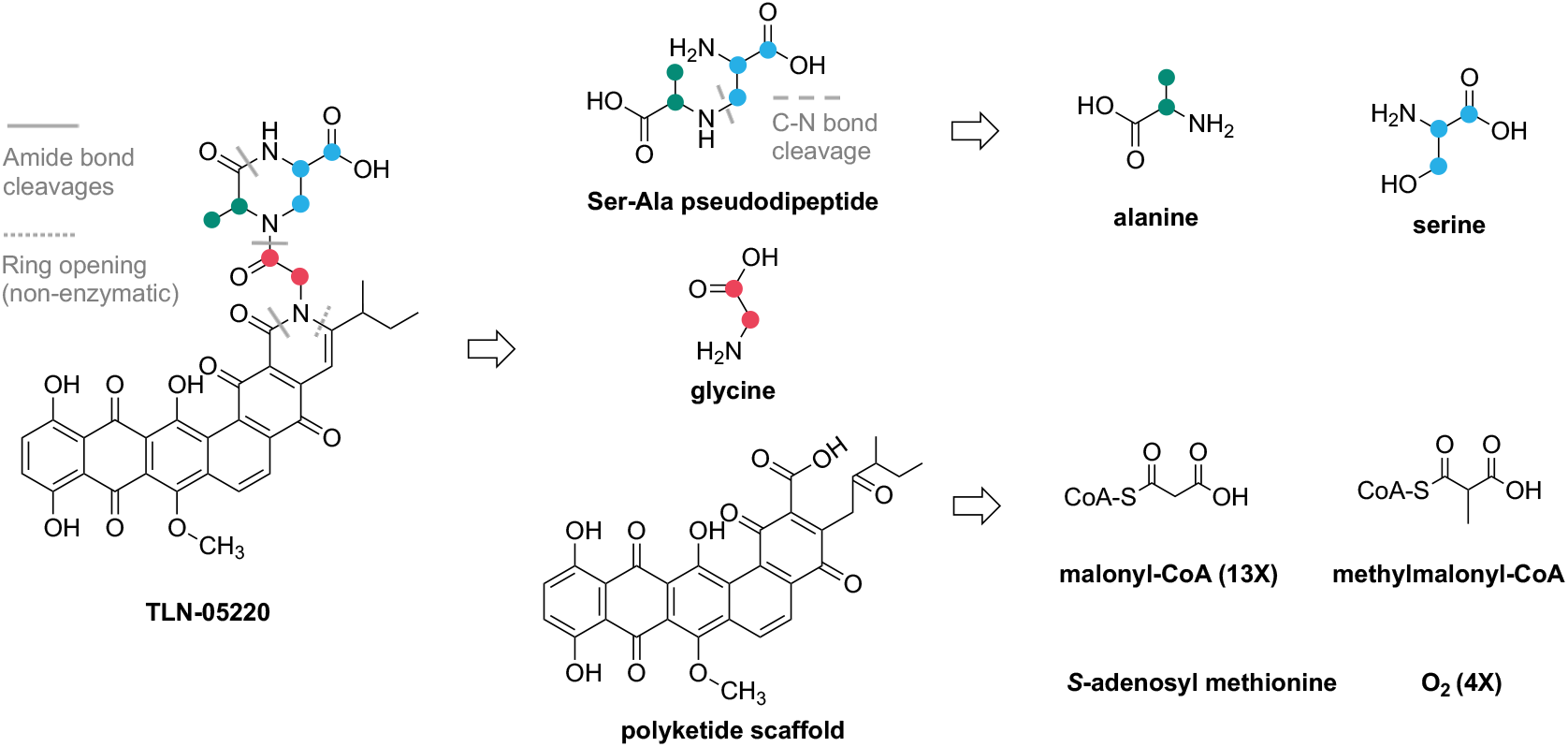
Retrobiosynthesis of TLN-05220 based on ^13^C stable isotope incorporation experiments and comparison to related biosynthetic pathways. Putative amide bond cleavages (solid lines) and non-enzymatic ring opening (dotted line) disconnect the polyketide scaffold from glycine (pink) and a pseudo-dipeptide (PDP), which can be further dissected into alanine (teal) and serine (blue) metabolites through C–N bond cleavage (dashed line). The polyketide scaffold is deconstructed into malonyl-CoA, methylmalonyl CoA, *S*-adenosylmethionine and molecular oxygen precursors.

To interrogate the role of these candidate genes, we obtained *Escherichia coli* (*E. coli*) codon-optimized sequences of both *M. echinospora* ATCC 15837 *tln1* and *tln5* genes and generated pET28a constructs for heterologous expression of each *N*-terminal His_6_-tagged protein. His_6_-Tln1 and His_6_-Tln5 (hereon Tln1 and Tln5) were purified using Ni-NTA affinity chromatography and size exclusion chromatography in respective yields of 9 and 80 mg/L of culture expressed (**Figure S8**). Both Tln1 and Tln5 co-purified with their cognate PLP cofactor, confirmed from UV-vis spectroscopic identification of the 420 nm internal aldimine peak (**Figure S9**). To test our retrobiosynthetic hypothesis that either Tln1 or Tln5 could catalyze the formation of an Ala-Ser PDP, 5 mol% of either PLP-dependent enzyme was incubated with putative substrates *O*-phospho-L-serine (L-OPS) and either D- or L-alanine for 1 h to assess activity in vitro. Enzyme assays were derivatized using 1-fluoro-2,4-dinitrophenyl-5-L-alanine amide (L-FDAA, Marfey’s reagent) and analyzed by ultra-performance liquid chromatography mass spectrometry (UPLC-MS) (**Figure 5A**). When Tln5 was incubated with L-OPS and either L-alanine or D-alanine, we observed the appearance of a 429.13 ion, corresponding to a Marfey-derivatized Ala-Ser PDP. However, the assay containing D-Ala produced a more intense signal for the derivatized product compared to L-Ala (**Figure 5A**). The Marfey-derivatized PDP ion was not observed in the equivalent Tln1 sample, in the no enzyme control conditions, or in the absence of either of the two amino acid substrates (**Figure 5A** and **S10**). Incubation of Tln5 with D-OPS and D-alanine under equivalent reaction conditions failed to yield a *m/z* signal for the putative Marfey-derivatized PDP ion, highlighting the stereospecificity of Tln5 for its serine donor (**Figure S11**). Replacement of L-OPS with alternative *O*-substituted L-serine molecules, including: *O*-acetyl-L-serine (L-OAS); *O*-succinyl-L-serine (L-OSS); or L-serine, showed significant reduction (L-OAS and L-OSS) or complete loss (L-serine) of Marfey-derivatized PDP [M+H]^+^ ion signal compared to a Marfey-derivatized L-glutamate standard, showing that Tln5 prefers L-OPS (**Figure S12**). We were able to recapitulate these results in an NMR-based assay with Tln5 and amino acid substrates (**Figures S13, S14**). NMR signals indicative of the new secondary amine group (δ 3.6 and δ 3.3) in the putative Ala-Ser PDP were observed and orthogonally supported the selectivity results obtained from the UPLC-MS experiments.

**Figure 5.**
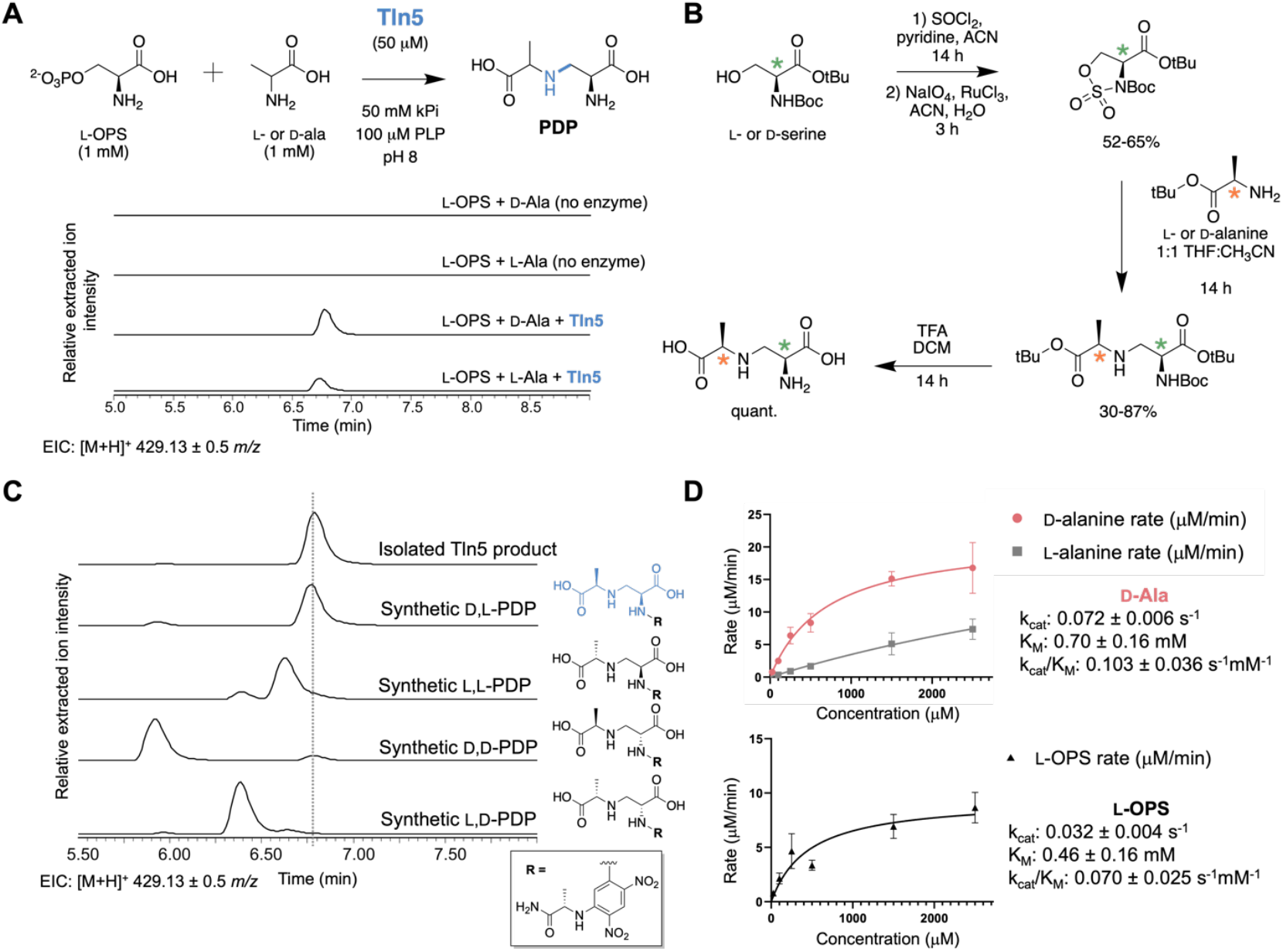
Tln5 generates D,L-pseudo-dipeptide, L-serine-Cβ-*N*-D-alanine (D,L-PDP). (**A**) Tln5 catalyzes a β-substitution reaction between *O*-phospho-L-serine (L-OPS) and either alanine enantiomer to generate a Ser-Ala pseudo-dipeptide (PDP) in vitro. Samples were derivatized with 1-fluoro-2,4-dinitrophenyl-5-L-alanine amide (L-FDAA) and analyzed by UPLC-MS. Relative intensities for extracted ion chromatograms in positive mode for Marfey-derivatized PDP ([M+H]^+^ 429.13 ± 0.50 *m/z*). (**B**) Chemical syntheses of four possible PDP diastereomers (D,D-PDP, D,L-PDP, L,L-PDP, and L,D-PDP). Positions of the varied stereocenters are noted with asterisk on the serine (green) and alanine (orange). (**C**) L-FDAA derivatization of each chemically synthesized PDP diastereomer and separation by UPLC-MS. The major chemoenzymatically synthesized product from Tln5 is corroborated to have a D,L-PDP stereochemistry. (**D**) Kinetic characterization of Tln5 with its preferred in vitro substrates D-alanine (top, pink line) and L-OPS (bottom, black line). Under equivalent reaction conditions, L-alanine (top, grey line) failed to reach maximum velocity (V_max_), suggesting L-alanine is a less preferred substrate for Tln5.

To unambiguously assign the stereochemistry of the major in vitro PDP product, we chemically synthesized all four possible diastereomers by adapting an established route used for the syntheses of AMA derivatives (**Figure 5B**).^26^ Briefly, L- or D-sulfamidates were prepared in two steps from their corresponding *N*-Boc-serine-*O*-tBu starting materials. The PDP scaffolds were generated by the nucleophilic attack of either L- or D-alanine-*O*-tBu to the sulfamidate β-carbon and purified by silica flash chromatography. A final trifluoroacetic acid treatment removed labile *N*-Boc and tert butyl ester protecting groups, affording each of the PDP diastereomers in modest yields after 4 synthetic steps. Derivatization with L-FDAA followed by UPLC-MS analysis enabled the separation of each chemically synthesized diastereomer (**Figure 5C**). Approximately 5-10% serine stereocenter epimerization is observed for all synthetic PDPs, likely occurring during the TFA deprotection. The major in vitro Tln5 product was isolated in a 62% yield from a scaled-up chemoenzymatic reaction. After derivatization with Marfey’s reagent, the Tln5 product aligned with the synthetic D-alanine, L-serine diastereomer (D,L-PDP), supporting its relative stereochemical assignment. With D,L-PDP firmly established as the Tln5 product, we used UPLC-MS to optimize this reaction for buffer, pH, and temperature (**Figures S15–S17**). Tln5 was found to synthesize the most product when 100 mM HEPES, 100 mM KCl, pH 8.0 (**Figure S15**), and room temperature were used for in vitro reaction conditions (**Figure S16**). The enzyme also retained appreciable activity at higher pHs (**Figure S17**) and was tolerant of temperatures up to 37 °C (**Figure S16**). Following the generation of a UPLC-MS based standard curve for Marfey-derivatized D,L-PDP and L,L-PDP products (**Figure S18**), optimized in vitro conditions were used to establish kinetic parameters for the preferred L-OPS and D-alanine substrates (**Figure 5D**). Both Tln5 substrates were fit to Michaelis-Menten kinetics and afforded substrate affinity (*K*_m_) and catalytic rates (*k*_cat_) at values comparable to AMA synthase with its cognate substrates, L-OPS and toxin A (**Figure S19**). Performing analogous experiments with L-alanine as the incoming nucleophile failed to reach enzyme saturation under the conditions tested (**Figure 5D**). Qualitative comparison of the Tln5 rates show a significant preference for D-alanine, which further expands the repertoire of β-substituting PLP-dependent enzymes in Nature.

Given the in vitro preference of Tln5 for D-alanine over its L-enantiomer, we postulated there may be a dedicated alanine racemase encoded in the *tln* biosynthetic gene cluster (**Figure 2**). Genes that annotate as PLP-dependent cystathionine β-lyases have shown in vitro alanine racemase activity,^27,28^ so we set out to explore whether *tln1* encodes a PLP-dependent racemase. An adjacent gene *tln2* that had similarity to PLP-independent epimerase enzymes (**Tables S3, S5**) also sparked our interest, so we created a codon-optimized construct for expression of a *N*-terminally His_6_-tagged Tln2 (hereon Tln2) construct in *E. coli* and purified it with a 25 mg/L yield using Ni-NTA affinity and size exclusion chromatography (**Figure S9**). Overnight incubation of either Tln1 or Tln2 with L-alanine showed a modest accumulation of D-alanine following L-FDAA derivatization and UPLC-MS analyses (**Figure 6A**). The analogous epimerization of D-alanine showed that both enzymes are reversible in vitro. Under equivalent buffer conditions, Tln5 did not appreciably racemize alanine (**Figure S20**) and no enzyme racemized L-OPS in vitro (**Figure S21**). Moreover, Tln1 showed high substrate selectivity for alanine over other amino acid substrates (**Figure S22**). Given the relatively slow epimerization via both PLP-dependent and PLP-independent routes, we wanted to assess which enzyme was capable of faster α-hydrogen abstraction.

**Figure 6.**
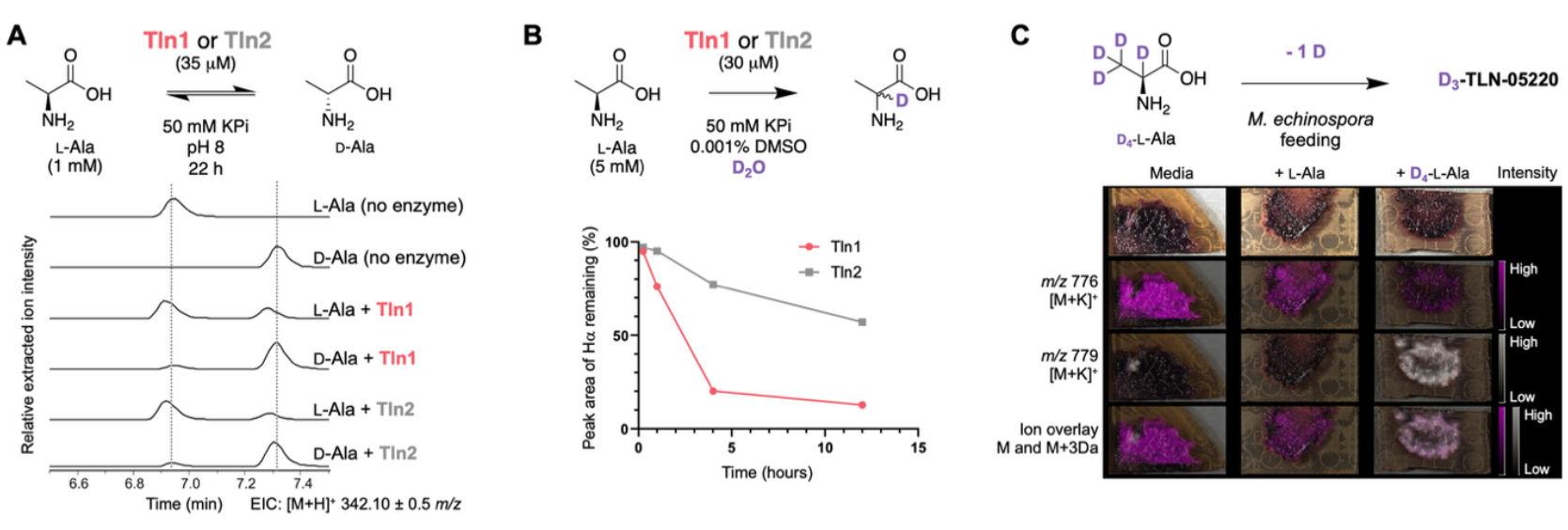
Tln1 and Tln2 catalyze the reversible epimerization of alanine in vitro. (**A**) Tln1 or Tln2 reactivity with either alanine epimer is shown using L-FDAA derivatization and UPLC-MS analyses. No enzyme controls depict where derivatized L- and D-alanine elute (∼6.9 and 7.3 min, respectively). Relative intensities for extracted ion chromatograms in positive mode for Marfey-derivatized alanine ([M+H]^+^ 342.10 ± 0.50 *m/z*) (**B**) ^1^H NMR time course experiments in buffered D_2_O show a time-dependent solvent exchange of the α-hydrogen in L-alanine with both Tln1 (pink) and Tln2 (grey). (**C**) MALDI–IMS with 2,3,3,3-D_4_-L-alanine shows a +3 Da shift when incorporated into TLN-05220.

Adapting previously established protocols,^29^ Tln1 and Tln2 were independently buffer exchanged into D_2_O conditions, and incubated with L-alanine at 0.6 mol%. ^1^H-NMR spectra were taken at various time points, and the integrated alanine α-hydrogen signal (δ 3.75 ppm) was compared to a DMSO internal standard (**Figure S23)**. While both racemases showed a time-dependent decrease of this signal, Tln1 showed a faster overall abstraction (**Figure 6B**). Subsequent UPLC-MS analyses of these D_2_O incubation experiments corroborated that Tln1 incorporated a +1 Da increase in alanine following L-FDAA derivatization (**Figure S24**). To support that alanine racemization is involved in TLN-05220 biosynthesis, we turned to MALDI–IMS to rapidly assess the metabolic profile of *M. echinospora* ATCC 15837 when fed with 2,3,3,3-D_4_-L-alanine in semi-solid agar plates. Intriguingly, we observed a +3 Da shift in TLN-05220 following supplementation of 2,3,3,3-D_4_-L-alanine in comparison to an unlabeled L-alanine control, indicating the loss of one deuterium during biosynthetic incorporation (**Figures 6C, S4, S25**). These complementary in vitro and in vivo results provide further evidence that L-alanine is epimerized prior to incorporation in the final TLN-05220 molecule. To validate that D,L-PDP is involved in TLN-05220 biosynthesis, we chemoenzymatically prepared an isotopically enriched product. Consistent with the mechanism of β-substituting PLP-dependent homologs,^21,22^ Tln5 in buffered D_2_O incorporated a +1 Da shift in the derivatized D,L-PDP product following UPLC-MS analysis (**Figure S24**). Assaying Tln5 in buffered D_2_O with L-OPS and 2,3,3,3-D_4_-DL-alanine generated a pentadeuterated D_5_-PDP product (2-D-L-serine-Cβ-*N*-2,3,3,3-D_4_-DL-alanine), which was fed to *M. echinospora* ATCC 15837 on solid N-Z-Amine agar (**Figure 7**). From these experiments and using measured accurate masses compared to calculated exact masses, we verified that the pseudodipeptide is incorporated into the TLN-05220 product but at a +4 Da shift. Although slightly counter-intuitive, the corresponding feeding experiment with 2,3,3,3-D_4_-DL-alanine generated a +3 Da shifted isotopologue. The overall incorporation was low such that we were not able to obtain enough signal for fragmentation analysis. However, this loss of one deuterium can be biochemically justified by reversible PLP reactivity in vivo by Tln1, Tln5 or endogenous actinobacterial enzymes to ‘wash out’ the α-deuterium on either labeled amino acid substrate prior to installation (**Figure S26**). These cumulative metabolomic and genomic experiments rationalize the transformations of amino acid precursors in the biosynthesis of TLN-05220.

**Figure 7.**
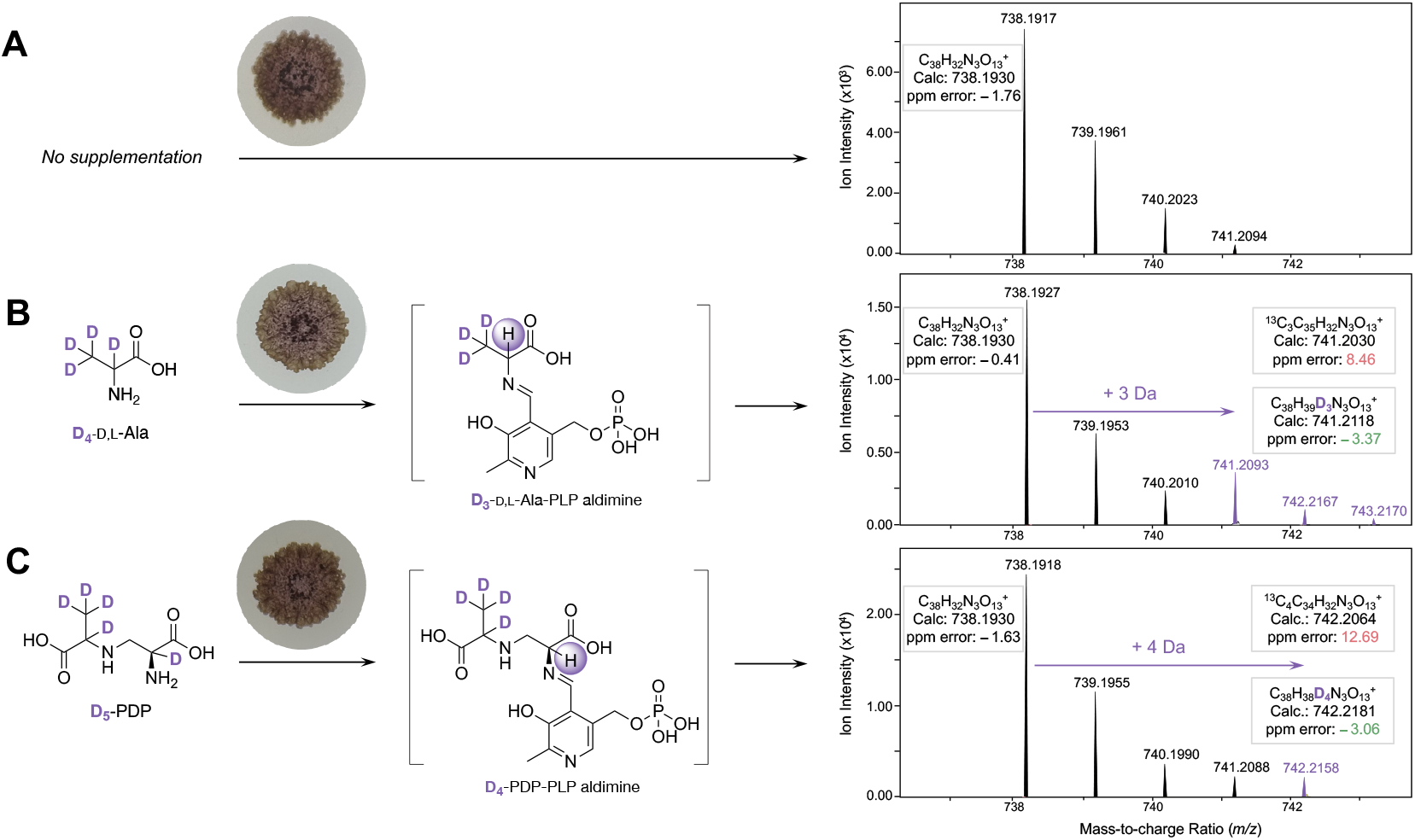
Stable isotope labeling experiments support D,L-PDP as a TLN-05220 biosynthetic intermediate. *M. echinospora* ATCC 15837 was grown on N-Z-Amine agar with (**A**) no amino acid supplementation showing the parent TLN-05220; (**B**) D_4_-D,L-Ala supplementation showing a +3 Da isotopologue shift; and **(C)** D_5_-PDP supplementation showing a +4 Da isotopologue shift. Reversible in vivo epimerization and/or transamination, via a PLP-bound aldimine species, likely removes the α-deuterium adjacent to the primary amine in the supplemented labeled substrates (**Figure S26**). Comparison of the mass defect of D_4_ incorporation supports the successful incorporation of this D_5_-PDP intermediate into TLN-05220, in contrast to the unlikely presence of multiple ^13^C_3_ atoms. A green color denotes formulae within the 5 ppm error threshold, while a red color indicates mass defect values outside of the accepted range.

## Discussion

We were able to verify our biosynthetic predications using a combination of analytical chemistry and chemical biology. From an enzymology perspective, the Tln5 reaction represents a novel addition to the broader biosynthetic β-substituting alanine synthase family of PLP-dependent enzymes. In addition to their established roles in plant biosynthesis and detoxification,^30^ important homologs have been involved in both siderophore and antibiotic biosyntheses in bacterial and fungal pathways (**Figure S27**).^23–25^ SbnA and CmnB both use the α-amine of L-glutamic acid to form the PDP *N*-(2-amino-2-carboxyethyl)glutamic acid (ACEGA) for subsequent oxidative degradation into L-diaminopropionic acid (L-Dap).^23,24^ Similarly, AMA synthase uses the α-amine of L-aspartic acid to form the corresponding PDP known as toxin A; the primary amine of toxin A can also be ligated to a second equivalent of L-OPS to form its namesake AMA natural product.^25^ These enzymes have been explored from a structural and functional perspective and have shown utility as scalable biocatalysts. The β-substitution reaction catalyzed by Tln5 is innovative due to preferring a smaller, hydrophobic and D-enantiomer amino acid. Additional insights into the structural recognition of this preference and its biocatalytic potential are currently ongoing.

We propose that additional amino acid incorporating enzymes in TLN-05220 biosynthesis include two asparagine synthetase homologs, Tln4 and TlnF. Asparagine synthetase homologs are encoded in over a dozen biosynthetic gene clusters that have been tied to production of a pentangular polyphenol (**Figures S3, S4**).^31^ Yet, only two of these enzymes have been experimentally characterized, FdmV and PdmN, each performing different biosynthetic functions. In the fredericamycin A pathway, FdmV donates an amine group from glutamine to a carboxylic acid, acting as an amidotransferase, in the formation of fredericamycin B (**Figure 1 and S28**).^5^ Conversely, PdmN catalyzes a ligation reaction between D-alanine or D-serine and a carboxylic acid of a type II polyketide in pradimicin A biosynthesis (**Figure 1** and **S28**).^6,7^ It has been proposed that the identity of the *N*-terminal amino acid, cysteine in FdmV and serine in PdmN, is a cause of this functional difference.^5^ Amino acid ligation reactions have also been proposed in arixanthomycin^32^ and enduracyclinone^33^ biosynthesis, where the asparagine synthetase homolog has an *N*-terminal serine or alanine, respectively. The correlation between a non-cysteine *N*-terminal residue and ligase activity in asparagine synthetase homologs from the pradimicin, arixanthomycin, and enduracyclinone pathways suggests that Tln4 and TlnF, which share this feature, may likewise catalyze an amino acid ligation reaction in TLN-05220 biosynthesis. (**Figure S5**). Due to the high sequence similarity of TlnF and PdmN, and the position of this gene within the putative TLN-05220 BGC, we predict that TlnF ligates glycine to the polyketide scaffold, most similar to the action of PdmN. Final biosynthetic steps of piperazinone ring formation and ligation to the polyketide-glycine scaffold could also be performed by Tln4, TlnF, and/or other enzymes encoded in the cluster (**Figure S5**) and attempts to characterize their function are underway.

Our work joins other studies that demonstrate stable isotope feeding coupled with MALDI–IMS is a powerful and resource-efficient methodology for studying biosynthesis within biological systems.^34,35^ This combined approach enables the elucidation of biosynthetic pathways and the visualization of the dynamics of metabolite production across entire microbial colonies. Researchers can track the incorporation of stable isotope-labeled precursors (e.g., ^2^H, ^13^C, or ^15^N) into newly synthesized compounds by cultivating microorganisms on media supplemented with minimal amounts of these often-expensive compounds. SIL-IMS is particularly valuable for capturing metabolite intermediates and spatial distributions, even for compounds produced in low concentrations that can still be ionized. Importantly, as new instrumentation becomes readily accessible, MS/MS or collisional cross sections could also be readily gathered.^36^

## Conclusions

Our cumulative efforts identified glycine, serine, and alanine as the three amino acid precursors of TLN-05220 using stable isotopic labeling coupled with LC-MS/MS and MALDI-IMS analyses. Adapting bioinformatic tools and retrobiosynthetic logic, we further expanded the putative BGC beyond its previous annotated boundaries and identified additional amino acid modifying PLP-dependent homologs for subsequent characterization. In vitro enzymology experiments with Tln5 support the condensation of L-OPS with a preferential D-alanine nucleophile and were corroborated with both synthetic chemistry and kinetic analyses. We determined that both Tln1 and Tln2 can act as reversible alanine racemases, yet in a PLP-dependent and cofactor independent manner respectively. Lastly, imaging mass spectrometry and high-resolution mass spectrometry techniques with deuterium-labeled precursors support the in vivo alanine racemase activity and intermediacy of the D,L-PDP Tln5 product in TLN-05220 biosynthesis. Future investigations into the substrate promiscuity of these enzymes will enable the biocatalytic preparation of novel PDPs and β-substituted non-canonical amino acids. Subsequent synthetic biology efforts will enable the mutasynthesis or engineering of novel TLN-05220 analogs following an improved understanding of asparagine synthetase ligation chemistry. These efforts contribute to a greater bioinformatic understanding of amino acid installation into bioactive type II PKS products with potential therapeutic benefit.

## Acknowledgements

This research was generously supported by the William and Linda Frost Fund in the Cal Poly Bailey College of Science and Mathematics, the American Society of Pharmacognosy, the National Science Foundation (CBE 2300890 awarded to K.R.W.; CBE 2300891 awarded to S.M.K.M and L.M.S.; MCB 2128044 awarded to L.M.S.), and the National Institutes of Health (NIH NIGMS R01 GM125943 awarded to L.M.S.; Dissertation year fellowship (G.T.L.).

## References

(1) Wang, J.; Zhang, R.; Chen, X.; Sun, X.; Yan, Y.; Shen, X.; Yuan, Q. Biosynthesis of aromatic polyketides in microorganisms using type II polyketide synthases. Microb. Cell Factories 2020, 19 (1), 110. 10.1186/s12934-020-01367-4.

(2) He, H.; Yang, H. Y.; Luckman, S. W.; Bernan, V. S.; Tsai, G.; Roll, D. M.; Carter, G. T. Echinosporamicin, a New Antibiotic Produced by Micromonospora echinospora ssp. echinospora, LL-P175. Helv. Chim. Acta 2004, 87 (6), 1385–1391. 10.1002/hlca.200490126.

(3) Bristol-Myers Squibb. United States Patent. Antibiotic Bravomicins, 1999.

(4) Banskota, A. H.; Aouidate, M.; Sørensen, D.; Ibrahim, A.; Piraee, M.; Zazopoulos, E.; Alarco, A. M.; Gourdeau, H.; Mellon, C.; Farnet, C. M.; et al. TLN-05220, TLN-05223, new Echinosporamicin-type antibiotics, and proposed revision of the structure of bravomicins. J. Antibiot. (Tokyo) 2009, 62 (10), 565–570. 10.1038/ja.2009.77.

(5) Chen, Y.; Wendt-Pienkowski, E.; Ju, J.; Lin, S.; Rajski, S. R.; Shen, B. Characterization of FdmV as an amide synthetase for fredericamycin A biosynthesis in Streptomyces griseus ATCC 43944. J. Biol. Chem. 2010, 285 (50), 38853–38860. 10.1074/jbc.M110.147744.

(6) Zhan, J.; Qiao, K.; Tang, Y. Investigation of Tailoring Modifications in Pradimicin Biosynthesis. ChemBioChem 2009, 10 (9), 1447–1452. 10.1002/cbic.200900082.

(7) Napan, K.; Zhang, S.; Morgan, W.; Anderson, T.; Takemoto, J. Y.; Zhan, J. Synergistic Actions of Tailoring Enzymes in Pradimicin Biosynthesis. ChemBioChem 2014, 15 (15), 2289–2296. 10.1002/cbic.201402306.

(8) Lautru, S.; Gondry, M.; Genet, R.; Pernodet, J.-L. The Albonoursin Gene Cluster of S. noursei. Chem. Biol. 2002, 9 (12), 1355–1364. 10.1016/S1074-5521(02)00285-5.

(9) Belin, P.; Moutiez, M.; Lautru, S.; Sequin, J.; Pernodel, J.-L.; Gonry, M. The nonribosomal synthesis of diketopiperazines in tRNA-dependent cyclodipeptide synthase pathways. Nat. Prod. Rep. 2012, 29 (9), 961. 10.1039/c2np20010d.

(10) Zheng, L.; Wang, H.; Fan, A.; Li, S.-M. Oxepinamide F biosynthesis involves enzymatic d-aminoacyl epimerization, 3H-oxepin formation, and hydroxylation induced double bond migration. Nat. Commun. 2020, 11 (1), 4914. 10.1038/s41467-020-18713-0.

(11) Guo, Y.; Frisvad, J. C.; Larsen, T. O. Review of Oxepine-Pyrimidinone-Ketopiperazine Type Nonribosomal Peptides. Metabolites 2020, 10 (6), 246. 10.3390/metabo10060246.

(12) Zdouc, M. M.; Blin, K.; Louwen, N. L. L.; Navarro, J.; Loureiro, C.; Bader, C. D.; Bailey, C. B.; Barra, L.; Booth, T. J.; Bozhüyük, K. A. J.; et al. MIBiG 4.0: advancing biosynthetic gene cluster curation through global collaboration. Nucleic Acids Res. 2025, 53 (D1), D678–D690. 10.1093/nar/gkae1115.

(13) Hillenmeyer, M. E.; Vandova, G. A.; Berlew, E. E.; Charkoudian, L. K. Evolution of chemical diversity by coordinated gene swaps in type II polyketide gene clusters. Proc. Natl. Acad. Sci. 2015, 112 (45), 13952–13957. 10.1073/pnas.1511688112.

(14) Chen, S.; Zhang, C.; Zhang, L. Investigation of the Molecular Landscape of Bacterial Aromatic Polyketides by Global Analysis of Type II Polyketide Synthases. Angew. Chem. Int. Ed. 2022, 61 (24), e202202286. 10.1002/anie.202202286.

(15) Blin, K.; Shaw, S.; Augustijn, H. E.; Reitz, Z. L.; Biermann, F.; Alanjary, M.; Fetter, A.; Terlouw, B. R.; Metcalf, W. W.; Helfrich, E. J. N.; et al. antiSMASH 7.0: new and improved predictions for detection, regulation, chemical structures and visualisation. Nucleic Acids Res. 2023, 51 (W1), W46–W50. 10.1093/nar/gkad344.

(16) Blin, K.; Shaw, S.; Vader, L.; Szenei, J.; Reitz, Z. L.; Augustijn, H. E.; Cediel-Becerra, J. D. D.; de Crécy-Lagard, V.; Koetsier, R. A.; Williams, S. E.; et al. antiSMASH 8.0: extended gene cluster detection capabilities and analyses of chemistry, enzymology, and regulation. Nucleic Acids Res. 2025, 53 (W1), W32–W38. 10.1093/nar/gkaf334.

(17) Demarque, D. P.; Crotti, A. E. M.; Vessecchi, R.; Lopes, J. L. C.; Lopes, N. P. Fragmentation reactions using electrospray ionization mass spectrometry: an important tool for the structural elucidation and characterization of synthetic and natural products. Nat. Prod. Rep. 2016, 33 (3), 432–455. 10.1039/C5NP00073D.

(18) Blum, M.; Andreeva, A.; Florentino, L. C.; Chuguransky, S. R.; Grego, T.; Hobbs, E.; Pinto, B. L.; Orr, A.; Paysan-Lafosse, T.; Ponamareva, I.; et al. InterPro: the protein sequence classification resource in 2025. Nucleic Acids Res. 2025, 53 (D1), D444–D456. 10.1093/nar/gkae1082.

(19) Banerjee, R.; Evande, R.; Kabil, Ö.; Ojha, S.; Taoka, S. Reaction mechanism and regulation of cystathionine β-synthase. Biochim Biophys Acta Proteins Proteom 2003, 1647 (1), 30–35. 10.1016/S1570-9639(03)00044-X.

(20) Buller, A. R.; Van Roye, P.; Murciano-Calles, J.; Arnold, F. H. Tryptophan Synthase Uses an Atypical Mechanism To Achieve Substrate Specificity. Biochemistry 2016, 55 (51), 7043–7046. 10.1021/acs.biochem.6b01127.

(21) Du, Y.-L.; Ryan, K. S. Pyridoxal phosphate-dependent reactions in the biosynthesis of natural products. Nat. Prod. Rep. 2019, 36 (3), 430–457. 10.1039/C8NP00049B.

(22) Mizutani, T.; Abe, I. Pyridoxal 5′-Phosphate (PLP)-Dependent β- and γ-Substitution Reactions Forming Nonproteinogenic Amino Acids in Natural Product Biosynthesis. J. Nat. Prod. 2025, 88 (1), 211–230. 10.1021/acs.jnatprod.4c01226.

(23) Kobylarz, M. J.; Grigg, J. C.; Liu, Y.; Lee, M. S. F.; Heinrichs, D. E.; Murphy, M. E. P. Deciphering the Substrate Specificity of SbnA, the Enzyme Catalyzing the First Step in Staphyloferrin B Biosynthesis. Biochemistry 2016, 55 (6), 927–939. 10.1021/acs.biochem.5b01045.

(24) Hsu, S.-H.; Zhang, S.; Huang, S.-C.; Wu, T.-K.; Xu, Z.; Chang, C.-Y. Characterization of Enzymes Catalyzing the Formation of the Nonproteinogenic Amino Acid l-Dap in Capreomycin Biosynthesis. Biochemistry 2021, 60 (1), 77–84. 10.1021/acs.biochem.0c00808.

(25) Guo, Q.; Wu, D.; Gao, L.; Bai, Y.; Liu, Y.; Guo, N.; Du, X.; Yang, J.; Wang, X.; Lei, X. Identification of the AMA Synthase from the Aspergillomarasmine A Biosynthesis and Evaluation of Its Biocatalytic Potential. ACS Catal. 2020, 10 (11), 6291–6298. 10.1021/acscatal.0c01187.

(26) Albu, S. A.; Koteva, K.; King, A. M.; Al-Karmi, S.; Wright, G. D.; Capretta, A. Total Synthesis of Aspergillomarasmine A and Related Compounds: A Sulfamidate Approach Enables Exploration of Structure–Activity Relationships. Angew. Chem. Int. Ed. 2016, 55 (42), 13259–13262. 10.1002/anie.201606657.

(27) Soo, V. W. C.; Yosaatmadja, Y.; Squire, C. J.; Patrick, W. M. Mechanistic and Evolutionary Insights from the Reciprocal Promiscuity of Two Pyridoxal Phosphate-dependent Enzymes. J. Biol. Chem. 2016, 291 (38), 19873–19887. 10.1074/jbc.M116.739557.

(28) Miyamoto, T.; Katane, M.; Saitoh, Y.; Sekine, M.; Homma, H. Cystathionine β-lyase is involved in damino acid metabolism. Biochem. J. 2018, 475 (8), 1397–1410. 10.1042/BCJ20180039.

(29) Cordoza, J. L.; Chen, P. Y.-T.; Blaustein, L. R.; Lima, S. T.; Fiore, M. F.; Chekan, J. R.; Moore, B. S.; McKinnie, S. M. K. Mechanistic and Structural Insights into a Divergent PLP-Dependent l-Enduracididine Cyclase from a Toxic Cyanobacterium. ACS Catal. 2023, 13 (14), 9817–9828. 10.1021/acscatal.3c01294.

(30) Watanabe, M.; Kusano, M.; Oikawa, A.; Fukushima, A.; Noji, M.; Saito, K. Physiological Roles of the β-Substituted Alanine Synthase Gene Family in Arabidopsis. Plant Physiol. 2008, 146 (1), 310–320. 10.1104/pp.107.106831.

(31) Lackner, G.; Schenk, A.; Xu, Z.; Reinhardt, K.; Yunt, Z. S.; Piel, J.; Hertweck, C. Biosynthesis of Pentangular Polyphenols: Deductions from the Benastatin and Griseorhodin Pathways. J. Am. Chem. Soc. 2007, 129 (30), 9306–9312. 10.1021/ja0718624.

(32) Kang, H. S.; Brady, S. F. Arixanthomycins A-C: Phylogeny-guided discovery of biologically active eDNA-derived pentangular polyphenols. ACS Chem. Biol. 2014, 9 (6), 1267–1272. 10.1021/cb500141b.

(33) Monciardini, P.; Bernasconi, A.; Iorio, M.; Brunati, C.; Sosio, M.; Campochiaro, L.; Landini, P.; Maffioli, S. I.; Donadio, S. Antibacterial Aromatic Polyketides Incorporating the Unusual Amino Acid Enduracididine. J. Nat. Prod. 2019, 82 (1), 35–44. 10.1021/acs.jnatprod.8b00354.

(34) Eckelmann, D.; Kusari, S.; Spiteller, M. Stable Isotope Labeling of Prodiginines and Serratamolides Produced by Serratia marcescens Directly on Agar and Simultaneous Visualization by Matrix-Assisted Laser Desorption/Ionization Imaging High-Resolution Mass Spectrometry. Anal. Chem. 2018, 90 (22), 13167–13172. 10.1021/acs.analchem.8b03633.

(35) Tat, V. T.; Lee, Y. J. Spatio-Temporal Study of Galactolipid Biosynthesis in Duckweed Using Mass Spectrometry Imaging and in vivo Isotope Labeling. Plant Cell Physiol. 2024, 65 (6), 986–998. 10.1093/pcp/pcae032.

(36) Shepherd, R. A.; Luu, G. T.; Sanchez, L. M. MALDI-TIMS-MS^2^ Imaging and Annotation of Natural Products in Fungal-Bacterial Coculture. Anal. Chem. 2025, 97 (35), 18867–18872. 10.1021/acs.analchem.5c02787.

