## Supporting Information for "Integrated metabolomic and genomic insights into amino acid incorporation within the hybrid polyketide-alkaloid antibiotic TLN-05220"

**This PDF file includes:**

General Materials and Methods
Figures S1–S28
Tables S1–S5
Chemical Synthesis
Compound Characterization
Supplementary Information References

Supplemental Information: **Integrated metabolomic and genomic insights into amino acid incorporation within the hybrid polyketide-alkaloid antibiotic TLN-05220**

**<u>Table of Contents</u>**


General Methods and Materials…… S4
Molecular Biology and Biochemical Methods…… S7
Enzyme Assay Methods…… S9
Supplementary Figures…… S12
- Figure S1…… S12
- Figure S2…… S13
- Figure S3…… S14
- Figure S4…… S15
- Figure S5…… S16
- Figure S6…… S18
- Figure S7…… S19
- Figure S8…… S20
- Figure S9…… S21
- Figure S10…… S22
- Figure S11…… S23
- Figure S12…… S24
- Figure S13…… S25
- Figure S14…… S26
- Figure S15…… S28
- Figure S16…… S28
- Figure S17…… S29
- Figure S18…… S30
- Figure S19…… S31
- Figure S20…… S32
- Figure S21…… S33
- Figure S22…… S34
- Figure S23…… S35
- Figure S24…… S36
- Figure S25…… S37
- Figure S26…… S38
- Figure S27…… S39
- Figure S28…… S40

Supplementary Tables…… S41
- Table S1…… S42
- Table S2…… S43
- Table S3…… S44
- Table S4…… S45
- Table S5…… S46

Synthetic Chemistry Protocols…… S47
- Synthesis of *O*-succinyl-L-serine SI-2…… S47
- Synthesis of D,L-PDP Tln5 diastereomer SI-6…… S48
- Synthesis of L,L-PDP Tln5 diastereomer SI-9…… S49
- Synthesis of D,D-PDP Tln5 diastereomer SI-13…… S50
- Synthesis of L,D-PDP Tln5 diastereomer SI-15…… S52

NMR and Compound Characterization…… S53
- SI-1…… S53
- SI-2…… S54
- SI-3…… S57
- SI-5…… S60
- SI-6…… S63

SI-8........................................................................................ S66
SI-9........................................................................................ S69
SI-10...................................................................................... S72
SI-11...................................................................................... S75
SI-12...................................................................................... S80
SI-13...................................................................................... S83
SI-14...................................................................................... S84
SI-15...................................................................................... S87
Enzymatic SI-6.......................................................................... S90
SI-16...................................................................................... S93
SI-17...................................................................................... S93
Supplemental References .......................................................... S94

Supplemental Information: **Integrated metabolomic and genomic insights into amino acid incorporation within the hybrid polyketide-alkaloid antibiotic TLN-05220**

### General Methods and Materials

#### Bacterial strains and reagents

*Micromonospora echinospora* ATCC 15837 was purchased from the American Type Culture Collection (ATCC). Isotopically labelled amino acids, 1-$^{13}$C-L-alanine (99%), 2,3,3,3-$D_4$-L-alanine, 2,3-$^{13}C_2$-L-alanine, 1,2,3-$^{13}C_3$-L-serine, 1,2-$^{13}C_2$-glycine and NMR solvents ($CDCl_3$, $CD_3OD$, and $D_2O$ with 0.1% $CH_3OH$) were purchased from Cambridge Isotope Laboratories. The following chemicals and solvents were purchased through Fisher Scientific: ethylenediaminetetraacetic acid (EDTA), nickel chloride, N-Z-Amine Type A, soluble starch, yeast extract, dextrose, imidazole, isopropyl *β*-D-1-thiogalactopyranoside (IPTG), L-alanine, glycine, L-serine, dichloromethane (DCM), methanol (MeOH), and formic acid (96+%).

#### Synthesis and analytical methods

For chemical synthesis, solvents were purchased from MilliporeSigma and Thermo Fisher Scientific, and reagents were used without further purification unless otherwise stated. Reactions were monitored using precoated thin-layer chromatography (TLC) plates (Merck, TLC silica gel 60 $F_{254}$) and visualized with potassium permanganate staining solution (0.75 g $KMnO_4$, 5 g $K_2CO_3$, 0.625 mL 10% NaOH in 100 mL of water). TLC $R_f$ values were calculated and rounded to the nearest 0.05. Silica gel (Alfa Aesar, 60 (215–400 mesh)) was used to purify products with flash column chromatography, and rotary evaporation was used to concentrate samples under reduced pressure. Nuclear magnetic resonance (NMR) spectra were obtained using an Avance III HD spectrometer (Bruker) equipped with a BBFO SmartProbe at 500 MHz ($^1$H NMR) or 125 MHz ($^{13}$C NMR) in $CDCl_3$, $CD_3OD$, or $D_2O$ + 0.1% $CH_3OH$ solvents. Chemical shifts for synthetic reactions are reported in ppm (δ) and references the $CDCl_3$ solvent signal (δ = 7.26 ppm for $^1$H, δ = 77.2 ppm for $^{13}$C NMR), $CD_3OD$ solvent signal (δ = 3.34 ppm for $^1$H, δ = 49.86 ppm for $^{13}$C), or methanol as an internal standard for $D_2O$ (δ = 3.34 ppm). NMR data are reported as: s = singlet, d = doublet, t = triplet, q = quartet, m = multiplet, *J* = coupling constant (Hz). High-resolution mass spectrometry (HRMS) data were obtained on a timsTOFfleX (Bruker) matrix-assisted laser desorption/ionization-quadruple time of flight (MALDI-QqTOF) mass spectrometer in positive ESI mode.

#### Stable isotope feeding on solid agar

*Micromonospora echinospora* ATCC 15837 (DSM 43816) was grown in liquid N-Z-Amine media (10 g dextrose, 10 g soluble starch, 5 g yeast extract, 5 g N-Z-Amine Type A, 1 g calcium carbonate, 1000 mL deionized water, pH 7.0–7.4) or on plates with variable agar concentrations, as noted. For all feeding studies, an ATCC 15837 spore stock was inoculated into N-Z-Amine media (1:250 v/v) with 100 µg/mL kanamycin for selection. Note that ATCC 15837 is a gentamycin producer and therefore has intrinsic aminoglycoside resistance.[1] Cultures were grown for 10 days at 30°C with 220 rpm, where the culture appeared a deep purple.

#### Sample growth with $^{13}$C-labeled amino acids and preparation for Liquid Chromatography (LC)-HRMS analysis

A series of 5 mL N-Z-Amine agar (1.5%) plates with 50 µg/mL kanamycin were prepared with additional supplementation of the following sterile amino acid solutions to a final concentration amino acid amount of 1.05 mg/mL: 2,3-$^{13}C_2$-L-alanine, L-alanine, 1,2,3-$^{13}C_3$-L-serine, L-serine, both 2,3-$^{13}C_2$-L-alanine and 1,2-$^{13}C_2$-glycine (0.525 mg/mL each), and both L-alanine and glycine (0.525 mg/mL each). Five plates of each type were inoculated with 100 µL of the 10-day ATCC 15837 culture on the center of the plate and left open to dry in a laminar flow hood for 10–20 min. Plates were incubated at 30 °C for 5 days.

After 5 days of growth, the center of the plate that contained bacterial growth was excised, cut into smaller pieces, and transferred to a 15 mL conical tube. The agar/bacteria sample was flash frozen in liquid nitrogen and lyophilized overnight. The dried sample was transferred to a 20 mL scintillation vial crushed and ground to a powder. 5 mL of 9:1 dichloromethane: methanol was added to each sample, followed by 5 mL of 0.1% aqueous formic acid (96+%, Alfa Aesar). Samples were mixed by inversion and allowed to incubate in the solvents for extraction for 3–14 h. The organic layer was removed using a Pasteur pipette, and the aqueous layer was extracted again with 10 mL of 9:1 dichloromethane: methanol. The extraction sat at room temperature for an additional 0.5–3 h. The organic layers of each sample were combined and concentrated in vacuo. Samples were stored or shipped at -20 °C for subsequent analyses.

<u>Sample growth with $D_5$-PDP (**SI-16**) and preparation for Liquid Chromatography (LC)-HRMS analysis</u>

A series of 5 mL N-Z-Amine agar (1.5% final) plates with 50 µg/mL kanamycin were prepared with the supplementation of either 3 mM $H_5$-PDP (**SI-17**) or 3 mM $D_5$-PDP (**SI-16**). $H_5$-PDP and $D_5$-PDP compound stocks were each prepared at 30 mM in water and sterile filtered using a 0.22 µm PES syringe filter (Sterlitech Corporation) prior to addition. Four plates of each type were inoculated with 100 µL of a 10-day ATCC 15837 culture on the center of the plate and left open to dry in a laminar flow hood for 10–20 min. Plates were incubated at 30 °C for 5 days. After 5 days of growth, the center of the plate that contained bacterial growth was excised, cut into small pieces using a sterile inoculation loop, and transferred to a 50 mL conical tube. The agar/bacteria samples were flash frozen in liquid nitrogen and lyophilized overnight. The dried samples were each transferred into a 20 mL scintillation vial for small scale extractions. 5 mL of 9:1 dichloromethane:methanol was added to each sample, followed by 5 mL of 0.1% aqueous formic acid. Samples were mixed by inversion and incubated at room temperature for 3–5 h. The organic layer was removed using a Pasteur pipette, and the aqueous layer was extracted once more with 10 mL of 9:1 dichloromethane:methanol and was incubated at room temperature for an additional 0.5–2 h. The organic layers of each sample were combined and concentrated in vacuo. Samples were stored and shipped at −20 °C for subsequent LC-HRMS analyses.

<u>(LC)-HRMS analysis</u>

Sample extracts were normalized to 1 mg/mL in methanol.

For samples containing $^{13}C$-labeled amino acids, LC-HRMS data were acquired on a Bruker timsTOF fleX qTOF in positive ESI mode with trapped ion mobility spectrometry disabled. Data were acquired in Hystar 6.2 and otofControl 6.2.201 with a detection window of 100–2000 *m/z*. Samples were separated on an Agilent Poroshell UPLC column (1.9 µm C18, 50 x 2.1 mm) with a flow rate of 0.5 mL/min the following gradient with MilliQ water with 0.1% formic acid (solvent A) and acetonitrile with 0.1% formic acid (solvent B): initial 2 min hold at 5% solvent B, 5–100% solvent B over 10 min, 2 min hold at 100% solvent B, 1 min re-equilibration at 5% solvent B. ESI-L Low Concentration Calibration Standard Tuning Mix (G1969-85000, Agilent Technologies Inc, Santa Clara, CA) was injected via divert valve from 0.04 to 0.30 min during the LC data acquisition for post analysis mass recalibration. For each sample, 2 µL of a 1 mg/mL solution was injected. The ESI conditions were set with the capillary voltage at 4.5 kV. For MS/MS, dynamic exclusion and the top 9 precursor ions from each $MS^1$ scan were subject to collision energies scaled according to the mass and charge state for a total of 9 data dependent MS/MS events per $MS^1$ via Auto MS/MS mode. Isolation widths and collision energies are interpolated from the table below. Values lower than the minimum (i.e. 50) or higher than the maximum (i.e. 1300) mass use the fixed values from the table.

| *Mass* | *Isolation Width* | *Collision Energy* | *Charge State* | *Priority Number* |
|---|---|---|---|---|
| *50* | 2 | 20 | 1 | 1 |
| *1000* | 6 | 20 | 1 | 2 |
| *500* | 4 | 20 | 1 | 3 |
| *1300* | 8 | 30 | 1 | 4 |

For samples containing $D_5$-PDP (**SI-16**), liquid chromatography and MS/MS analysis were carried out on a Bruker TimsTOF fleX coupled with an Elute ultra-performance liquid chromatography (UPLC) system equipped with a separation Kinetex column (1.7 µm C18, 50 x 2.1 mm). Solvents used were A: Milli-Q water with 0.1% FA and B: ACN with 0.1% FA. The flow rate was 0.5 mL/min, and the separation method was from 0 to 6 mins linear gradient to 95% B, from 6 to 12 mins linear gradient to 50% B, and from 12 to 15 mins isocratic. The mass spectrometer was operated in positive mode in the scan range of 50 – 1500 m/z and at a spectra rate of 6 Hz, choosing the top 9 most intense peaks. The instrument was calibrated using ESI-L low concentration Tuning mix (Agilent). The method was calibrated using the enhanced quadratic algorithm with zooming set to +/- 0.01%; the calibration was only accepted when a 100% match was achieved which is equivalent to a standard deviation of <= 0.50 ppm. CID (Collision Induced Dissociation) was the fragmentation method used to collect MS/MS sequentially. The parameters set for the collision cell were 7.0 eV collision energy.

Sample growth and preparation for MALDI analysis

A series of 5 mL N-Z-Amine agar (1.5% w/v) plates with 2.5 µg/mL rifampicin were prepared with additional supplementation of the following sterile amino acid solutions to a final concentration of 1.05 mg/mL: 2,3,3,3-$D_4$-L-alanine, and L-alanine. Three plates of each type were inoculated with 50 µL of the 10-day ATCC 15837 culture on the center of the plate and left open to dry in a laminar flow hood for 10–20 min. Plates were incubated at 30 °C for 16 days.

MALDI-IMS analysis

MALDI-TOF IMS was carried out on a Bruker Autoflex Speed LRF mass spectrometer. For each experiment, media controls were run alongside ATCC 15837 grown on NZ-amine media, on amino acid supplemented media, or on SIL amino acid supplemented media for comparison of production (ionization) under each growth condition. These colonies were wet mounted onto a stainless steel 96 spot MALDI target plate, coated with 50:50 α-cyano-hydroxycinnamic acid (CHCA) and dihydroxybenzoic acid (DHB) matrix using a 53-micron strain steel sieve (Hogentogler & Co Inc.) and dried in a desiccator for 4 h at 37 °C and analyzed by MALDI-TOF IMS. Pictures of the sample were taken wet, after matrix application and after desiccation for teach points in the software. Samples were analyzed by MALDI-TOF IMS at 80% laser power, 10.5 X gain 500 µm raster in positive ion reflectron mode. Data was normalized using root mean square (RMS) as indicated using FlexImaging v. 5.0.

Dried droplet spectra were also acquired of the SIL amino acids to confirm the measured accurate masses using 50:50 CHCA:DHB matrix in positive ion reflectron mode on a Bruker Autoflex Speed LRF mass spectrometer.

Ultra-performance liquid chromatography-mass spectrometry (UPLC-MS) methods

UPLC-MS data were obtained on an Elute UHPLC system (Bruker) coupled to an ESI-ion trap IT (amaZon SL, Bruker) mass spectrometer in positive ion mode using the eluents of 0.1% aqueous formic acid (Solvent A) and 0.1% formic acid in acetonitrile (Solvent B). Method A was used for general assay analysis, and Method B was used for diastereomeric separation of PDP compounds.

Method A is as follows: 10% to 20% B over 3 minutes, 20% to 45% B over 3 minutes, 45% - 100% B over 2 minutes, hold at 100% B for 2 minutes, 100% to 10% B over 1 minute, hold at 10% B for 4 minutes.

Method B is as follows: 10% B hold for 1 minute, 10% to 20% B over 5 minutes, 20% to 45% B over 3 minutes, 45% to 100% B for 2 minutes, 100% B hold for 2 minutes, 100% B to 10% B over 2 minutes, then 10% hold for 2 minutes.

General Marfey's derivatization for UPLC-MS analysis

The protocol for Marfey's derivatization of enzymatic assays were followed from previously used methods. Saturated aqueous sodium bicarbonate (20 μL) was added to a 1 mM enzyme assay solution (50 μL), followed by 100 μL of 1% w/v solution of 1-fluoro-2,4-dinitrophenyl-5-L-alanimide (L-FDAA, Marfey's reagent) prepared in acetone. The derivatization reaction was incubated at 37 °C for 90 min, then neutralized with 1 N HCl (25 μL), and centrifuged at 14,000 × *g* for 10 min. The subsequent supernatant was extracted and analyzed via UPLC-MS.

Reagents for $D_2O$ assays

A premade solution of 500 mM buffered $K_2HPO_4/KH_2PO_4$ buffer (KPi, pH 8) in $H_2O$ was lyophilized for 16 h, resuspended in $D_2O$, then lyophilized again. This sample was resuspended in $D_2O$, then used as buffer for preparing enzyme assays and preparing the storage solution for purified enzymes. Stock solutions of PLP, L-OPS, L- or D-alanine, and any additional substrates or cofactors were prepared in $D_2O$. All solutions in $D_2O$ were stored under Ar gas at -20 °C.

### Molecular Biology and Biochemical Methods

Plasmid transformation and purification

*E. coli* codon-optimized pET28α(+) plasmids were designed, then purchased from Twist Biosciences with a *N*-terminal hexahistidine tag for *tln1*, *tln2*, and *tln5*. Plasmids were transformed into chemically competent *E. coli* DH10β or BL21(DE3) using the same protocol as previously done.

Plasmid purifications were performed with the QIAprep Spin Miniprep Kit (QIAGEN) or the GeneJET Plasmid Miniprep Kit (Thermo Scientific) following the manufacturer's protocol. Warm, sterile MilliQ water or elution buffer (10 mM Tris-HCl, pH 8.5) were used for plasmid elution from the column, and plasmid concentration and purity were measured via NanoDrop.

Tln protein expression

pET28a-*tln1*, -*tln2*, and -*tln5* plasmids were transformed into *E. coli* BL21(DE3) cells and selected for on an LB-agar plate containing 50 μg/mL kanamycin. Starter cultures were made by selecting single colonies and inoculating into 15 mL LB supplemented with 50 μg/mL kanamycin, which was incubated overnight at 37 °C at 200 rpm for 12–18 h. 10 mL of the starter culture was inoculated into 1 L of Terrific Broth media containing 50 μg/mL kanamycin, which was incubated at 37 °C, 200 rpm. Cultures containing pET28a-*tln1* or -*tln2* were brought to an $OD_{600}$ of 0.6–0.8, whereas pET28a-*tln5* cultures were brought to an $OD_{600}$ of 1.0–1.2. Subsequently, cultures were cooled to 18 °C at 200 rpm for 1 h. IPTG (100 μM, final concentration) and PLP (50 mg for *tln1* and *tln5* cultures) were added to induce protein expression and promote cofactor incorporation and cultures were incubated for 12–18 h at 18 °C and 200 rpm. Cultures were harvested by centrifugation at 2500 × *g* at 10 °C for 30 min and cells were resuspended in 30 mL of cell lysis buffer (1 M NaCl, 20 mM Tris-HCl pH 8.0, 20 mM imidazole, 10% glycerol) and stored at -70 °C until protein purification.

Tln protein purification

*E. coli* BL21(DE3) cell pellets containing *N*-terminal hexahistidine-tagged Tln1, Tln2 or Tln5 were thawed and then sonicated on ice at 40% amplitude with a cycle of 15 s on, and 45 s off for a total of 6 min (FisherBrand Model 505 Sonic Dismembrator, 3.2 mm microtip). The lysate was clarified by centrifugation at 15000 × *g* for 30 min at 12 °C. Proteins were purified at 4 °C using an AKTAGo FPLC system with a HisTrap FF (5 mL) column (Cytiva Life Sciences) that was pre-equilibrated with 30 mL Buffer A (20 mM Tris-HCl pH 8.0, 30 mM imidazole, 1 M NaCl, 100 μM PLP) at a flow rate of 2 mL/min. Buffers used for FPLC purification were filtered through a 0.22 mM nitrocellulose membrane prior to use. After loading the clarified cell lysate onto the HisTrap FF column, the column was washed with Buffer A until the UV absorbance returned to baseline, and then with 10% Buffer B (20 mM Tris-HCl pH 8.0, 1 M NaCl, 250 mM imidazole, 100 μM PLP) to remove non-specifically bound proteins until the UV absorbance returned to baseline (maximum 25 mL). Hexahistidine-tagged proteins were eluted using a linear gradient of 10–100% B over 60 mL (12 column volumes) collecting 5 mL fractions. The eluted fractions were assessed for purity using a 10% SDS-PAGE gel, and fractions with >90% purity for Tln1, Tln2, or Tln5 (39, 28, and 38 kDa, respectively) were combined and concentrated to 2.5 mL using an Amicon ultra centrifugal filter (10 kDa molecular weight cut-off (MWCO), EMD-Millipore, Inc.). Concentrated proteins were buffer exchanged into a protein storage buffer (50 mM HEPES pH 8.0, 300 mM KCl) using a preequilibrated PD-10 column (Cytiva Life Sciences). Protein stock concentrations were quantified using a Bradford assay with a bovine serum albumin standard and, if necessary, proteins were further concentrated with the Amicon ultra centrifugal filter (10 kDa MWCO). After obtaining purified protein, the protein was aliquoted into 100 µL aliquots and stored at −70 °C. Harvested *E. coli* cultures of Tln1 yielded 8.9 mg/L, Tln2 yielded 25.0 mg/L, and Tln5 yielded 80.2 mg/L.

For Tln5 steady-state kinetics assays, Tln5 was further purified on a Superdex 75 HiLoad 16/600 size exclusion column (Cytiva Life Sciences) equilibrated with storage buffer. Tln5 (1000 µL) was injected onto the column and eluted with storage buffer at a rate of 1 mL/min. Tln5 purity was assessed using a 10% SDS-PAGE gel and pure fractions were concentrated using an Amicon ultra centrifugal filter (10 kDa MWCO, EMD-Millipore, Inc.). Protein concentrations were assessed using Bradford assay and immediately used for kinetics assays. Any Tln5 that remained post-assay was stored at −70 °C.

Tln5 buffer exchange for $^{1}$H NMR assays

For solvent suppressed $^{1}$H NMR assays, a 1 mL aliquot of purified Tln5 enzyme was thawed on ice, then buffer exchanged into buffered 50 mM $K_2HPO_4/KH_2PO_4$ (KPi) aqueous solution at pH 8.0. The Tln5 aliquot was pipetted in a 10 MWCO Amicon ultra centrifugal filter followed by the addition of 5 mL KPi, and the sample was centrifuged for 15 min at 4 °C and 3400 × *g*. An additional 5 mL of KPi was added and centrifuged and this process was repeated a total of 3 times. The concentration of the resulting sample was assessed by Bradford, then the buffer exchanged Tln5 was directly used in the $^{1}$H NMR assay and any remaining Tln5 was stored at -70 °C until further use.

Tln1, Tln2, and Tln5 buffer exchange for $D_2O$ assays

For deuterium incorporation assays, from the stock 500 mM KPi (pD 8.0) in $D_2O$, a solution of 50 mM KPi (pD 8.0) was prepared for buffer exchanging purified enzymes into deuterated conditions. A 1 mL aliquot of purified enzyme was thawed on ice, then pipetted into an Amicon ultra centrifugal filter (10 kDa) along with 5 mL of 50 mM KPi (pD 8.0) in $D_2O$, and centrifuged for 15 min at 4 °C and 3400 × *g*. An additional 5 mL of 50 mM KPi (pH 8.0) in $D_2O$ was added to the filter, centrifuged, and this process was repeated a total of three times. The concentration of the deuterated buffer enzyme was assessed by Bradford assay and enzymes were directly used for in vitro assays. Any remaining enzyme was stored at -70 °C until further use.

### Enzyme Assay Methods

#### Tln1 in vitro functional assays

Characterization assays of Tln1 were carried out in 50 mM KPi (pH 8.0), using 1 mM of substrate (L- or D-alanine), 100 μM PLP, and 50 μM Tln1. The total reaction volume was brought up to 50 μL using ultrapure water and incubated at room temperature overnight (16 h). After incubation, assays were derivatized with Marfey's reagent using the above protocol then analyzed via UPLC-MS.

#### Tln2 in vitro functional assays

Characterization assays of Tln2 were carried out in 50 mM KPi (pH 8.0) using 1 mM of substrate (unless otherwise stated) and 50 μM Tln2. The total reaction volume was brought up to 50 μL using ultrapure water and incubated at room temperature overnight (16 h). After incubation, assays were derivatized with Marfey's reagent using the above protocol prior to UPLC-MS analysis.

#### Tln5 in vitro functional assays

Characterization assays of Tln5 were carried out in 50 mM KPi (pH 8.0), using 1 mM of each substrate (unless otherwise stated), 100 μM PLP, and 50 μM Tln5. The total reaction volume was brought up to 50 μL using MilliQ water and incubated at room temperature for 1 h. After incubation, assays were derivatized with Marfey's reagent using the above protocol prior to UPLC-MS analysis.

#### Tln1 and Tln5 UV-Vis spectroscopy

Assays were analyzed in a 500 μL, 10 mm quartz cuvette cell (VWR International, LLC) with an Eppendorf BioSpectrometer between 200 and 800 nm. Enzymatic assays were prepared in 50 mM KPi (pH 8.0), with 50 μM Tln1 or Tln5, in a final volume of 500 μL.

#### Tln5 standard curve preparation

Standard curves of D,L- and L,L-PDP were generated for Tln5 steady-state kinetics assays. Synthetic standards of D,L- (SI-6) and L,L-PDP (SI-9) were prepared in triplicate as 50 μL aliquots at concentrations ranging from 25 to 1000 μM in 100 mM HEPES (pH 8.0), 100 mM KCl, and 100 μM PLP. Prior to derivatization, 2.75 μL of 10 mM L-glutamic acid was added with 2.25 μL of ultrapure water to maintain L-glutamic acid as an internal standard at 500 μM (total sample volume of 55 μL). Standards were derivatized with Marfey's reagent using the above protocol prior to UPLC-MS analysis.

#### Tln5 kinetics assay

Tln5 steady-state kinetics assays were performed in triplicate with 1500 μM OPS, 5 μM SEC-purified Tln5 protein, 100 mM HEPES (pH 8.0), 100 mM KCl, and 25-2500 μM D- or L-alanine. Assays were incubated at 23 °C (room temperature) for 15 min, quenched with 25 μL of a Marfey's derivatization master mix (20 μL sat. aq. $NaHCO_3$, 2.25 μL ultrapure water, and 2.75 μL 10 mM L-glutamic acid for each assay), then derivatized with Marfey's reagent using the protocol described above, then subjected to UPLC-MS analysis using Method A. For analysis, product peak areas were converted to rates of conversion using D,L- and L,L-PDP standard curves, which were plotted using Michaelis-Menten kinetic parameters to calculate $k_{cat}$, $K_m$, and $k_{cat}/K_m$ using GraphPad Prism.

For obtaining kinetic parameters of L-OPS, a similar protocol was followed, however the concentration of D-alanine was held constant at 1500 μM, and the concentration of L-OPS varied from 25–2500 μM.

Tln5 chemoenzymatic assay and product purification
Enzymatic production of PDP (chemoenzymatic SI-16) was achieved through an in vitro enzyme assay in 50 mM KPi (pH 8) with 25 mM L-OPS, 25 mM D-alanine, 100 μM PLP, and 30 μM Tln5. The reaction incubated at room temperature overnight (15 h), subsequently quenched with 500 μL of acetonitrile, vigorously vortexed, and centrifuged (14000 × *g*, 10 min). The supernatant was collected and concentrated in vacuo; dried supernatant was redissolved in water to a 15 mg/mL concentration to purify via HPLC. Samples were run on a Synergi 4 μm Polar-RP 80 Å LC column (250 x 10 mm, Manufacturer) at a flow rate of 2.5 mL/min using the following method: 3 min 10% B, 7 min gradient from 10–50% B, 3 min gradient from 50–100% B, 2 min 100% B, 2 min 100–10% B, 3 min 10% B with the desired product eluting at 4.7 min.. The final product was lyophilized overnight to yield the product **SI-16** as a white solid. $^1H$ NMR (500 MHz, in $D_2O$ + 0.1% $CH_3OH$): δ 4.10 (dd, *J* = 8.9, 5.4 Hz, 1H), 3.85 (q, *J* = 7.2 Hz, 1H), 3.61–3.47 (m, 2H), 1.57 (d, *J* = 7.2 Hz, 3H). $^{13}C$ NMR (126 MHz, $D_2O$ + 0.1% $CH_3OH$): δ 174.29, 169.75, 58.44, 49.58, 45.03, 14.87. HRMS (MALDI) Calculated for $C_6H_{13}N_2O_4$ 177.0870, observed 177.0887 $[M+H]^+$.

$D_2O$ in vitro assay protocols
Tln1 and Tln5 deuterium incorporation assays were conducted in a similar manner to a literature reference.[2] Tln1 and Tln5 were each buffer exchanged into 50 mM $K_2HPO_4$ (pD 8.0) in $D_2O$ using separate 30 kDa MWCO Amicon centrifugal filter. After thawing protein aliquots on ice, the protein was transferred to the centrifugal filter with 50 mM $K_2HPO_4$ (pD 8.0, 5 mL), then centrifuged at 3400 × *g* for 20 min. Buffer exchange continued with two additional washes of $K_2HPO_4$ in $D_2O$ (5 mL). The concentration of the final buffer-exchanged protein stock was determined using a Bradford assay. Fresh stock solutions of L-OPS, L- and D-alanine, and PLP were prepared in $D_2O$, and stock $K_2HPO_4$ was lyophilized, refrozen in 5 mL of $D_2O$, and lyophilized again (process repeated for 2 $D_2O$ washes). To keep concentrations consistent, the buffer stock was resuspended in the original volume instead of $D_2O$.

Deuterium incorporation assays were set up in >98% deuterium conditions. All stock solutions were prepared in $D_2O$, and purified proteins were buffer exchanged into 50 mM KPi using the protocol above. Assays were set up in 50 mM KPi (pH 8.0) with 1 mM substrate, 100 µM PLP, and 50 µM purified enzyme, and reaction volumes were brought to 50 µL with $D_2O$. Assays were incubated at room temperature overnight and derivatized with Marfey's reagent.

Chemoenzymatic $D_5$-PDP (**SI-16**) production and purification
A scaled up reaction for the 2-D-L-serine-Cβ-*N*-2,3,3,3-$D_4$-DL-alanine (abbreviated to $D_5$-PDP herein) product for feeding studies was assembled in $D_2O$ with 50 mM KPi (pD 8.0), 100 μM PLP, 25 mM L-OPS, 25 mM 1:1 D:L-$D_4$-alanine, and 30 μM purified Tln5 protein. The reaction was brought up to the final volume of 12 mL with $D_2O$, then stirred at room temperature (23 °C) for 18 h. Afterward, the reaction was quenched with 1 CV $CH_3OH$ (12 mL), then centrifuged (5 min, 3500 x *g*) at 10 °C to promote protein precipitation and clarify the assay supernatant. The supernatant was decanted, concentrated in vacuo, then purified on a silica flash column using a 3:1:1.5 *n*-butanol: acetic acid: water eluent system. The desired product was isolated as a mixture of alanine diastereomers as an acetate salt that was a white powder (39.0 mg, 36% yield). $^1H$ NMR (500

MHz, $D_2O$ + 0.1% $CH_3OH$) δ 4.15 (dh, *J* = 11.8, 4.0, 3.3 Hz, 1H), 4.10 – 4.01 (m, 1H). HRMS (MALDI) Calculated for $C_6H_8D_5N_2O_4$ 182.1184, found 182.1198 $[M+H]^+$.

Chemoenzymatic $H_5$-PDP (**SI-17**) production and purification

As a control for the feeding studies that $d_5$-PDP was used for, a protiated version of the molecule, $H_5$-PDP was also conducted. The same protocol for the $D_5$-PDP was used for producing the $h_5$-PDP, with the exception of the reaction using 1:1 D:L-alanine, and the reaction proceeding in $H_2O$ instead of $D_2O$. This reaction was assembled in $H_2O$ with 50 mM KPi (pH 8.0), 100 μM PLP, 25 mM L-OPS, 25 mM 1:1 D:L-alanine, and 30 μM purified Tln5 protein. The reaction was brought up to the final volume (12 mL) using $H_2O$, then stirred at room temperature (23 °C) for 18 h. Afterward, the reaction was quenched with 1 CV $CH_3OH$ (12 mL) then centrifuged (5 min, 3500 x *g*) at 10 °C to promote protein precipitation and clarify the reaction supernatant. The supernatant was decanted, concentrated in vacuo, then purified on a silica flash column using a 3:1:1.5 *n*-butanol: acetic acid: water eluent system. The desired product was isolated as a mixture of alanine diastereomers with trace alanine contamination as an acetate salt that was a white powder (30.5 mg, 31% yield). $^1H$ NMR (500 MHz, $D_2O$ + 0.1% $CH_3OH$) δ 4.06 – 3.97 (m, 1H), 3.79 – 3.70 (m, 1H), 3.53 – 3.37 (m, 2H), 3.00 – 2.85 (m, 1H), 1.49 (d, *J* = 6.9 Hz, 3H), 1.44 (d, *J* = 7.2 Hz, 1H). HRMS (MALDI) Calculated for $C_6H_{13}N_2O_4$ 177.0870, found 177.0880 $[M+H]^+$.

### Supplementary Figures

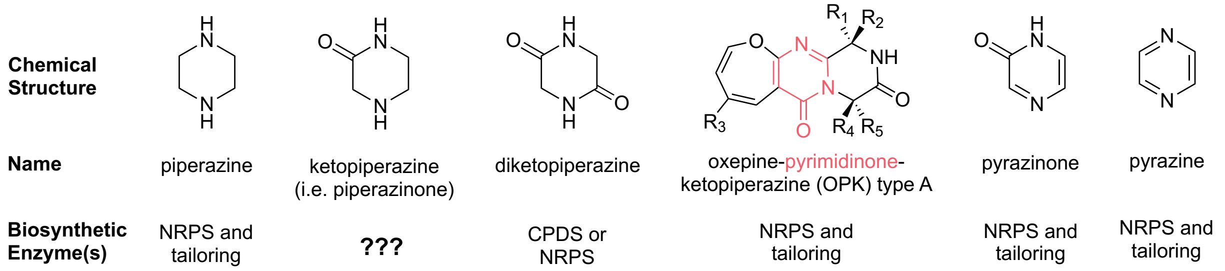


**Figure S1.** Chemical structures of the related piperazine, ketopiperazine (i.e. piperazinone), diketopiperazine, pyrimidinone, pyrazinone and pyrazine moieties. The pyrimidinone group within oxepine-pyrimidinone-ketopiperazine (OPK) structure is highlighted in pink.[3] Enzymes like cyclic dipeptide synthetases (CDPSs), non-ribosomal peptide synthetases (NRPSs), often with oxidative or reductive tailoring enzymes or domains, are known to be involved in the biosynthesis of these structural classes.[3–8] The piperazinone biosynthetic enzymes are unknown and the focus of this work.

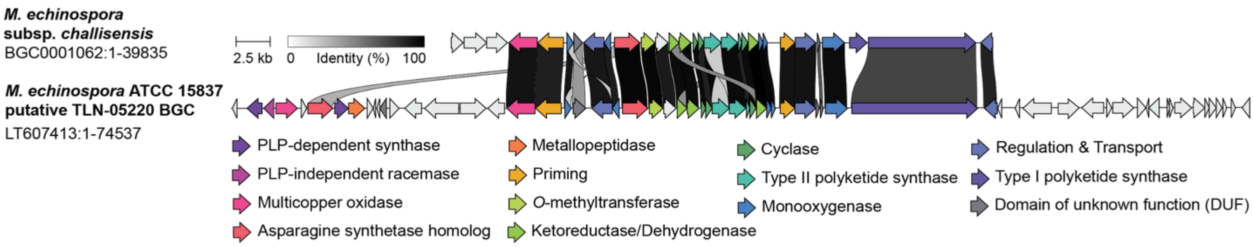


**Figure S2.** Alignment of candidate TLN-05220 biosynthetic gene cluster (BGC) from ATCC 15837. (Top) BGC0001062 available in Minimal Information about a Biosynthetic Gene Cluster (MIBiG) repository,[9] the TLN-05220 BGC previously reported in *Micromonospora echinospora* subsp. *challisensis* by Banskota et al.[10] (Bottom) Putative TLN-05220 BGC from *M. echinospora* ATCC 15837 identified by antiSMASH.[11] Ribbon diagram created with clinker on CAGECAT with a 30% minimum sequence identity alignment.[12] The percent identity of the aligned genes is denoted by the grayscale color of the connecting ribbons.

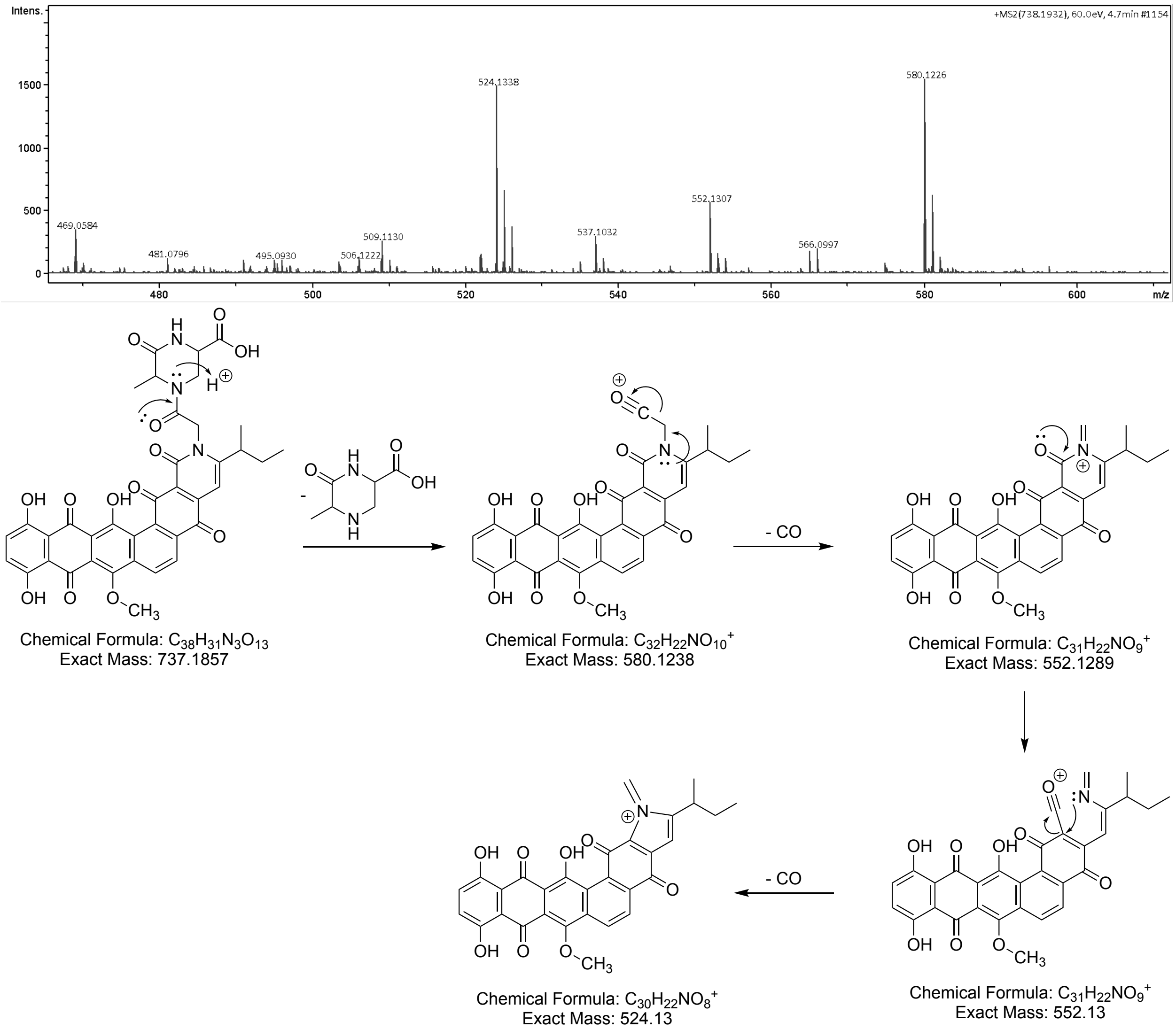


**Figure S3.** TLN-05220 fragmentation pattern by MS/MS. TLN-05220 $[M+H]^+$ measured accurate mass 738.1913; calc. exact mass 738.1930.

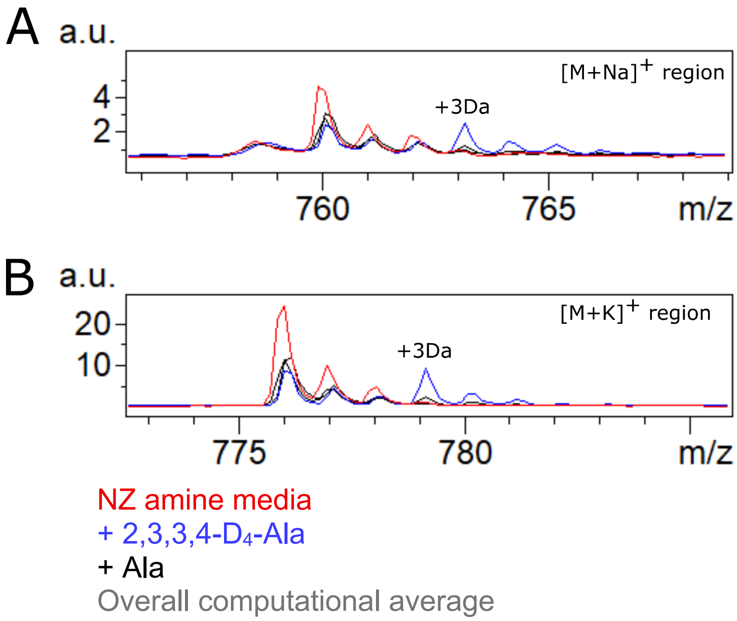


**Figure S4.** Detection of TLN-05220 on semi-solid agar plates using MALDI-IMS. A. Zoom in of the sodiated adduct, production was observed in all conditions at *m/z* 760.1 with $D_4$-Ala added to the media a peak is observed at *m/z* 763.1. B. Zoom in of the potassiated adduct, production was observed in all conditions at *m/z* 776.1 with 2,3,3,3-$D_4$-Ala added to the media a peak is observed at *m/z* 779.1. These spectra are shown from specific regions of interest and compared to the overall computational averaged spectrum displayed in grey using FlexImaging v. 5.0.

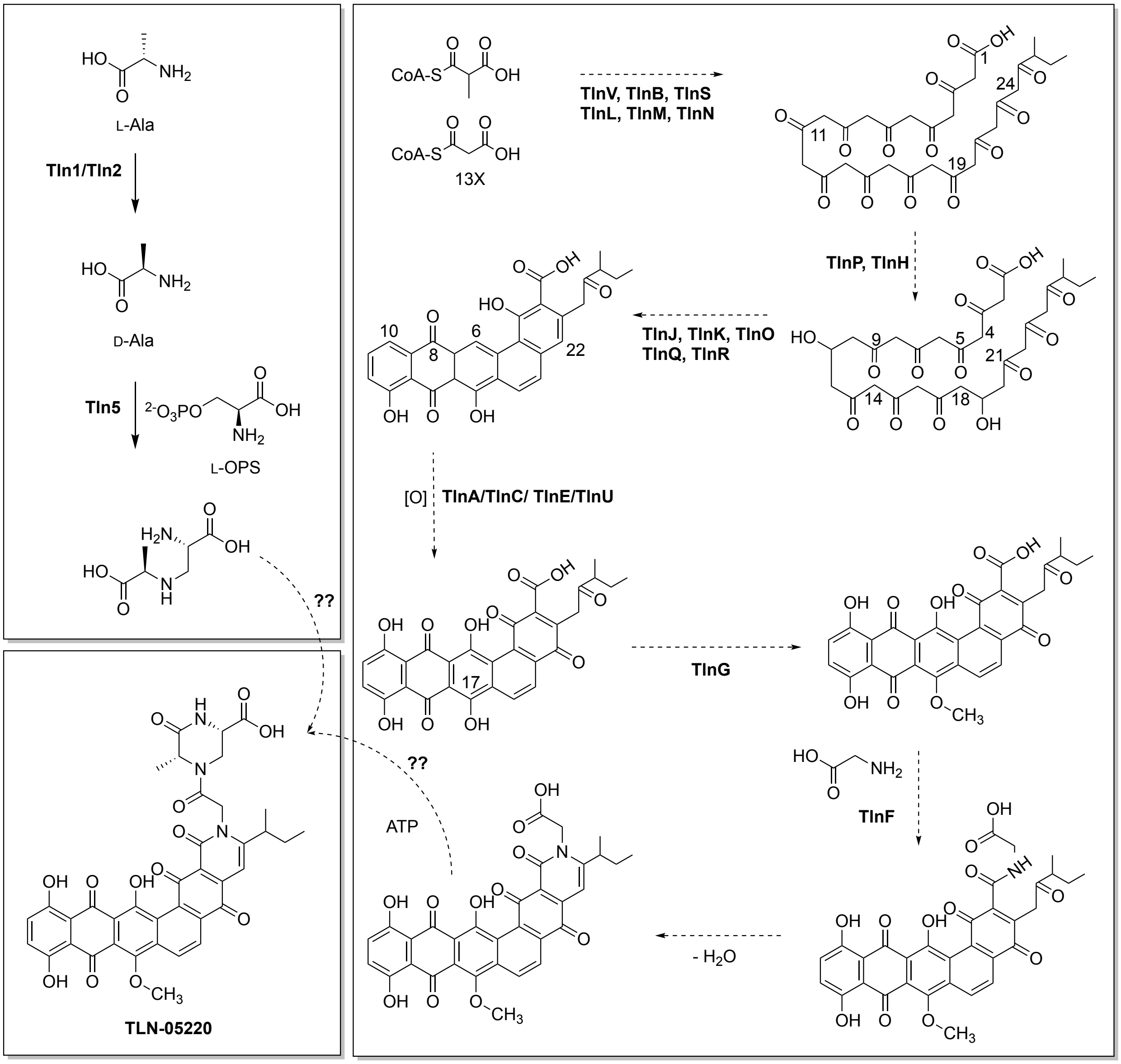


**Figure S5.** Biosynthetic hypothesis for TLN-05220 production in *M. echinospora* ATCC 15837. Analyses with PRISM 4.0[13] and antiSMASH 8.0[11] identify TlnV as a modular type I PKS composed of a loading module with a decarboxylating ketosynthase ($KS_Q$), followed by a single reducing extension module. Both acyltransferase (AT) domains carry the canonical YASH motif associated with methylmalonyl-CoA specificity.[14] Notably, YASH-containing ATs downstream of $KS_Q$ in loading modules have been shown to accept malonyl-CoA in vitro in some systems.[15] Depending on the substrate preference of the TlnV loading AT ($AT_L$), TlnV is poised to generate a 2-methylbutyryl starter unit for the downstream type II PKS. Transfer of the 2-methylbutyryl group from the terminal ACP of TlnV to the type II ACP TlnN likely involves TlnS, a KSIII, together with the acyl-transferase–like protein TlnB. KSIII/AT pairs are implicated in the assembly of non-acetate primers in several type II polyketides (hedamycin, daunomycin, aclacinomycin, R1128, frenolicin, oxytetracycline),[16] and among these only the hedamycin cluster encodes a type I PKS. Consistent with this, antiSMASH's type II PKS starter-unit prediction favors 2-methylbutyrate as a substrate for both TlnS and TlnB.[17] By analogy to HedS, KSIII TlnS may impose starter-unit specificity, selecting 2-methylbutyryl over an acetate generated by simple malonyl decarboxylation. As in HedS, DpsC and AknE2, the catalytic cysteine of TlnS is replaced by serine, a hallmark of KSIIIs engaged in starter selection.[16] The TLN-05220 cluster lacks an auxiliary ACP outside the minimal type II PKS genes, supporting a model in which TlnS mediates direct transfer from the TlnV ACP. The AT-like TlnB may assist in loading this starter onto the minimal PKS, similar to the proposed role of HedF in hedamycin biosynthesis.[18] TlnB also shows close similarity to San2 from A-74528 biosynthesis, an enzyme proposed to act as an acyl-ACP thioesterase that could purge the 2-methyl butyrate starter unit from the type I PKS.[19,20] An alternative, not mutually exclusive, possibility is that TlnV primes the minimal type II PKS directly, as observed in reconstituted hedamycin systems.[21] Ongoing experiments in our laboratory are aimed at resolving this type I/type II handoff. The minimal type II PKS, composed of ketosynthase α TlnL, ketosynthase β (chain length fator) TlnM, and acyl carrier protein TlnN is predicted to extend the 2-methylbutyryl starter with twelve malonyl-CoA units to afford a C29 polyketide chain. Formation of the pentangular polyphenolic scaffold requires two programmed ketoreductions. Reduction at C19 is known to enforce the pentangular geometry in related pathways.[22] antiSMASH annotates TlnP as a C19 ketoreductase (KR), and phylogenetic analysis places TlnP with characterized C19 KRs, whereas TlnH clusters with XanZ4[23] and other enzymes assigned as C11 KRs, in agreement with its antiSMASH annotation. Cyclization/aromatization of the nascent polyketide is likely catalyzed by a set of Tcm/Pdm-like cyclases and monooxygenases TlnJ, TlnK, TlnO, TlnQ, TlnR.[24–26] TlnO is TcmN-like and is proposed to install the first two rings with bond formations C9–C14 and C7–C16. Subsequent ring closures at C5–C18, C4–C2, and C2–C23 are plausibly catalyzed by TlnJ (TcmI-like) and TlnK (TcmJ-like, cupin domain). TlnQ and TlnR resemble PdmH- and PdmI-family monooxygenases, respectively. As in pradimicin biosynthesis, monooxygenases likely act cooperatively with cyclases to drive aromatization of two of the newly closed rings and to introduce oxygen at C8.[24,27] Subsequent tailoring requires installation of three oxygens; conversion of the anthrone to a quinone via oxidation at C22, and hydroxylations at C6 and C10. Four cluster enzymes could mediate these steps. TlnA is a predicted multicopper oxidase, TlnC and TlnE are annotated as anthrone-type oxygenases, and TlnU is an FAD-dependent monooxygenase. Phylogenetically, TlnC and TlnE are closely related to FdmM1 (65% similarity) and FdmM (60% similarity), enzymes that hydroxylate C6 and C8 in fredericamycin biosynthesis.[28] By analogy, TlnC and TlnE are candidates for C6 and C10 hydroxylation in TLN-05220. Oxidation at C22 is expected to precede quinone formation. TlnU clusters with polyketide monooxygenases such as PnxO4 from FD-594 biosynthesis[29] (56% similarity), and could contribute to this step, though roles for TlnC and TlnE cannot be excluded. TlnA shares 58% similarity with a multicopper oxidase (ORF16) from the diazopinomicin pathway.[30] If functionally analogous, TlnA could oxidize a hydroquinone intermediate to the C22 quinone. *O*-methylation at C17 is most reasonably assigned to TlnG, an *O*-methyltransferase with high similarity to PdmT (pradimicin, 68%), XanM3 (xantholipin, 61%), and Hex9 (hexaricin, 60%), and is expected to use *S*-adenosyl-L-methionine as the methyl donor. Finally, based on its strong homology to the amino-acid ligase PdmN, TlnF likely ligates glycine to the polyketide core.[31] In parallel, Tln1, Tln2, and Tln5 assemble a pseudodipeptide from L-alanine and L-*O*-phosphoserine, as reported herein. The closing steps that generate the piperazinone and couple it to the polyketide-glycine intermediate may be executed by Tln4, TlnF, and/or additional cluster enzymes; efforts to define these activities are in progress.

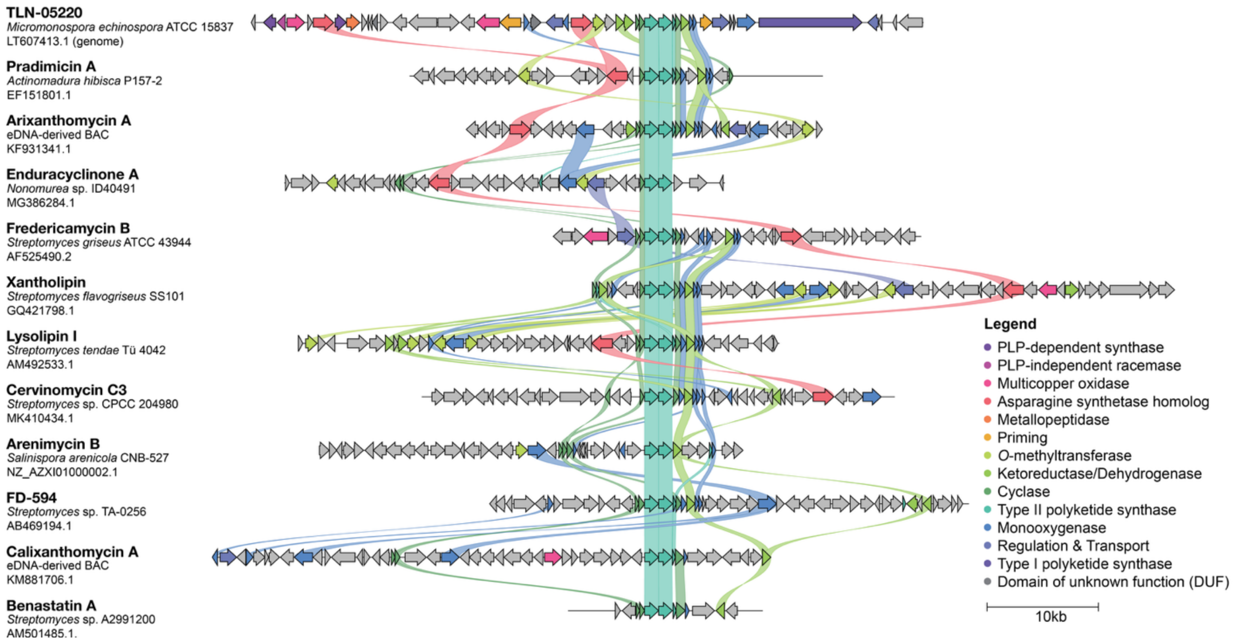


**Figure S6.** Alignment of TLN-05220 biosynthetic gene cluster with gene clusters encoding structurally related pentangular polyphenols (also see **Figure S4**).[29,32–42] The TLN-05220 cluster has conserved components with known pentangular polyphenol biosynthetic gene clusters. Ribbon diagram of the TLN-05220 cluster from *Micromonospora echinospora* ATCC 15837 was created using clinker on CAGECAT with a 40% similarity score cut-off.[8] Ribbons are colored by sequence similarity group and drawn at 30% identity threshold. The ribbon diagram is centered on highly conserved KS-CLF components of the type II PKS system (teal). NCBI Genbank accession numbers listed beneath structure and producing organism name.

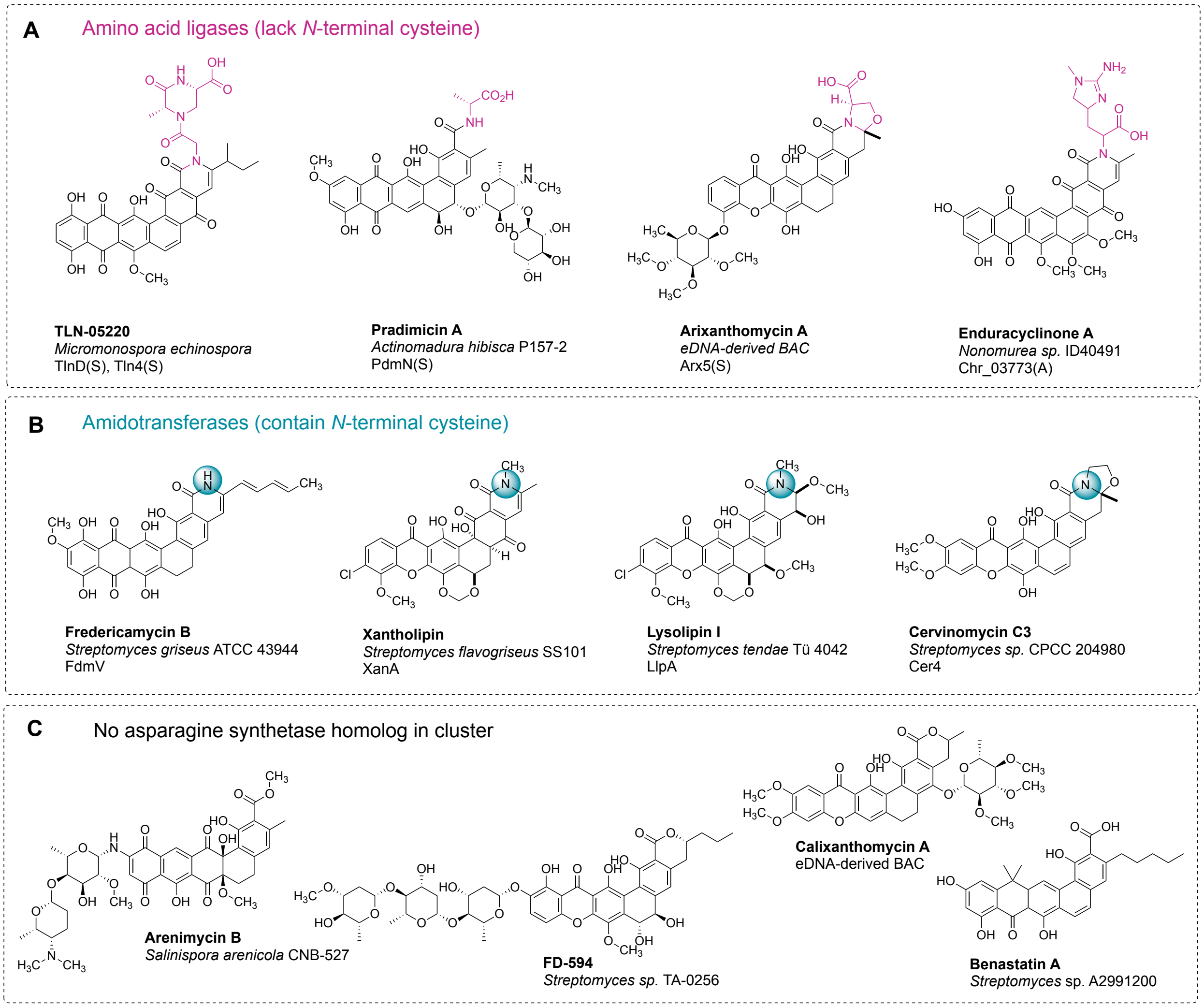


**Figure S7.** Molecular structures of TLN-05220 and related pentangular polyphenols, divided into groups by presence and type of asparagine synthetase homolog in biosynthetic gene cluster (BGC, see also **Figure S3**).[29,32–42] All structures contain a pentacyclic polyphenol core, and structure names and producing organisms are listed under each structure. (A) Structures correspond to a BGC containing one or more asparagine synthetase homologs that lack an *N*-terminal cysteine and are putative amino acid ligases. Structural regions likely derived from amino acids are colored in pink. Amino acid ligase protein name listed beneath producing organism, and single-letter code for *N*-terminal residue. (B) Structures correspond to a BGC containing one asparagine synthetase homolog with an *N*-terminal cysteine and are putative amidotransferases. Nitrogen atoms likely derived from an amino acid are highlighted with a teal circle. Amidotransferase protein name listed beneath producing organism. (C) Structures correspond to a BGC lacking an asparagine synthetase homolog.

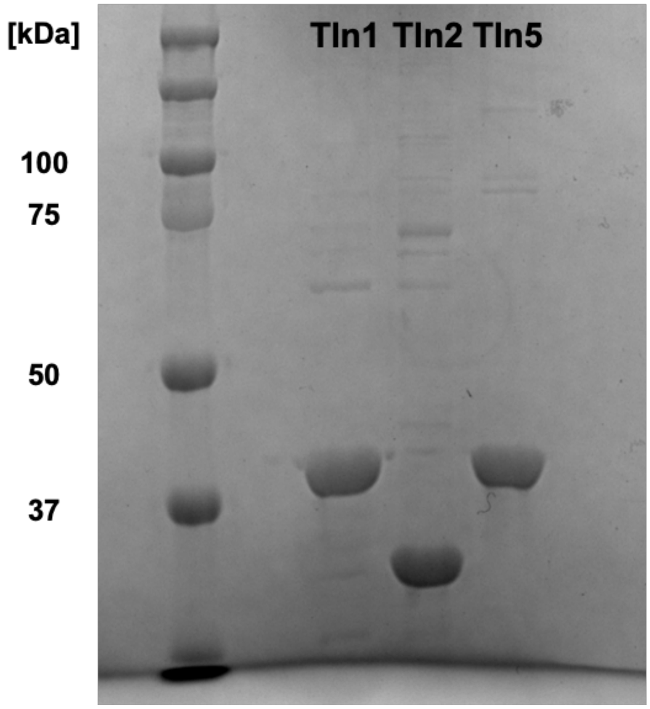


**Figure S8.** SDS-PAGE of Tln1, Tln2, and Tln5 hexahistadine-tagged proteins. A 4–10% acrylamide protein gel was prepared and loaded with Precision Plus Protein™ Dual Color Standards (Bio-Rad) and each respective Tln protein (10 µg). The expected molecular weights for Tln1, Tln2 and Tln5 are: 40, 27, and 39 kDa, respectively.

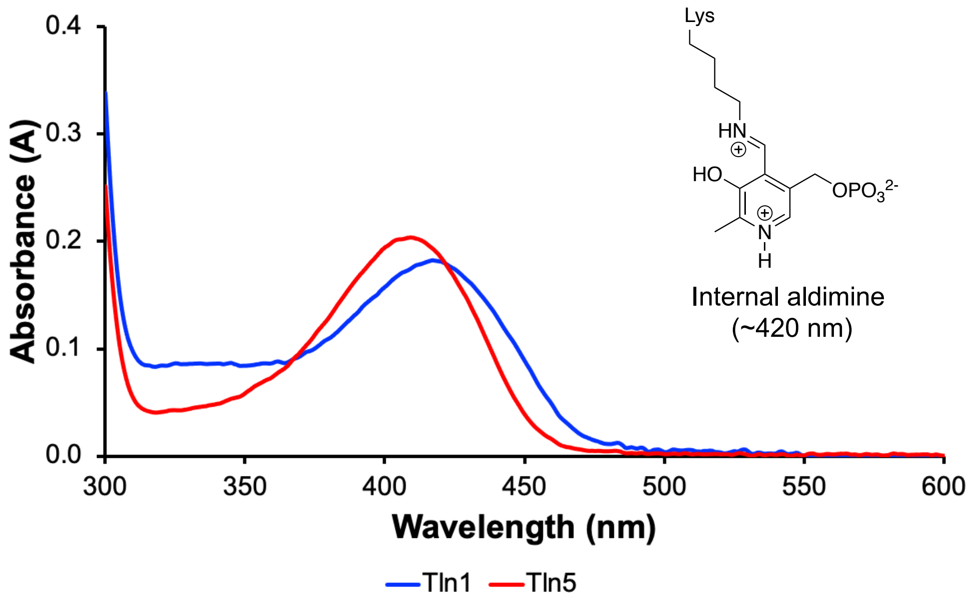


**Figure S9.** UV-Vis spectral data of Tln1 (blue) and Tln5 (red) reveal bound internal aldimine state, indicative of bound PLP cofactor. Spectral readings were taken with 50 μM of protein in 50 mM KPi (pH 8.0). Tln1 and Tln5 have a peak at 420 nm, which is indicative of the active-site-bound PLP as an internal aldimine.

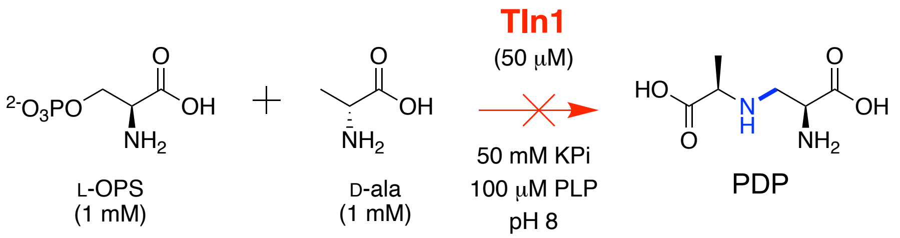


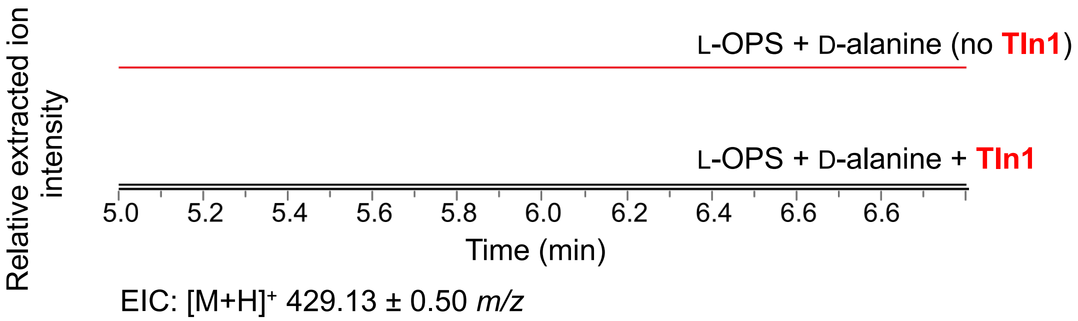


**Figure S10.** Tln1 does not catalyze a β-substitution reaction. Tln1 assays were assembled using '<u>Tln1 in vitro functional assays'</u> protocols described above, then incubated at room temperature for 16 h. Reactions were derivatized with Marfey's reagent for optimized retention time prior to UPLC-MS analysis. Relative intensities for extracted ion chromatograms in positive mode were extracted from UPLC-MS traces (EIC ± 0.50 *m/z*) for Marfey-derivatized PDP ($[M+H]^+$ 429.13 *m/z*) to reveal that no PDP was produced in the presence of Tln1 (black trace) when compared to the trace omitting Tln1 (red trace).

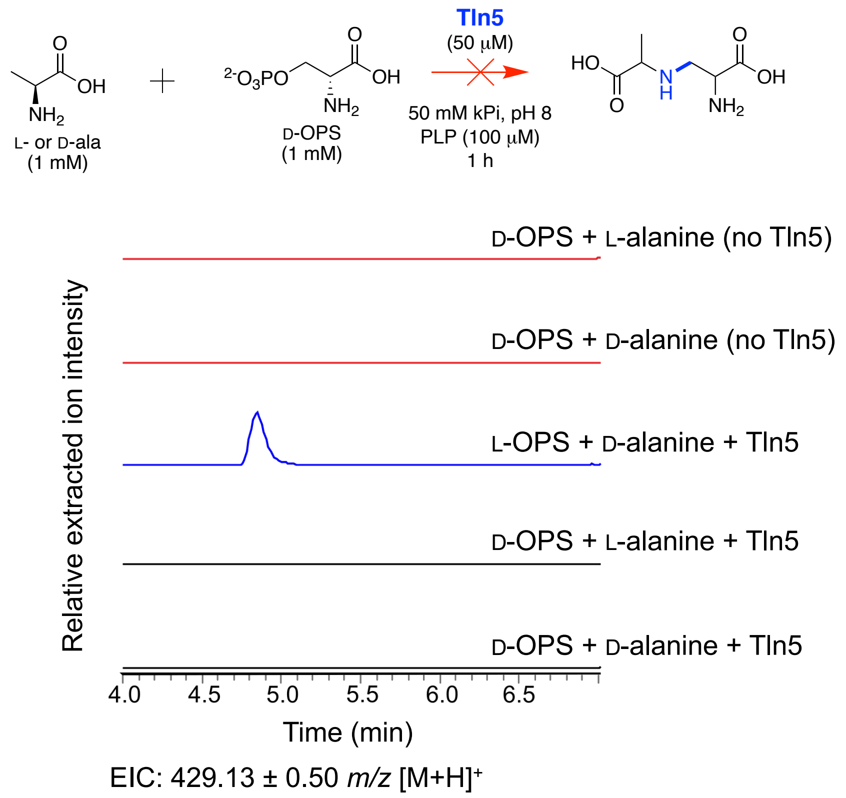


**Figure S11.** Tln5 does not react with D-OPS. Tln5 reactions and controls were set up with D-OPS and either D- or L-alanine and incubated at room temperature for 1 h. Samples were then derivatized with Marfey's reagent prior to UPLC-MS analysis. Relative intensities of extracted ion chromatograms in positive mode were extracted from UPLC-MS traces (EIC ± 0.50 *m/z*) for Marfey-derivatized PDP ($[M+H]^+$ 429.13 *m/z*). Compared to a positive control using L-OPS (blue) as the serine donor, no PDP is produced in the presence of D-OPS.

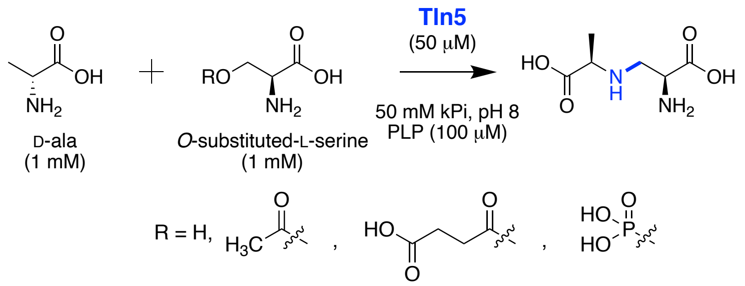


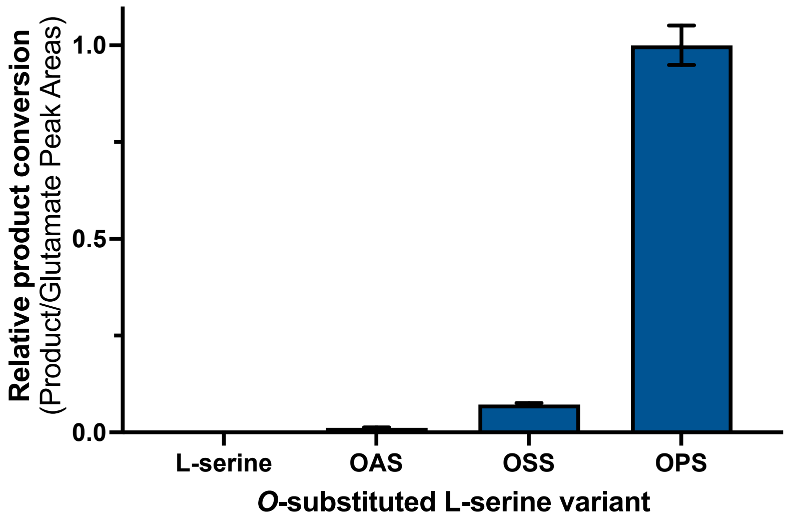


**Figure S12.** Relative quantitation of Tln5 activity reveals *O*-phospho-L-serine (L-OPS) as the preferred serine substrate. Assays were set up set up in biological triplicate according to the 'Tln5 in vitro functional assays' protocol above and incubated at room temperature for 1 h. An internal standard of 500 mM L-glutamic acid was added to the assay at the time of Marfey derivatization, then assays were subjected to UPLC-MS analysis. Relative intensities of positive mode extracted ion chromatograms were extracted from UPLC-MS traces (EIC ± 0.50 *m/z*) for both PDP and L-glutamic acid ($[M+H]^+$ 429.13, 400.11 *m/z*), then the peak area of PDP was normalized to the internal standard (L-glutamic acid). Here, the averages of the normalized peak areas are depicted with the standard deviation as error bars. The data exhibit that L-OPS is the preferred serine substrate, but Tln5 can still use L-OSS and L-OAS as substrates.

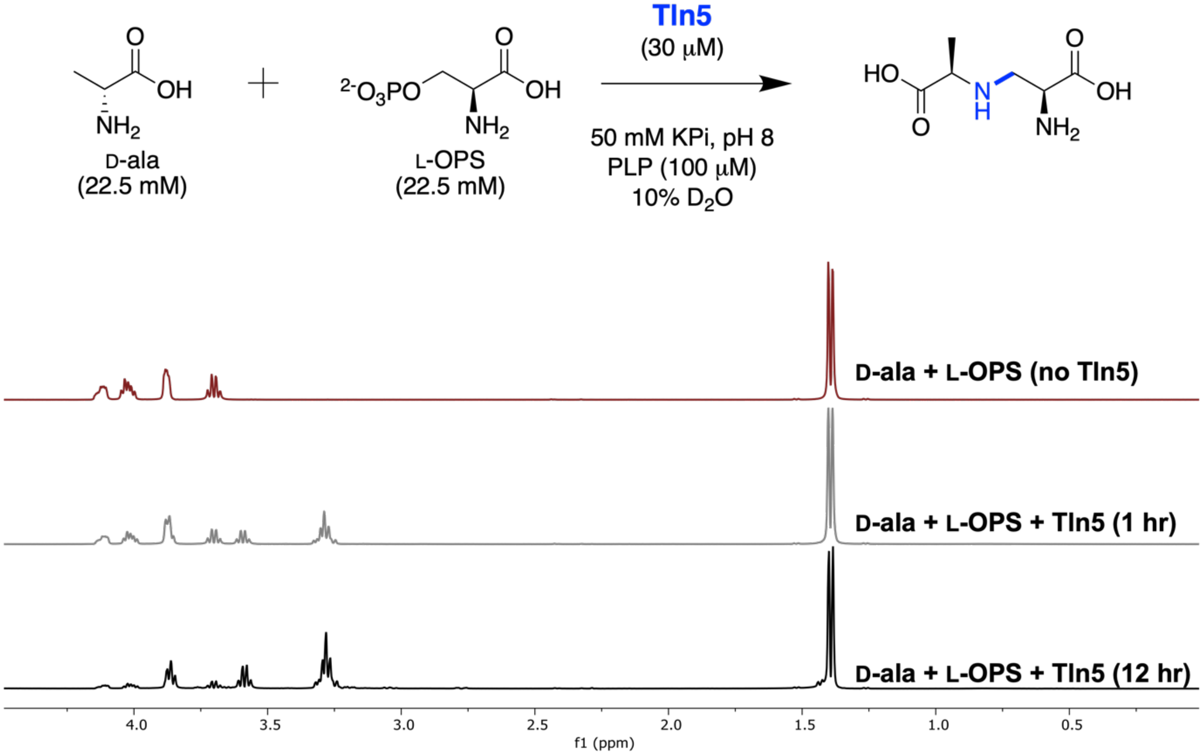


**Figure S13.** $^{1}$H NMR assay of Tln5 reveals the appearance of new $^{1}$H signals over the course of 12 h. The assay was set up using the 'Tln5 in vitro functional assays' protocol above but at the listed concentrations. $^{1}$H spectral readings were obtained after 1 and 12 h with the alteration of taking 128 scans for each reading.

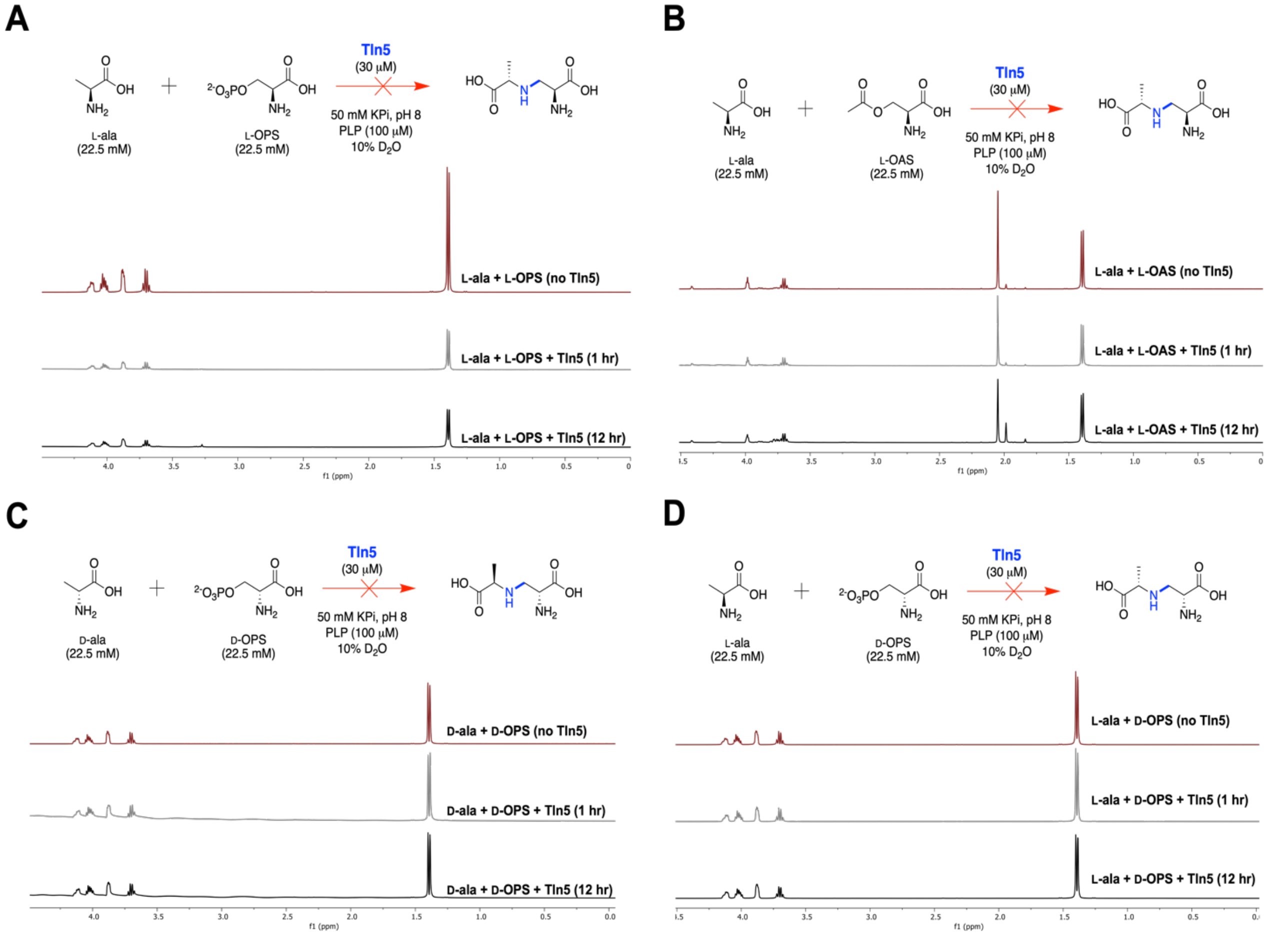


**Figure S14**. ¹H NMR assays of Tln5 determine which substrate permutations are not accepted by Tln5. Each assay was set up following the the 'Tln5 in vitro functional assays' protocol above but at the listed concentrations. Control samples with no Tln5 were obtained, as well as scans of samples 1 h and 12 h following the addition of Tln5. (A) L-alanine (L-ala) and L-*O*-phosphoserine (L-OPS) were screened, no new peaks formed after 12 h. (B) L-ala and L-acetylserine (L-OAS) were screened, no new peaks formed after 12 h. (C) D-alanine (D-ala) and D-*O*-phosphoserine (D-OPS) were screened, no new peaks formed after 12 h. (D) D-ala and D-OPS were screened, no new peaks formed after 12 h.

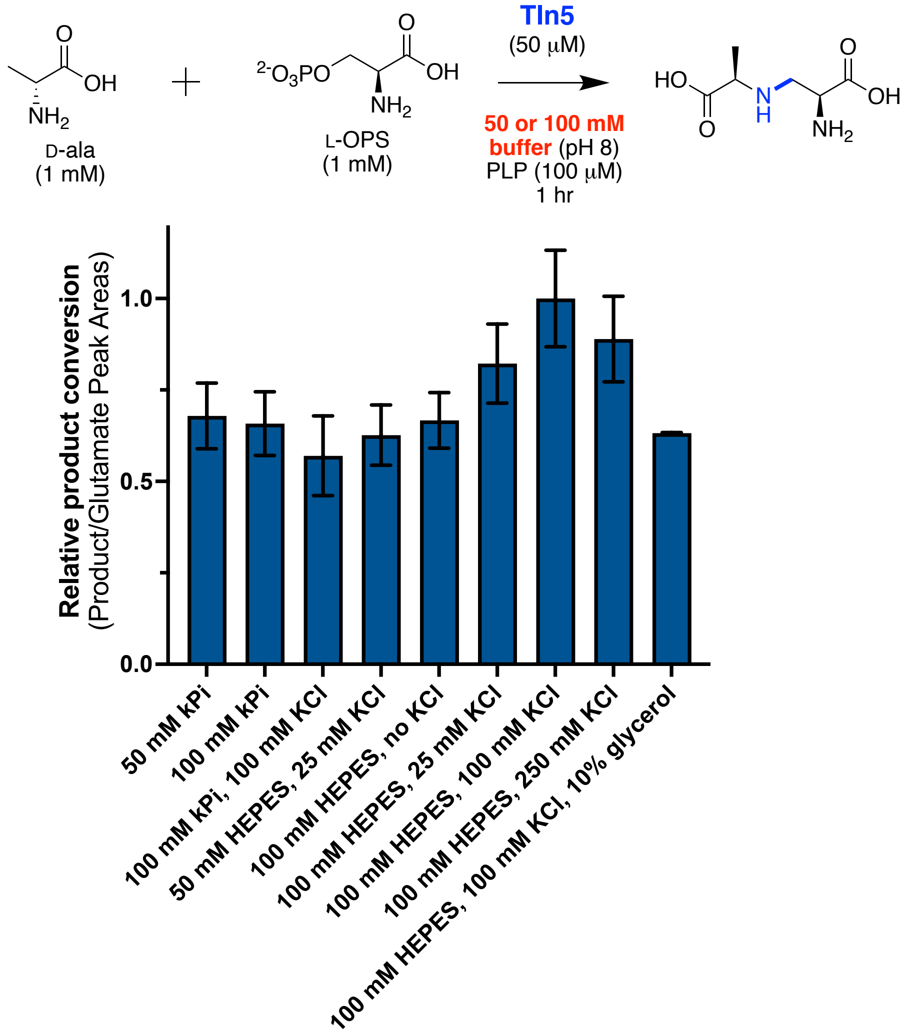


**Figure S15.** Tln5 buffer optimization reveals enzyme tolerance. The Tln5 buffer tolerance assay was set up in triplicate for each condition and incubated at room temperature for 1 h. An internal standard of 500 μM L-glutamic acid was added at the time of derivatization with Marfey's reagent and analyzed via UPLC-MS in positive mode ionization. Relative intensities for extracted ion chromatograms in positive mode (EIC ± 0.50 *m/z*) for Marfey-derivatized PDP and L-glutamic acid ($[M+H]^+$ 429.13, 400.11 *m/z*, respectively). The peak area for the PDP ion was calculated, then normalized to the internal standard. The averages of the relative product conversion for each condition is depicted with the standard deviation depicted as error bars. The selected condition for subsequent kinetics assays was the 100 mM HEPES, 100 mM KCl (pH 8.0) condition.

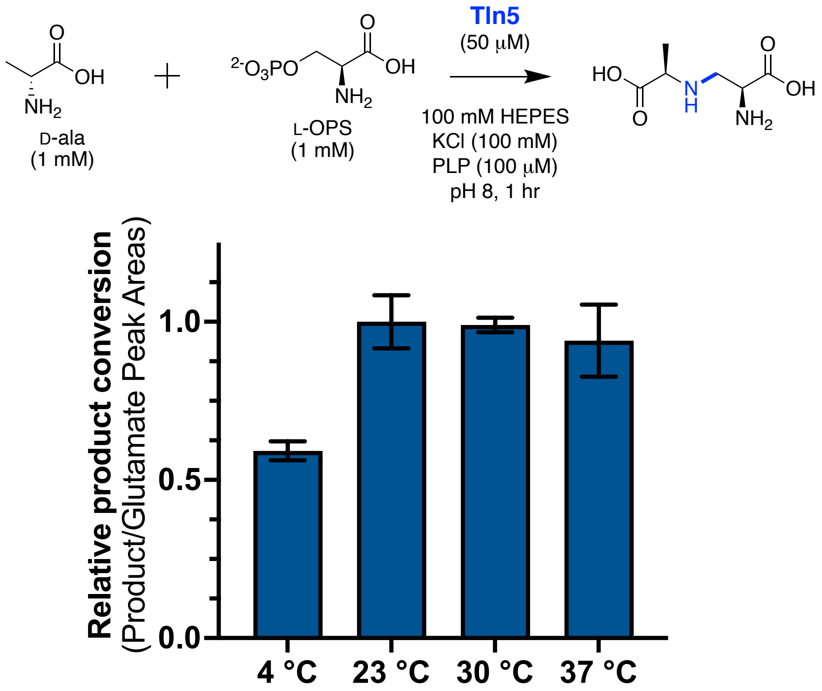


**Figure S16.** Tln5 temperature screen reveals preference at ambient or slightly elevated temperature. (≥23 °C). Assays were set up using the optimized buffer (**Figure S15**) and pH (**Figure S17**) conditions in triplicate and incubated for 1 h. A 500 μM L-glutamic acid internal standard was added prior to derivatization at the time of Marfey's reagent and subjected to UPLC-MS analysis. Extracted ion chromatograms were generated for Marfey-derivatized PDP ($[M+H]^+$ 429.13 ± 0.50 *m/z*) and L-glutamic acid ($[M+H]^+$ 400.11 ± 0.50 *m/z*), and the ratio of the peak areas were calculated to relatively quantify the amount of product. A temperature of 23 °C was selected for optimized steady-state kinetic assay conditions.

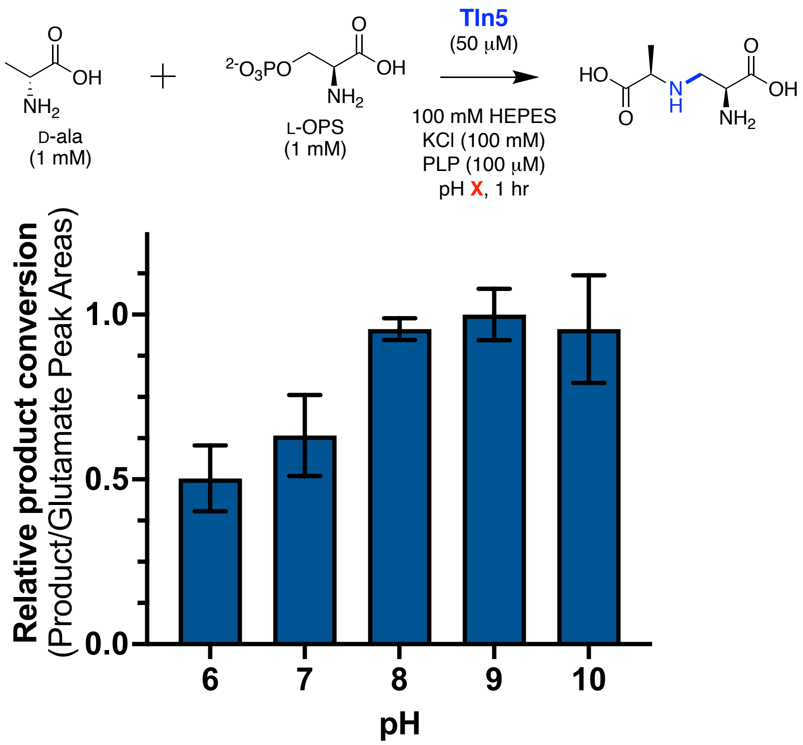


**Figure S17.** Tln5 pH screen reveals enzymatic preference towards basic conditions. Assays were set up using the optimized buffer conditions (**Figure S15**, 100 mM HEPES, 100 mM KCl) in triplicate and incubated for 1 h. A 500 μM L-glutamic acid internal standard was added at the time of derivatization with Marfey's reagent and analyses by UPLC-MS. Extracted ion chromatograms were generated for Marfey-derivatized PDP ($[M+H]^+$ 429.13 ± 0.50 *m/z*) and L-glutamic acid ($[M+H]^+$ 400.11 ± 0.50 *m/z*), and the ratio of the peak areas were calculated to relatively quantify the amount of product. Conditions above pH 8 and above had better product conversion compared to neutral and acidic conditions, and the pH 8 condition was selected for steady-state kinetic assay analysis of Tln5 due to the smallest standard deviation.

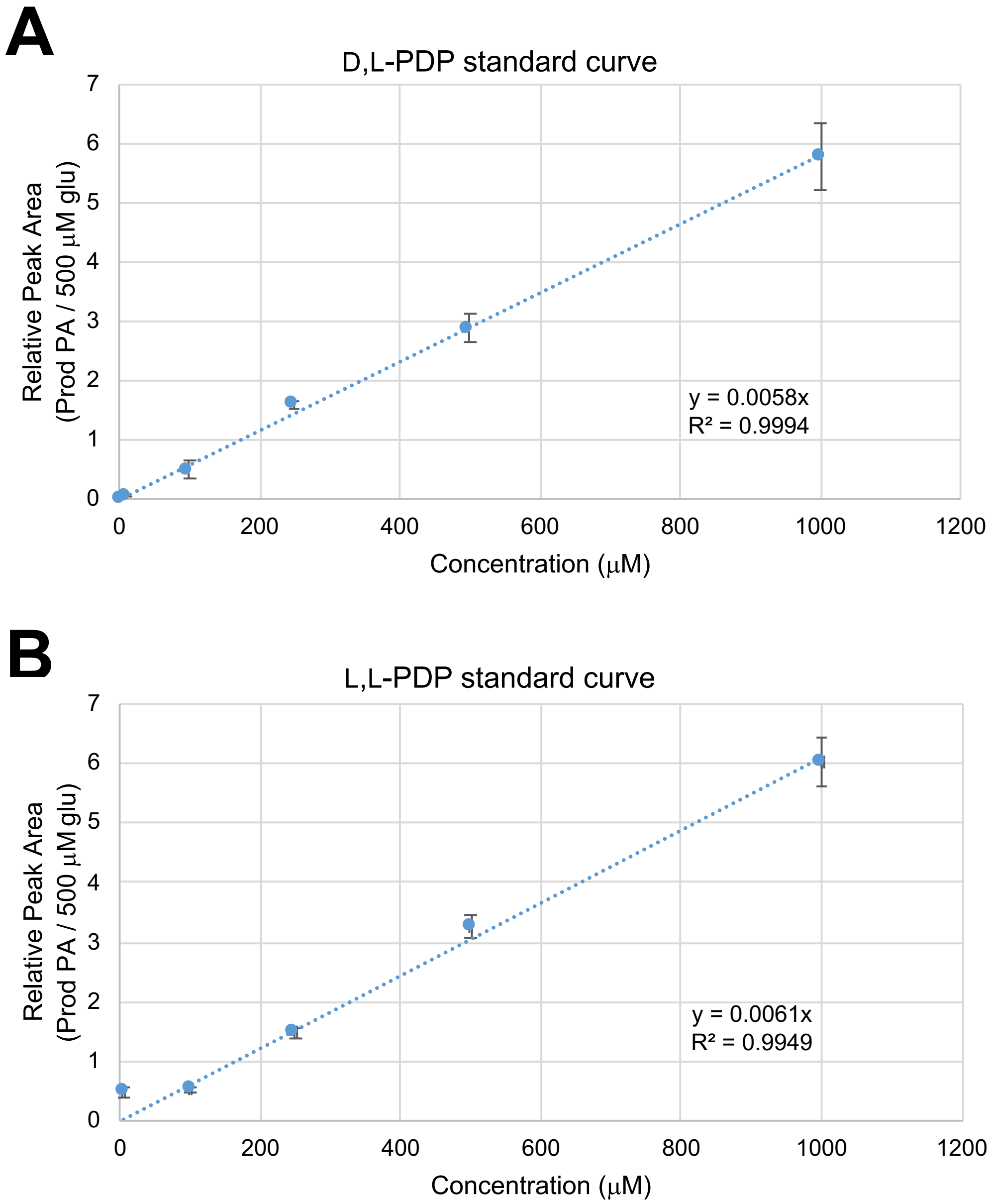


**Figure S18.** Standard curves for the (A) D,L- and (B) L,L-PDP. Standards of 10, 100, 250, 500, and 1000 μM of synthetically prepared PDP were set up in triplicate under the same reaction conditions omitting substrates and purified Tln5. Each sample was spiked with an internal standard of 500 μM L-glutamic acid and subsequently derivatized with Marfey's reagent for UPLC-MS analysis.

| **Tln5** | **AMA synthase** | **TrpB** |
|---|---|---|
| | | $k_{cat}$: 2.9 $s^{-1}$ |
| D-ala | Toxin A | indole |
| $K_m$: 0.702 ± 0.164 mM | $K_m$: 0.276 ± 0.04 mM | $K_m$: 0.020 mM |
| $k_{cat}$ = 0.072 ± 0.006 $s^{-1}$ | $k_{cat}$ = 0.104 ± 0.00261 $s^{-1}$ | |
| $k_{cat}$ / $K_m$ = 0.103 ± 0.036 $mM^{-1}s^{-1}$ | $k_{cat}$ / $K_m$ = 0.377 $mM^{-1}s^{-1}$ | $k_{cat}$ / $K_m$ = 330 $mM^{-1}s^{-1}$ |
| OPS | OPS | L-serine |
| $K_m$: 0.456 ± 0.162 mM | $K_m$: 1.313 ± 0.177 mM | $K_m$: 0.70 mM |
| $k_{cat}$ = 0.032 ± 0.004 $s^{-1}$ | $k_{cat}$ = 0.132 ± 0.005 $s^{-1}$ | |
| $k_{cat}$ / $K_m$ = 0.0702 ± 0.025 $mM^{-1}s^{-1}$ | $k_{cat}$ / $K_m$ = 0.101 $mM^{-1}s^{-1}$ | $k_{cat}$ / $K_m$ = not reported |

**Figure S19.** Steady state kinetic comparison data of Tln5 to other PLP-dependent β-substitution enzymes. Tln5 has comparable values to AMA synthase,[43] an enzyme of similar function to Tln5, in which both are kinetically slower when compared to engineered *Pyrococcus furiosus* TrpB.[44]

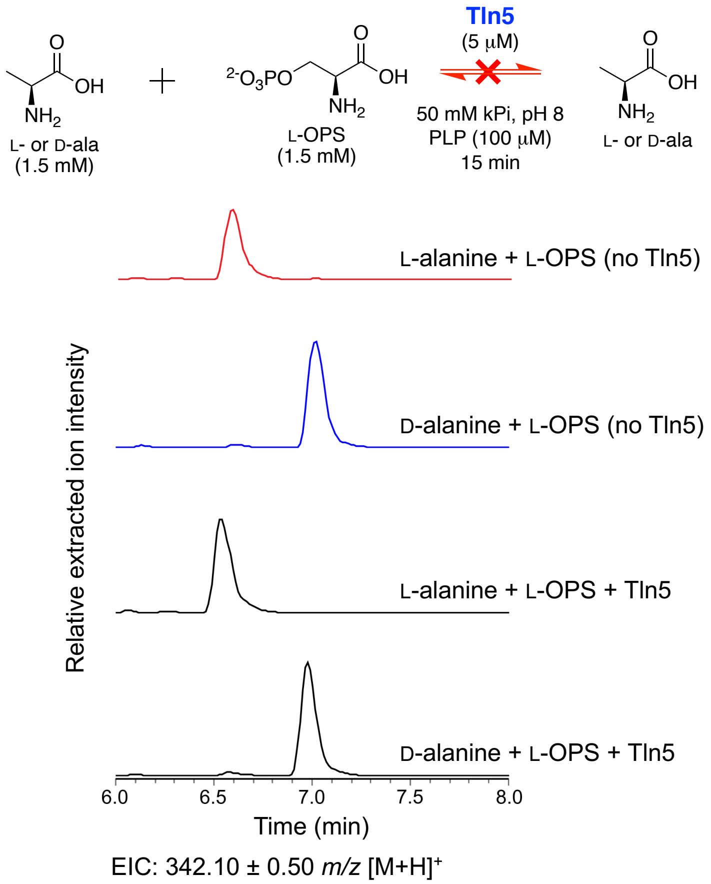


**Figure S20.** Tln5 does not catalyze the racemization of L- or D-alanine. In vitro Tln5 reactions were assembled using the protocols above then incubated at room temperature for 15 min. The assays were derivatized using Marfey's reagent for diastereomeric separation and improved retention times, then subjected to UPLC-MS analysis. The reactions were analyzed using UPLC-MS in positive mode ionization. Relative intensities of positive mode extracted ion chromatograms were extracted from UPLC-MS traces (EIC ± 0.50 *m/z*) for Marfey-derivatized alanine ($[M+H]^+$ 342.10 *m/z*). in the presence of Tln5 (black traces) a second peak does not appear for either trace that could align for the enantiomer of alanine.

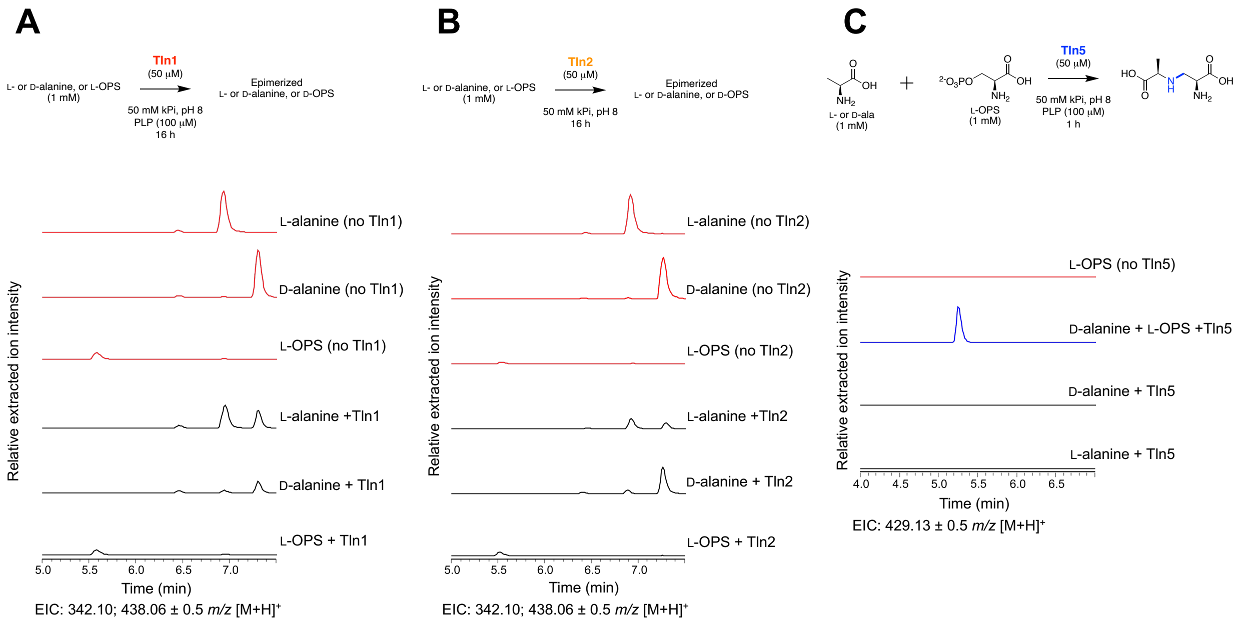


**Figure S21.** Alanine enantiomer activity assays for Tln1, Tln2, and Tln5. (A) Tln1 can react with L- and D-alanine, but not L-OPS. (B) Tln2 reacts with L- and D-alanine but not L-OPS. (C) The Tln5 reaction requires L-OPS to form the PDP product. Substrate dependency assays incubated at room temperature for 16 h, then derivatized with Marfey's reagent, and subjected to UPLC-MS analyses in positive ion mode. Relative intensities of extracted ion chromatograms in positive mode (EIC ± 0.50 *m/z*) are displayed for Marfey-derivatized PDP and alanine (429.13, 342.10 *m/z*, respectively) for Panels A and B, then only Marfey-derivatized PDP (429.13 *m/z*) for Panel C. EIC traces of no enzyme control samples are shown in red and samples containing the respective enzyme are in black.

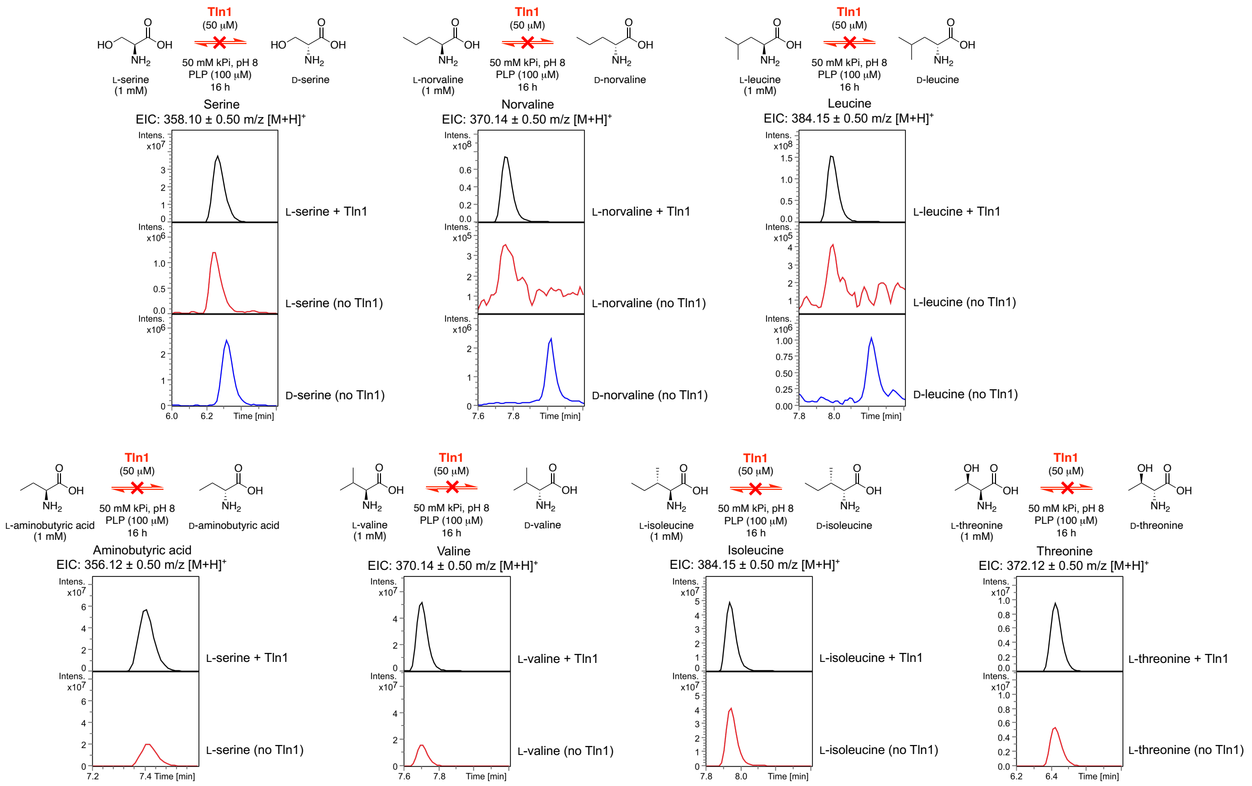


**Figure S22.** Tln1 alternative substrate screen suggests Tln1 does not racemize other amino acids. In vitro enzyme assays were assembled using the protocols above with the adaption of using other L-amino acids rather than alanine. Assays were incubated at room temperature for 16 h then derivatized using Marfey's reagent for improved retention time and diastereomeric separation. Derivatized assays were subjected to UPLC-MS analysis, in which relative intensities of positive mode extracted ion chromatograms were generated from UPLC-MS traces (EIC ± 0.50 *m/z*) for Marfey-derivatized serine, norvaline, leucine, etc ($[M+H]^+$ 358.10, 370.14, 384.15, 356.12, 370.14, 384.15, 372.12, respectively), depending on the substrate used in the assay condition. Some reactions were compared to a positive control using the D-enantiomer of the amino acid (blue), in which none of the peaks aligned with the experimental assay (black), or the assay omitting Tln1 (red).

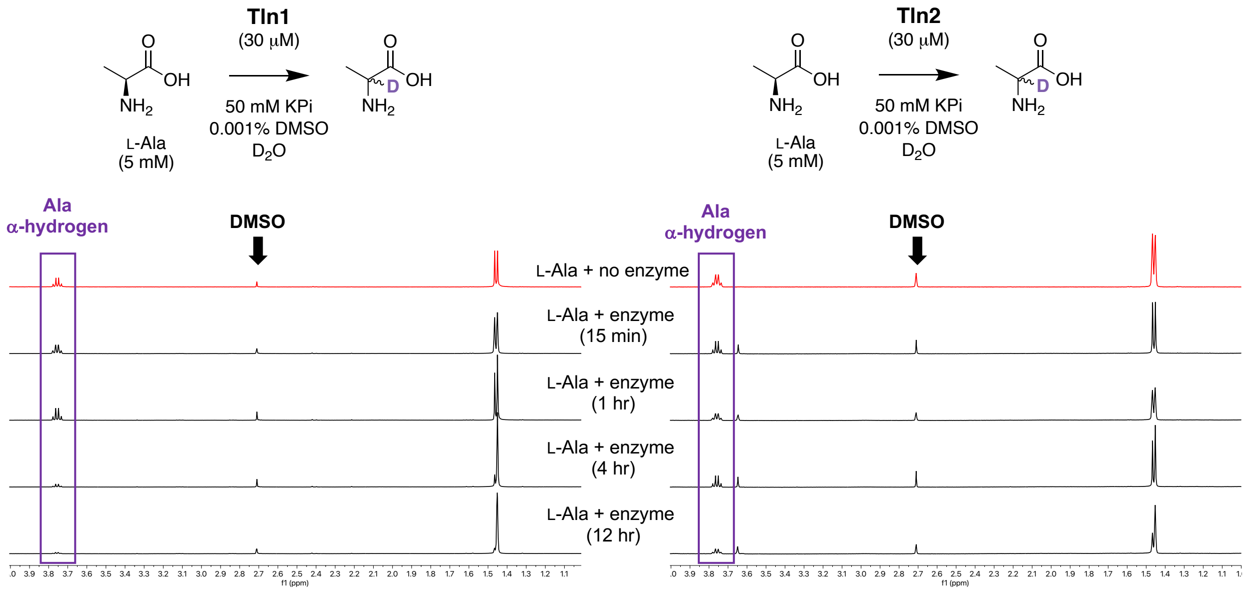


**Figure S23**: In vitro 1H NMR Tln1 or Tln2 assay with L-alanine in buffered $D_2O$. In vitro enzyme assays were set up in a ~167:1 substrate:enzyme molar ration in buffered $D_2O$ and analyzed using 500 MHz $^1$H NMR over the time points listed. The intensity of the alanine alpha-hydrogen signal (purple box) was compared to the 0.001% DMSO internal standard (black arrow) and quantified over time.

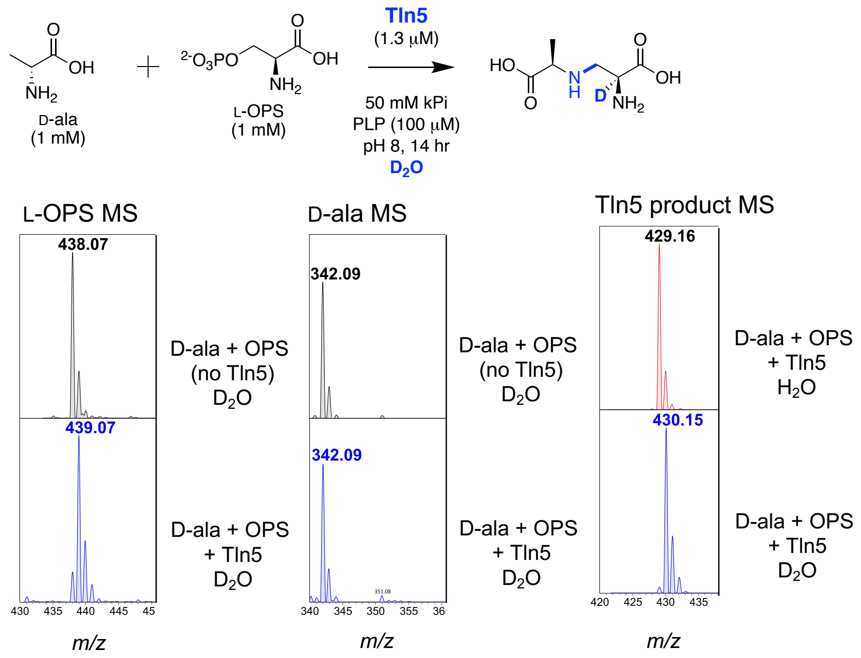


**Figure S24.** Deuterium incorporation assay of Tln5 indicates one deuterium atom is incorporated into the product of Tln5. In vitro enzymatic assays were assembled using protocols detailed above, with the assay conducted in >95% $D_2O$ conditions. After overnight incubation, assays were derivatized with Marfey's reagent, then subjected to UPLC-MS analysis. From our assay, a +1 Da shift is observed in the presence of Tln5 for both L-OPS and the PDP product, indicating that one deuterium was incorporated into L-OPS and the product. Our data aligns with the proposed PLP-dependent β-substitution mechanism in that only one reversible deprotonation is occurring, presumably at the step that forms the external aldimine intermediate between L-OPS and PLP.

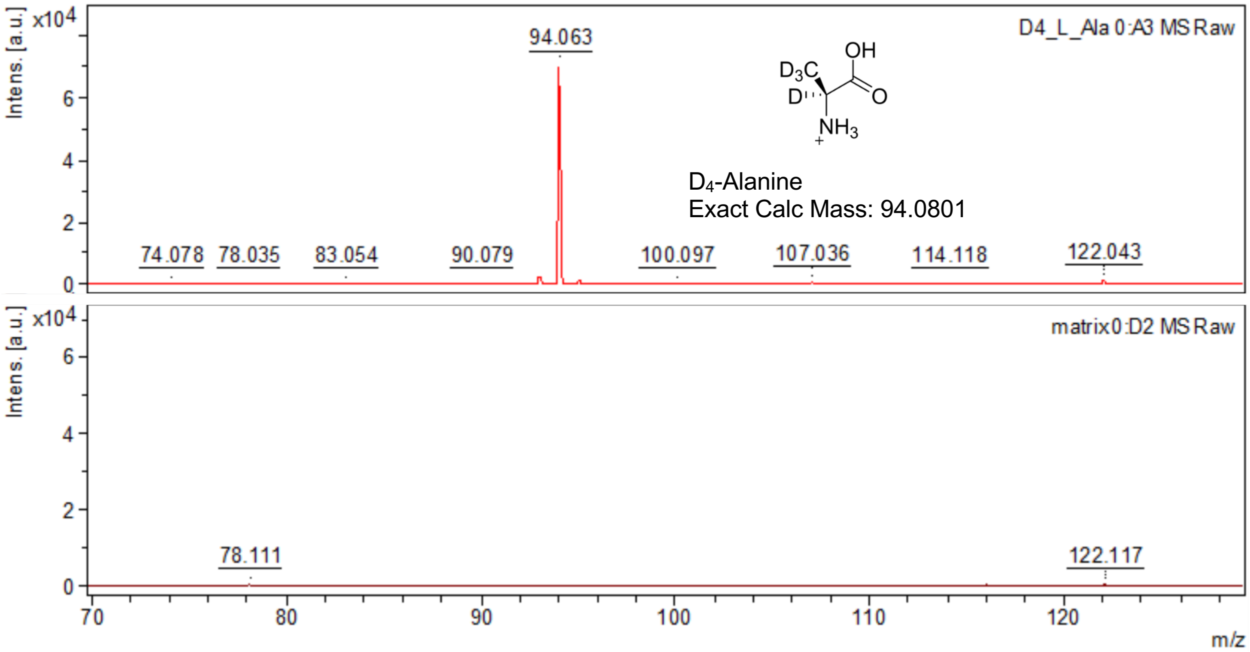


**Figure S25.** Dried droplet spectrum of the commercial $D_4$-Alanine measured on a Bruker Autoflex LRF MALDI-TOF mass spectrometer (top spectrum). This confirms the correct mass of the reagents used in the MSI experiments. The bottom spectrum corresponds to the matrix (CHCA:DHB) matrix used in subsequent MSI experiments displaying no overlap with background ions. This was conducted in positive, reflectron mode.

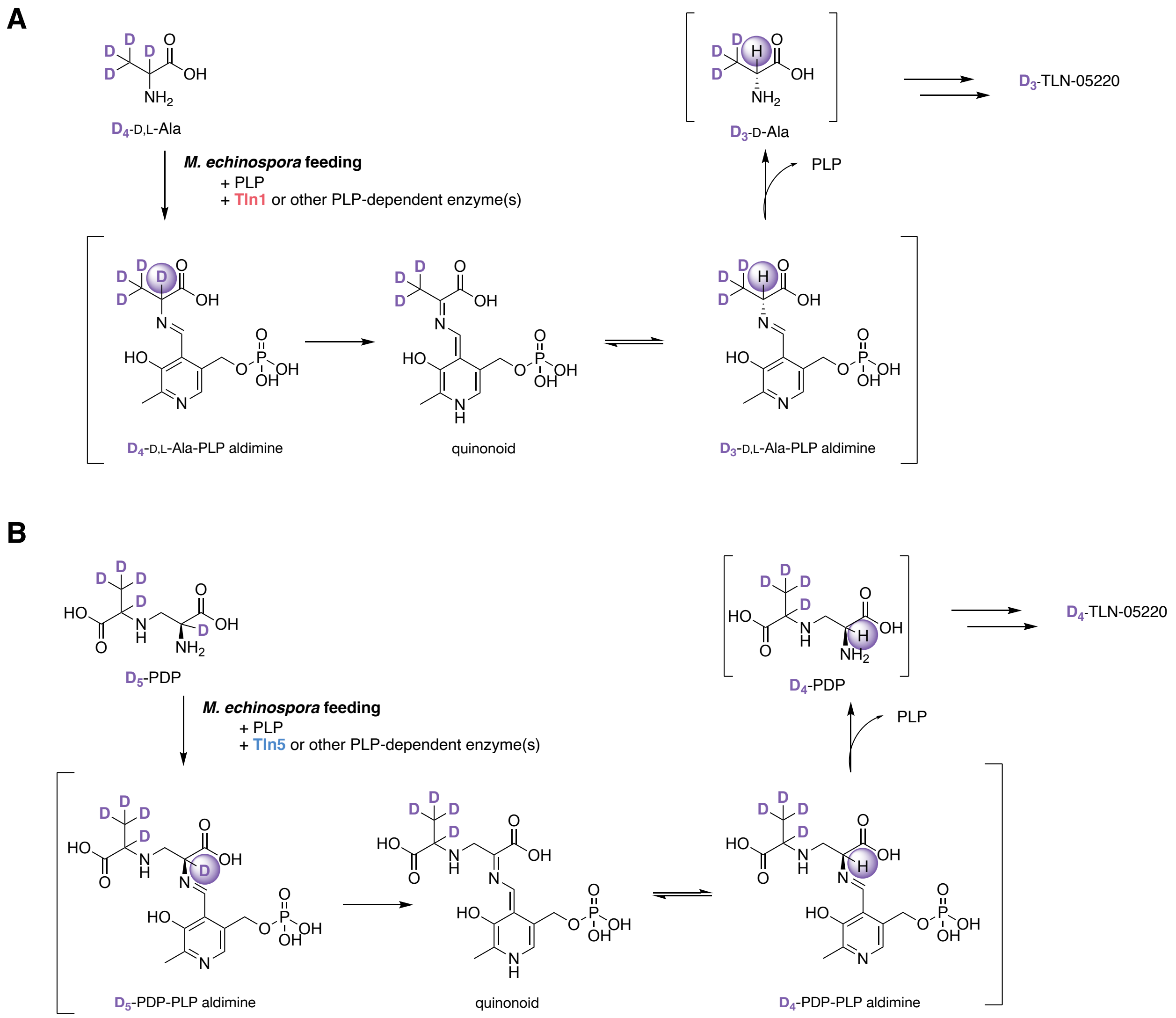


**Figure S26.** Proposed mechanism for the α-deuterium 'wash out' during *M. echinospora* feeding experiments with exogenously supplied **A**) $D_4$-DL-Ala and **B**) chemoenzymatically prepared $D_5$-PDP. Formation of the external PLP aldimine activates the α-deuterium adjacent to the primary amine in both substrates, enabling its abstraction and reprotonation via the established reversible aldimine to quinonoid mechanism. **A**) Tln1 putatively epimerizes this intermediate to generate intracellular $D_3$-D-Ala for subsequent incorporation into $D_3$-TLN-05220. **B)** Tln5 or another PLP-dependent enzyme could facilitate the α-deuterium removal and reversible reprotonation to generate intracellular $D_4$-PDP for subsequent incorporation into $D_4$-TLN-05220.

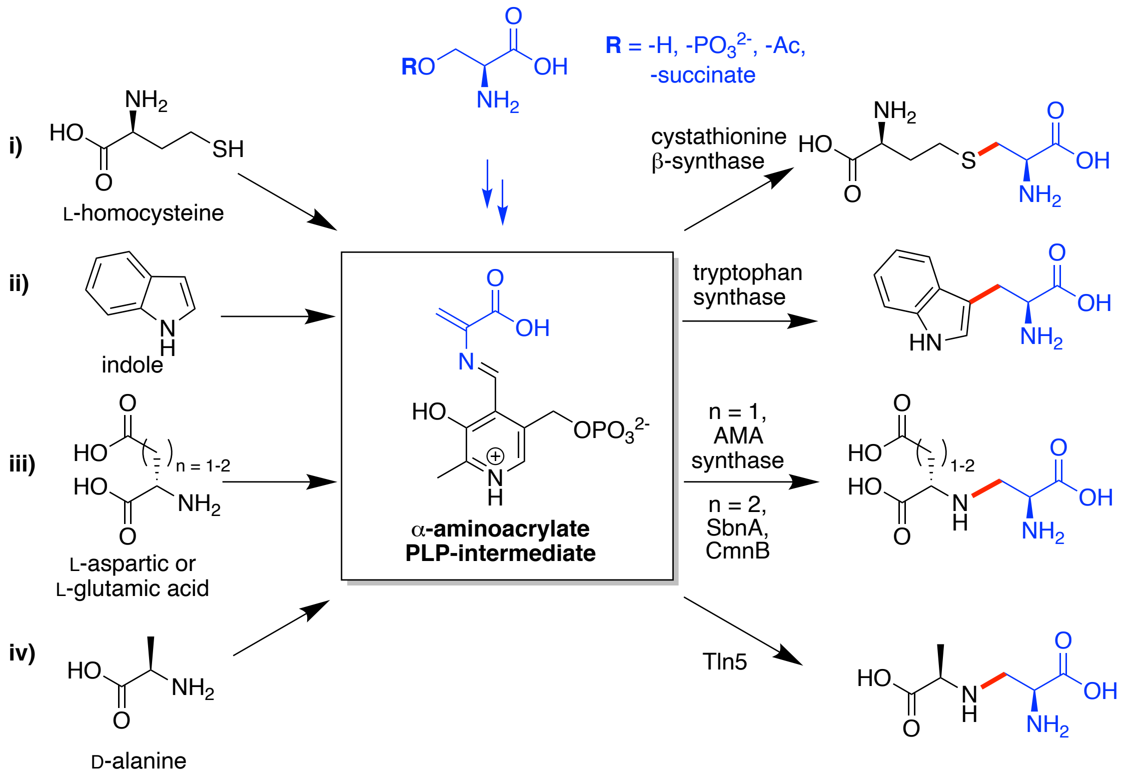


**Figure S27.** Examples of PLP-dependent β-substitution enzymes in primary and secondary metabolism.

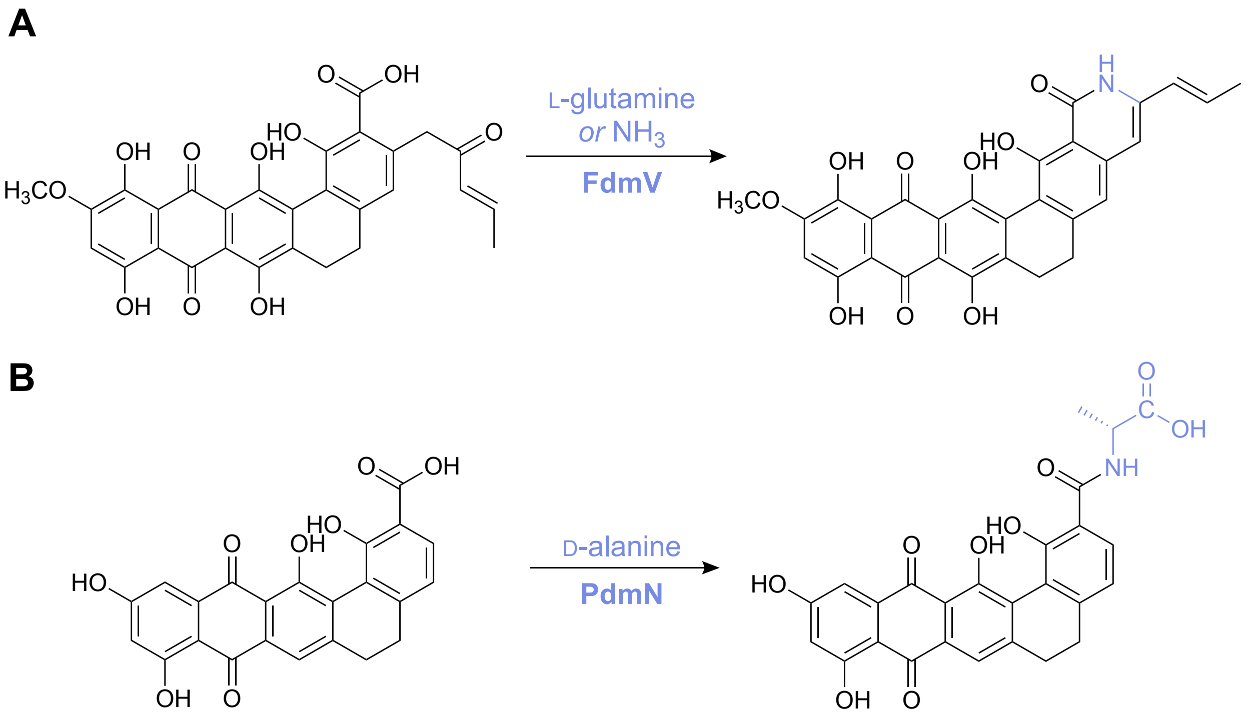


**Figure S28.** Known reactions performed by asparagine synthetase homologs in pentagular polyphenol biosynthesis. (A) FdmV in the fredericamycin pathway catalyzes an amidotransferase reaction with L-glutamine or free ammonia as the nitrogen donor to form fredericamycin B.[45] (B) PdmN in the pradimicin pathway ligates D-alanine to the carboxylic acid of a polyketide intermediate. PdmN was also found to accept D-serine.[31]

### Supplementary Tables

**Table S1.** Minimal inhibitory concentrations (MICs) of TLN-05220, TLN-052230, and the last resort antibiotic vancomycin[10] against Gram-positive bacteria.

| Bacterial Strain[a] | MIC (nM)[d] | | |
|---|---|---|---|
| | **TLN-05220[e]** | **TLN-05223[e]** | **vancomycin** |
| *S. aureus* ATCC 6538P | 22 | 85 | 690 |
| *S. aureus* MRS3 ATCC 700699[b] | 22 | 170 | 2760 |
| *S. pneumoniae* LSPQ13412 | 691 | 1356 | 345 |
| *S. pneumoniae* PenR LSPQ 3349[c] | 5527 | 1356 | 345 |
| *E. faecalis* VRE-1 ATCC 29212 | 87 | 1356 | 2760 |
| *E. faecalis* VRE-2 ATCC 51299 | 173 | 1356 | 22080 |
| *Clostridium difficile* ATCC 9689 | 83 | 2713 | 345 |

[a]*Staphylococcus aureus* (*S. aureus*); *Streptococcus pneumoniae* (*S. pneumoniae*); *Enterococcus faecalis* vancomycin-resistant *Enterococcus* (*E. faecalis* VRE); American Type Culture Collection (ATCC); Laboratoire de Santé Publique du Québec (LSPQ)

[b]methicillin-resistant Staphylococcus type III (MRS3)

[c]penicillin-resistant (PenR)

[d]MICs are reported as nanomolar (nM) concentrations, converted from µg/mL reported in the original reference[6] using their molecular weights (723.6910, 737.2221, 1449.2650 g/mol).

[e]Cells colored purple denote a significantly lower MIC (≥8-fold less) compared to vancomycin on a color scale from 8–128-fold difference.

**Table S2.** Protein accession codes for KS–CLF query and strain of interest.

| Protein | ***Micromonospora echinospora* subsp. *challisensis*** | ***Micromonospora echinospora* ATCC 15837** | **Sequence Identity (%)** |
|---|---|---|---|
| TlnL ($KS_\alpha$) | ADB23391.1 | WP_088985096.1 | 97.4 |
| TlnM (CLF or $KS_\beta$) | ADB23392.1 | WP_088985097.1 | 96.3 |

**Table S3.** *M. echinospora* ATTC 15837 homologous TLN-05220 cluster gene table with putative enzyme functions.

| Locus Tag[a] | Gene | Putative enzyme product function[b] | Protein homologs in MiBIG 4.0 database (% similarity/% identity), BGC accession, molecule name[c] |
|---|---|---|---|
| 6572 | *tln1* | PLP-dependent alanine racemase | BEV36910.1 (47/40), 2919, cirratiomycin A |
| 6573 | *tln2* | PLP-independent alanine racemase | KKZ73245.1 (45/23), 1778, showdomycin C |
| 6574 | *tln3* | Multicopper oxidase | XanP (69/55), 279, xantholipin |
| 6576 | *tln4* | Amino acid ligase | PdmN (58/43), 256, pradimicin A |
| 6577 | *tln5* | PLP-dependent serine-alanine ligase | SapC (57/43), 2510, youssoufene A1 |
| 6578 | *tln6* | Metallopeptidase | OE174828.1 (44/24), 2977, aminocoumacins D |
| 6589 | *tlnA* | Multicopper oxidase | Orf3 (55/45), 224, fredericamycin A |
| 6590 | *tlnB* | Acyl-ACP thioesterase | San2 (57/48), 190, A-74528 |
| 6591 | *tlnC* | Anthrone-type oxygenase | PnxE2 (79/60), 222, FD-594 |
| 6593 | *tlnD* | MFS transporter | AtI (62/43), 809, AT2433-A1 |
| 6594 | *tlnE* | Anthrone-type oxygenase | PnxE1 (70/53), 222, FD-594 |
| 6595 | *tlnF* | Amino acid ligase | PdmN (71/56), 256, pradimicin A |
| 6596 | *tlnG* | *O*-methyltransferase | PdmT (68/54), 256, pradimicin A |
| 6598 | *tlnH* | Ketoreductase | PnxG (71/57), 222, FD-594 |
| 6599 | *tlnI* | Dehydrogenase | XanS1 (75/60), 279, xantholipin |
| 6600 | *tlnJ* | TcmI-like cyclase | SanD (86/75), 190, A-74528 |
| 6601 | *tlnK* | TcmJ-like cyclase | PnxL (76/59), 222, FD-594 |
| 6602 | *tlnL* | Ketosynthase α-subunit | PnxA (83/74), 222, FD-594 |
| 6603 | *tlnM* | Ketosynthase β-subunit | SanG (76/61), 190, A-74528 |
| 6604 | *tlnN* | Acyl Carrier Protein | RubC (74/51), 266, collinomycin |
| 6605 | *tlnO* | TcmN-like cyclase | WP_029025952.1 (80/66), 198, arenimycin A |
| 6606 | *tlnP* | C19 Ketoreductase | PnxG (78/63), 222, FD-594 |
| 6607 | *tlnQ* | PdmH-like monooxygenase | PnxH (77/59), 222, FD-594 |
| 6608 | *tlnR* | PdmI-like monooxygenase | ACZ87076.1 (74/58), 2732, WS 79089B |
| 6609 | *tlnS* | Ketosynthase III | CosE (72/59), 1074, cosmomycin D |
| 6610 | *tlnT* | Peptide MFS transporter | Rub3 (68/55), 266, collinomycin |
| 6613 | *tlnU* | FAD-dependent monooxygenase | RubL (65/54), 266, collinomycin |
| 6614 | *tlnV* | Type I polyketide synthase | RdmG, (64/52), 1755, reedsmycin A |
| 6615 | *tlnW* | SARP family transcriptional regulator | AFJ52668.1, (81/73), 1073, kosinostatin C |
| 6617 | *tlnX* | MMPL family transporter | MrqT2 (73/56), 1675, murayquinone C |

[a]GA0070618_ precedes all gene locus tags listed. [b]Putative function assigned based on conserved function of homologs,[46,47] or data presented herein. [c]Closest protein homolog in the MiBIG 4.0 database[9] identified using BLAST+.[48] Hits from originally reported TLN-05220 cluster were excluded.

**Table S4.** AntiSMASH identified A-domains in NRPS/NRPS-like enzymes and the putative substrates in the ATCC 15837 genome (NCBI accession number LT607413.1).

| Region[a] | Gene Locus Tag[b] | NRPS A-domain position[c] | Predicted NRPS substrate(s)[d,e] | Nearest Stachelhaus code(s)[f] | Stachelhaus code match[g] |
|---|---|---|---|---|---|
| 1 | 0157 | 463..864 | Tyr, bOH-Tyr | Phe DAWVLAGIQK (50%) | 60% (w) |
| | 0161 | 14..367 | Ala, Gly, Val, Leu, Ile, Abu, Ival, Ser, Thr, Hpg, Dhpg, Cys, Pro, Hpr | Ala DLGIVGGMIK (26%) | 60% (w) |
| | 0168 | 489..878 | Ala, Gly, Val, Leu, Ile, Abu, Ival, Ser, Thr, Hpg, Dhpg, Cys, Pro, Hpr | Ala DAGGCAMVAK (50%), Pro DLFYLALVCK (50%) | 70% (w) |
| 10 | 3389 | 220..621 | Glu | Glu DVWHIGSIGK (62%), Val DFWNIGGIFK (47%) | 70% (w) |
| 16 | 4167 | 2709..3106 | Leu | Leu DAWIVGAIVK (62%) | 80% (m) |
| 17 | 4288 | 1190..1593 | (unknown) | Lys DAESVGTIIK (71%) | 80% (m) |
| | 4301 | 205..606 | (unknown) | Fo-OH-Orn DVWILGATNK (44%) | 80% (m) |
| | 4305 | 11..391 | Ser | Ser DVWSIAMVHK (71%) | 90% (m) |
| | 4311 | 11..407 | Val, Ala | Val DVFWLGGTFK (79%), Ala DVFWLGGTFK (79%) | 100% (s) |
| | 4315 | 500..903 | (unknown) | Lys DAEDIGTVSK (62%), Lys/Arg DVEDIGSVAK (59%) | 70% (w) |
| 19 | 4866 | 446..866 | Gly | Ala DILQIGQIYK (71%), Gly DILQVGMIWK (59%) | 70% (w) |
| | 4868 | 503..904 | Asp | Asp DLTKIGHVGK (71%) | 90% (m) |
| | 4934 | 20..362 | (unknown) | Acc DLCHVAIIAK (29%) | 60% (w) |
| | 4936 | 9..377 | Asp, Asn, Glu, Gln, Aad | | 0% (w) |
| | 4956 | 479..889 | Val | Val DALWLGGTFK (85%), bOH-Val DALWLGGTFK (82%) | 100% (s) |
| | 4957 | 468..857 | Gly | Ala DILQIGQIYK (74%) | 80% (m) |
| | 4960 | 512..907 | Ala | Ala DIVQLGLVYK (68%) | 90% (m) |
| | 4961 | 500..926 | Gly, Ala, Val, Leu, Ile, Abu, Ival | CysA DATKMGHVGK (56%) | 70% (w) |
| | 4972 | 36..191 | Ala, Gly, Val, Leu, Ile, Abu, Ival, Ser, Thr, Hpg, Dhpg, Cys, Pro, Hpr | D-Hiv GALMIVGSIK (29%) | 60% (w) |
| | 4975 | 85..259 | Ala, Gly, Val, Leu, Ile, Abu, Ival, Ser, Thr, Hpg, Dhpg, Cys, Pro, Hpr | Ala/D-Ala DLLFGISVLK (35%) | 60% (w) |

| | | | | | |
|---|---|---|---|---|---|
| 22 | 5227 | 519..941 | Gly, Ala, Val, Leu, Ile, Abu, Ival | *N*6-ohLys DALHPGHVCK (65%), CysA DATKMGHVGK (59%) | 70% (w) |
| | 5228 | 525..927 | Gly | Gly DILQLGVVWK (82%) | 100% (s) |
| | 5229 | 458..850 | Ala | Gly DIYHLGLVWK (62%), D-Ala DMPQLGMVWK (59%) | 60% (w) |
| | 5230 | 14..408 | Phe, Trp, Ph-Gly, Tyr, bOH-Tyr | | 0% (w) |
| 24 | 5334 | 499..903 | Thr | Thr DFWNIGMVHK (97%) | 100% (s) |
| | 5334 | 1572..1983 | Fo-OH-Orn | Fo-OH-Orn DINYWGGIGK (100%) | 100% (s) |
| | 5335 | 504..925 | Asn | Asn DFTKVGEVGK (65%) | 90% (m) |
| | 5336 | 39..448 | Sal | Sal PLPAQGVLNK (91%) | 100% (s) |
| | 5337 | 581..972 | Ser | Ser DLFNLGLIHK (94%) | 100% (s) |
| 25 | 5534 | 546..953 | Arg | Arg DVADVGAIDK (91%) | 100% (s) |
| | 5534 | 1606..2013 | Pro | Pro DMENVSHVVK (56%) | 80% (m) |
| | 5549 | 22..412 | Ala, Gly, Val, Leu, Ile, Abu, Ival, Ser, Thr, Hpg, Dhpg, Cys, Pro, Hpr | Leu DAWFLGNIVK (32%), Hpr DFQFFGVAVK (29%) | 60% (w) |
| | 5559 | 48..430 | (unknown) | | 0% (w) |
| | 5565 | 25..359 | Gly, Ala, Val, Leu, Ile, Abu, Ival | | 0% (w) |
| 27 | 5836 | 464..852 | Phe | His DTWTIASVDK (68%), Phe DAWTIAGVCK (65%) | 80% (m) |
| | 5836 | 1509..1906 | Dhpg, Hpg | Dhpg DAYNAGTLCK (59%) | 70% (w) |
| | 5838 | 476..871 | (unknown) | Gly DMLQIGTVWK (62%) | 80% (m) |
| 28 | 5902 | 474..863 | Phe | His DTWTIASVDK (68%), Phe DAWTIAGVCK (65%) | 80% (m) |
| | 5902 | 1516..1913 | Dhpg, Hpg | Dhpg DAYNAGTLCK (59%) | 70% (w) |
| | 5906 | 31..414 | Arg, Asp, Glu, Asn, Lys, Gln, Orn, Aad | | 0% (w) |
| 29 | 6232 | 496..902 | Ser | Ser DVWHISLIDK (97%) | 100% (s) |
| | 6233 | 455..849 | Asn | Asn DLTKVGEVGK (88%), OH-Asn/Asn DLTKVGEVGK (85%) | 100% (s) |
| | 6237 | 26..410 | (unknown) | Leu DALLVGAVAK (59%) | 80% (m) |
| | 6239 | 11..394 | (unknown) | Leu/Val DAIALGGCAK (41%) | 70% (w) |
| | 6262 | 525..926 | (unknown) | Trp/Tyr DVMFYTALVK (94%) | 90% (m) |
| | 6263 | 16..406 | Ala, Gly, Val, Leu, Ile, Abu, Ival, Ser, Thr, Hpg, Dhpg, Cys, Pro, Hpr | mxAnt AATHLSATLK (76%) | 70% (w) |

[a]antiSMASH version 8.0.2 identified 30 biosynthetic gene cluster regions with relaxed strictness.[11]
[b]GA0070618_ precedes all gene locus tags listed.
[c]Amino acid position range of the adenylation-domain (A-domain) in the non-ribosomal peptide synthetase (NRPS) or NRPS-like module.
[d]The antiSMASH support vector machine (SVM) substrate specificity prediction abbreviations are as follows: Ala, Alanine; Abu, 2-Aminobutyric acid; Aad, 2-Aminoadipic acid; AHBA, 3-Amino-5-hydroxybenzoic acid; Acc, 1-Aminocyclohexane-1-carboxylic acid; Arg, Arginine; Asn, Asparagine; Asp, Aspartic acid; bOH-Tyr, β-Hydroxytyrosine; bOH-Val, β-Hydroxyvaline; Cys, Cysteine; CysA, Modified cysteine analogue (often

cysteic acid or derivative); Dhb, Dehydrobutyrine; Dhpg, 3,5-Dihydroxyphenylglycine; D-Hiv, D-α-Hydroxyisovaleric acid; Fo-OH-Orn, *N*-Formyl-β-hydroxyornithine; Gly, Glycine; Hpg, 4-Hydroxyphenylglycine; Hpr, 4-Hydroxyproline; His, Histidine; Ile, Isoleucine; Ival, Isovaline; Leu, Leucine; Lys, Lysine; mal, Malonyl-CoA derived extender unit; mmal, Methylmalonyl-CoA derived extender unit; mxAnt, Modified anthranilic acid derivative; *N*6-ohLys, *N*6-Hydroxylysine; OH-Asn, Hydroxyasparagine; Orn, Ornithine; Phe, Phenylalanine; Ph-Gly, Phenylglycine; pk, Polyketide extender unit (unspecified); Pro, Proline; Sal, Salicylic acid; Ser, Serine; Thr, Threonine; Trp, Tryptophan; Tyr, Tyrosine; Val, Valine.

[e]Amino acids are the L enantiomer unless specified in the naming convention.

[f]List of the predicted substrate abbreviation and reference Stachelhaus code sequence (match score). The match score is the percentage of amino acid residues in the A-domains substrate-binding pocket that match those of the reference Stachelhaus code with an 8 Å distance cutoff from a modelled bound ligand.

[g]Percent of the 10 amino acids in Stachelhaus positions that match the reference code. Confidence abbreviations: (w) = weak match (<70%); (m) = moderate match (70–89%); (s) = strong match (≥90%).

**Table S5.** Putative ATCC 15837 protein functions by National Center for Biotechnology Information (NCBI) and InterPro.[49]

| Protein | NCBI protein accession | NCBI Definition | InterPro family |
|---|---|---|---|
| Tln1 | WP_088985070.1 | PLP-dependent lyase/thiolase | Cysteine synthase/Cystathionine beta-synthase (IPR050214) |
| Tln2 | WP_157749058.1 | maleate cis-trans isomerase family protein | Maleate isomerase/Arylmalonate decarboxylase (IPR026286) |
| Tln4 | WP_088985074.1 | Asparagine synthetase B family protein | Asparagine Synthetase/Amidase (IPR051786), Asparagine synthase, glutamine-hydrolyzing (IPR006426) |
| Tln5 | WP_088985075.1 | PLP-dependent cysteine synthase family protein | Cysteine synthase/Cystathionine beta-synthase (IPR050214) |
| TlnF | WP_088985089.1 | asparagine synthase (glutamine-hydrolyzing) | Asparagine synthase, glutamine-hydrolyzing (IPR006426), Asparagine Synthetase/Amidase (IPR051786) |

### <u>Synthetic Chemistry Protocols</u>

#### Synthesis of *O*-succinyl-L-serine (SI-2)

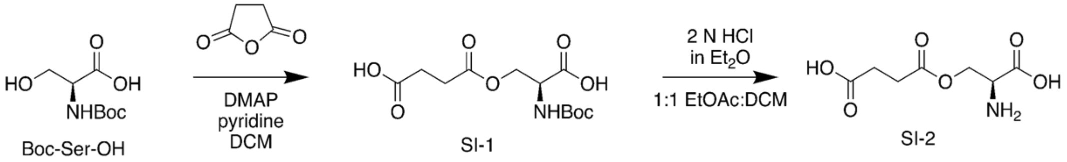


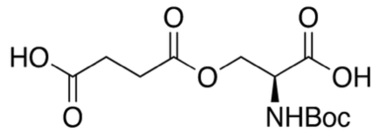

**SI-1**

The protocol used to prepare **SI-2** was adapted from a reference.[50] To a solution of *N*-Boc-L-serine (1.004 g, 4.89 mmol) in dry DCM (60 mL), DMAP (0.060 g, 0.49 mmol) and succinic anhydride (588 mg, 5.88 mmol) were added sequentially. Allow the reaction to stir at room temperature for 18 h, then concentrate the reaction in vacuo. After concentration, resuspend the reaction mixture in 1:1 $H_2O$: EtOAc (60 mL) and checked for acidity. If the reaction mixture is not acidic, add 1 N HCl until the pH is 3-4. The acidified aqueous layer was extracted with EtOAc (2 x 60 mL), pool the organic layers, dry over $MgSO_4$, then concentrate in vacuo. The crude mixture was purified first via flash column chromatography using a 5%–10% $CH_3OH$ in DCM with 0.1% acetic acid gradient. Afterword, a second purification was performed via flash column chromatography using a 5-10% $CH_3OH$ in $CHCl_3$ with 0.1% acetic acid column to yield the product as a white solid (0.156 g, 10%). $^1$H NMR (500 MHz, $CD_3OD$) $\delta$ 4.50–4.41 (m, 2H), 4.34 (dd, J = 10.7, 5.2 Hz, 1H), 2.65–2.60 (m, 4H), 1.47 (s, 9H). $^{13}$C NMR (126 MHz, $CD_3OD$) $\delta$ 176.15, 173.84, 157.82, 80.79, 65.19, 54.27, 29.89, 29.85, 28.68. HRMS (MALDI) Calculated for $C_{12}H_{20}NO_8$ 306.1183, found 306.1687 $[M+H]^+$.

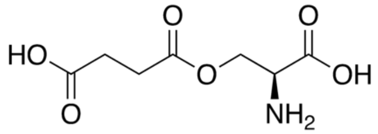

**SI-2**

A stirred solution of **SI-1** (0.058g, 0.19 mmol) was prepared in 1:1 EtOAc: dry DCM (1.14 mL) and cooled to 10 °C. Afterword, 2 N HCl in $Et_2O$ (1.43 mL) was added to the solution, and the reaction stirred for 6 h at room temperature. The reaction was concentrated in vacuo, then lyophilized overnight to yield a white solid (0.008 g, 17%). $^1$H NMR (500 MHz, $D_2O$ + 0.1% $CH_3OH$) $\delta$ 4.66 (dd, *J* = 12.5, 4.7 Hz, 1H), 4.59 (dd, *J* = 12.5, 2.9 Hz, 1H), 4.41 (dd, *J* = 4.5, 3.1 Hz, 1H), 2.74 (m, *J* = 3.5 Hz, 4H). $^{13}$C NMR (126 MHz, $D_2O$ + 0.1% $CH_3OH$) $\delta$ 176.92, 174.12, 169.56, 62.42, 52.55, 28.65. HRMS (MALDI) Calculated for $C_7H_{12}NO_6$ 206.0659, found 206.0667 $[M+H]^+$.

**General Procedure A: Sulfamidate ring opening**
This protocol was adapted from the following references.[51,52] A stirred solution of sulfamidate (1 eq) in THF (1 mL) was prepared and heated to 50 °C. A solution of desalted free amine (2 eq) in $CH_3CN$ (1 mL) was prepared and added to the reaction flask. An additional aliquot of 1:1 THF:$CH_3CN$ was added to the reaction vessel to minimize solvent evaporation. After overnight incubation (14 h) at 50 °C, the crude reaction was concentrated in vacuo, then purified via flash column chromatography using a 1–4% $CH_3OH$ in $CHCl_3$ gradient eluent system to yield the final product as a yellow oil.

**General Procedure B: Product deprotection procedure**
A solution of protected dipeptide (1 eq) was prepared in DCM (5 mL), then trifluoroacetic acid (2.5 mL) was added and the reaction mixture stirred at room temperature overnight (14 h). The reaction was concentrated in vacuo, then washed with DI water (2 x 5 mL), then redissolved in DI water (5 mL) prior to overnight lyophilization. After lyophilization, the reaction mixture was redissolved to a concentration of 15 mg/mL in DI water, then purified via HPLC. The crude mixture was purified via HPLC using a Synergi 4 um Polar-RP 80 A LC column (250 x 10 mm, Manufacturer) at a flow rate of 2.5 mL/min using the following gradient system to yield final products as white solids. The HPLC method is as follows: 3 min at 10% B, 7 min at 10–50% B, 3 min at 50–100% B, 2 min at 100% B, 2 min 100–10% B, 3 min.

**Synthesis of D,L-PDP Tln5 diastereomer (SI-6)**

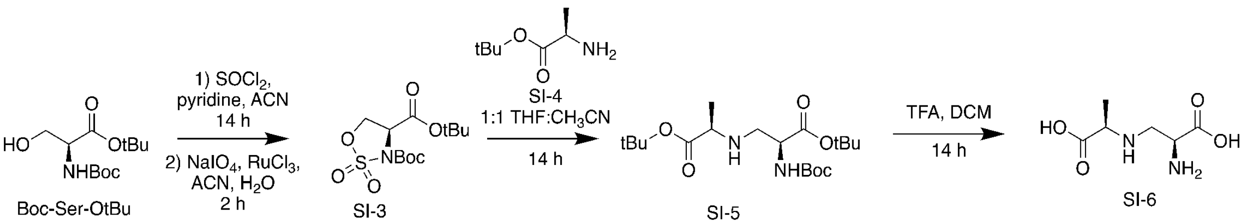


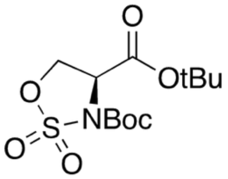

**SI-3**

The protocol was adapted from the following references.[51,52] To a stirred solution of $SOCl_2$ (0.28 mL, 3.83 mmol) in $CH_3CN$ (6.5 mL) at −40 °C, Boc-ser-*O*-tBu (0.399 g, 1.53 mmol) was added as a solution in dry $CH_3CN$ (16 mL) over the course of 20 min under Ar (g). After stirring for 1.5 h at −40 °C, dry pyridine (0.74 mL, 9.18 mmol) was added over 5 min. The reaction was slowly brought up to room temperature and stirred for an additional 16 h, before quenching the reaction by adding ice water (5 mL). The aqueous layer was extracted with EtOAc (2 x 25 mL), the organic layers were pooled, washed with water (7 mL), brine (10 mL), dried over $Na_2SO_4$, filtered, then dried in vacuo to yield the crude reaction mixture as an oil. The crude mixture was taken into the next step without further characterization or purification, where the mixture was dissolved in 1:1 $CH_3CN$: $H_2O$ (13 mL), and $RuCl_3$ (0.018 g, 0.09 mmol) and $NaIO_4$ (0.288 g, 1.43 mmol) were added sequentially. The reaction stirred at 0 °C for 2.25 h, then diluted with EtOAc (16 mL). The aqueous layer was extracted with EtOAc (2 x 10 mL), then the organic layers were pooled, washed with brine (15 mL), dried over $Na_2SO_4$, then concentrated in vacuo. The crude reaction mixture was purified via flash column chromatography using a 4:1 hexanes: EtOAc eluent system to yield the final product as a white solid (0.282 g, 65%). $^1$H NMR (500 MHz, $CDCl_3$) δ 4.76–4.71 (m, 1H), 4.63 (d, *J* = 8.1 Hz, 2H), 1.56 (s, 9H), 1.51 (s, 9H). $^{13}$C NMR (126 MHz, $CDCl_3$) δ 166.16, 86.02, 84.64, 67.90, 58.27, 28.04, 27.97. HRMS (MALDI) Calculated for $C_{12}H_{21}NO_7SNa$ 346.0931, found 346.0945 $[M+Na]^+$.

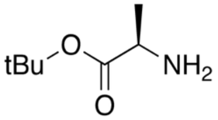


**SI-4**

The protocol was adapted from the following reference.[53] To a stirred solution of H-D-Ala-*O*-tBu hydrochloride salt (0.230 g, 1.26 mmol) in DCM (2.5 mL), aqueous saturated $Na_2CO_3$ (1.25 mL) was added at room temperature. After stirring for 15 min, the aqueous layer was extracted with DCM (2 x 2.5 mL), then the organic layers were pooled, washed with water (2.5 mL), brine (2.5 mL), dried over $Na_2SO_4$, filtered, then concentrated in vacuo. The crude product (0.103 g, 56%) was utilized without further purification for the next step.

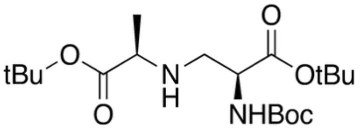


**SI-5**

**SI-3** (0.053 g, 0.17 mmol) and **SI-4** (0.060 g, 0.33 mmol) were used in General Procedure A to produce **SI-5** (0.056 g, 87%). $^1$H NMR (500 MHz, $CDCl_3$) δ 5.39 (d, *J* = 6.1 Hz, 1H), 4.18 (s, 1H), 3.16 (q, *J* = 6.9 Hz, 1H), 2.99 (dd, *J* = 12.2, 4.1 Hz, 1H), 2.83 (dd, *J* = 12.0, 3.5 Hz, 1H), 1.47 (s, 8H), 1.45 (s, 9H), 1.44 (s, 8H), 1.21 (d, *J* = 7.0 Hz, 3H). $^{13}$C NMR (126 MHz, $CDCl_3$) δ 174.85, 170.96, 155.59, 82.03, 81.13, 79.73, 57.57, 54.63, 49.13, 28.49, 28.21, 28.16, 19.10.
HRMS (MALDI) Calculated for $C_{19}H_{37}N_2O_6$ 389.2646, found 389.2634 $[M+H]^+$.

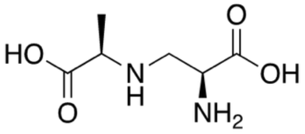


**SI-6**

**SI-5** was used to produce **SI-6** using General Procedure B. $^1$H NMR (500 MHz, $D_2O$ + 0.1% $CH_3OH$) δ 4.12 – 4.07 (m, 1H), 3.87 – 3.81 (m, 1H), 3.61 – 3.46 (m, 2H), 1.56 (d, *J* = 7.1 Hz, 3H); $^{13}$C NMR (126 MHz, $D_2O$ + 0.1% $CH_3OH$) δ 174.36, 170.80, 58.44, 49.63, 45.05, 14.90; HRMS (MALDI) Calculated for $C_6H_{13}N_2O_4$ 177.0870, found 177.0871 $[M+H]^+$.

**Synthesis of L,L-PDP Tln5 diastereomer (SI-9)**

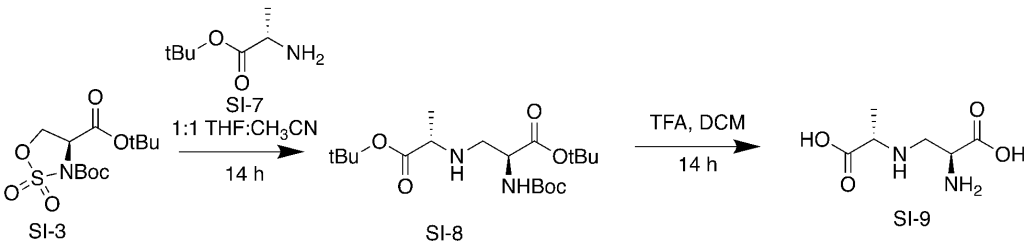


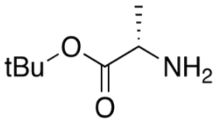


**SI-7**

H-Ala-*O*-tBu hydrochloride salt (0.451 g, 2.48 mmol) was used to produce **SI-7** (0.229 g, 63%) with the above desalt protocol. This product was used in downstream reactions without further purification.

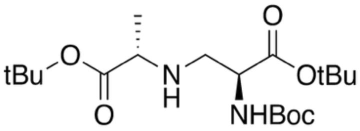


**SI-8**

**SI-3** (0.084 g, 0.26 mmol) and **SI-7** (0.075 g, 0.52 mmol) were used in General Procedure A to produce **SI-8** (0.061 g, 61%). $^{1}$H NMR (500 MHz, $CDCl_3$) δ 5.40 (d, *J* = 7.2 Hz, 1H), 4.24–4.16 (m, 1H), 3.20 – 3.13 (m, 1H), 3.01 (dd, *J* = 11.9, 4.6 Hz, 1H), 2.73 (dd, *J* = 11.9, 3.7 Hz, 1H), 1.47 (s, 9H), 1.45 (s, 8H), 1.45 (s, 8H), 1.22 (d, *J* = 7.0 Hz, 3H). $^{13}$C NMR (126 MHz, $CDCl_3$) δ 174.83, 170.96, 155.80, 82.05, 81.13, 79.75, 57.41, 54.34, 49.07, 28.48, 28.22, 28.15, 19.17. HRMS (MALDI) Calculated for $C_{19}H_{37}N_2O_6$ 389.2646, found 389.2640 $[M+H]^+$.

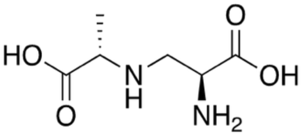


**SI-9**

**SI-8** was used to produce **SI-9** using General Procedure B. $^{1}$H NMR (500 MHz, $D_2O$ + 0.1% $CH_3OH$) δ 4.11 (td, *J* = 8.5, 5.8 Hz, 1H), 3.84 (q, *J* = 7.2 Hz, 1H), 3.56 (qd, *J* = 13.0, 7.3 Hz, 2H), 3.38 (s, 2H), 1.57 (d, *J* = 7.2 Hz, 3H); $^{13}$C NMR (126 MHz, $D_2O$ + 0.1% $CH_3OH$) δ 174.23, 170.67, 58.47, 49.91, 45.17, 14.96; HRMS (MALDI) Calculated for $C_6H_{13}N_2O_4$ 177.0870, found 177.0882 $[M+H]^+$.

**Synthesis of D,D-PDP Tln5 diastereomer (SI-13)**

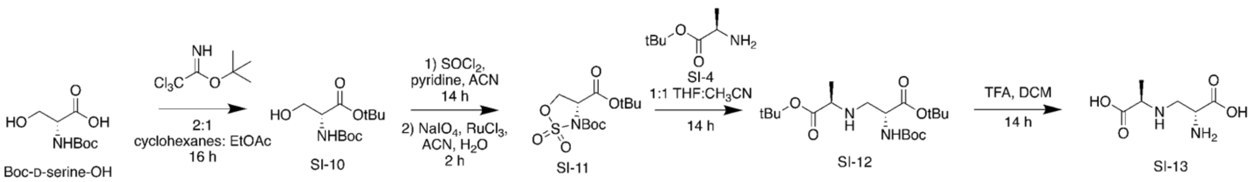


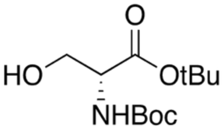

**SI-10**

This protocol was adapted from a literature reference.[51] Boc-D-Ser-OH (0.750 g, 3.65 mmol) was dissolved in 2:1 EtOAc: cyclohexane (38 mL) at room temperature. The tert-butyl ester reagent (2.62 mL, 14.7 mmol) was added and the reaction mixture stirred for 16 h, then concentrated in vacuo. The crude reaction mixture was purified via flash column chromatograph (3:2 hex: EtOAc eluent system) to obtain the final product as a white solid (0.965 g, 89%). $^1$H NMR (500 MHz, $CD_3OD$) $\delta$ 4.08 (d, $J$ = 4.0 Hz, 1H), 3.78 (qd, $J$ = 11.1, 4.5 Hz, 3H), 1.48 (s, 11H), 1.45 (s, 11H); $^{13}$C NMR (126 MHz, $CD_3OD$) $\delta$ 170.27, 164.57, 156.55, 92.56, 81.53, 79.28, 61.84, 56.64, 27.30, 26.89. HRMS (MALDI) Calculated for $C_{12}H_{24}NO_5Na$ 284.3168, found 284.3328 $[M+H]^+$.

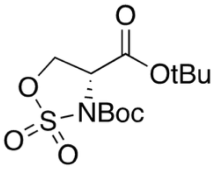

**SI-11**

The protocol was adapted from the following references.[51,52] To a stirred solution of $SOCl_2$ (0.76 mL, 10.52 mmol) in $CH_3CN$ (18 mL) at -40 °C, **SI-10** (1.103 g, 4.22 mmol) was added as a solution in dry $CH_3CN$ (44 mL) over the course of 20 min under Ar (g). After stirring for 1.5 h at −40 °C, dry pyridine (2.003 mL, 25.33 mmol) was added over 15 min. The reaction was slowly brought up to room temperature and stirred for an additional 16 h, before quenching the reaction by adding ice water (14 mL). The aqueous layer was extracted with EtOAc (2 x 70 mL), the organic layers were pooled, washed with water (20 mL), brine (27 mL), dried over $Na_2SO_4$, filtered, then dried in vacuo to yield the crude reaction mixture as an oil. The crude mixture was taken into the next step, where the mixture was dissolved in 1:1 $CH_3CN$: $H_2O$ (36 mL), and $RuCl_3$ (0.046 g, 0.022 mmol) and $NaIO_4$ (1.678 g, 8.48 mmol) were added sequentially. The reaction stirred at 0 °C for 2.5 h, then diluted with EtOAc (44 mL). The aqueous layer was extracted with EtOAc (2 x 28 mL), then the organic layers were pooled, washed with brine (41 mL), dried over $Na_2SO_4$, then concentrated in vacuo. The crude reaction mixture was purified via flash column chromatography using a 4:1 hexane: EtOAc eluent system to yield the final product as a white solid (0.716 g, 52%). $^1$H NMR (500 MHz, $CDCl_3$) $\delta$ 4.73 (t, $J$ = 8.3 Hz, 1H), 4.63 (d, $J$ = 8.7 Hz, 2H), 1.56 (s, 11H), 1.51 (s, 9H). $^{13}$C NMR (126 MHz, $CDCl_3$) $\delta$ 166.01, 148.10, 85.87, 84.49, 67.75, 58.13, 27.89, 27.82. HRMS (MALDI) Calculated for $C_{12}H_{21}NO_7SNa$ 346.0931, found 346.0931 $[M+Na]^+$.

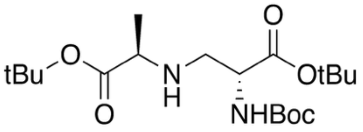

**SI-12**

**SI-10** (0.061 g, 0.19 mmol) and **SI-4** (0.055 g, 0.38 mmol) were used in General Procedure A to produce **SI-8** (0.063 g, 43%). $^{1}H$ NMR (500 MHz, $CDCl_3$) δ 5.40 (d, *J* = 7.2 Hz, 1H), 4.24–4.15 (m, 1H), 3.16 (q, *J* = 6.6 Hz, 1H), 3.01 (dd, *J* = 11.9, 4.6 Hz, 1H), 2.72 (dd, *J* = 11.8, 3.7 Hz, 1H), 1.47 (s, 8H), 1.45 (s, 8H), 1.45 (s, 8H), 1.22 (d, *J* = 7.0 Hz, 3H). $^{13}C$ NMR (126 MHz, $CDCl_3$) δ 174.84, 170.96, 57.41, 54.34, 49.07, 28.48, 28.22, 28.15, 19.17. HRMS (MALDI) Calculated for $C_{19}H_{37}N_2O_6$ 389.2646, found 389.2654 $[M+H]^+$.

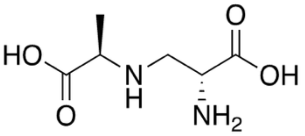

**SI-13**

**SI-12** was used to produce **SI-13** using General Procedure B. $^{1}H$ NMR (500 MHz, $D_2O$ + 0.1% $CH_3OH$) δ 4.11 (t, *J* = 7.1 Hz, 1H), 3.83 (q, *J* = 7.1 Hz, 1H), 3.54 (qd, *J* = 13.0, 7.3 Hz, 2H), 1.56 (d, *J* = 7.2 Hz, 3H); $^{13}C$ NMR (126 MHz, $D_2O$ + 0.1% $CH_3OH$) δ 174.37, 170.78, 58.49, 50.00, 45.24, 15.01. HRMS (MALDI) Calculated for $C_6H_{13}N_2O_4$ 177.0870, found 177.0881 $[M+H]^+$.

**Synthesis of L,D-PDP Tln5 diastereomer (SI-15)**

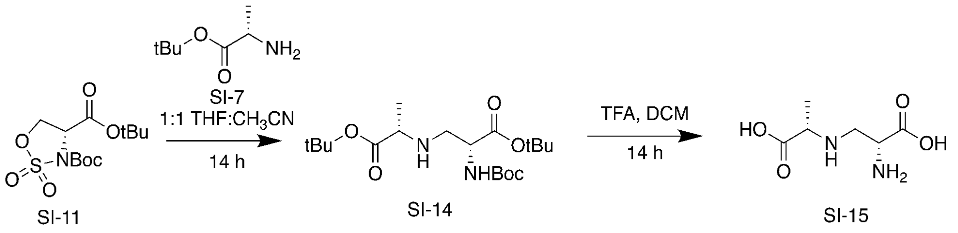


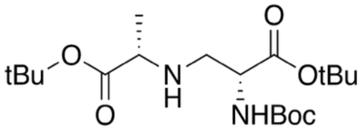

**SI-14**

**SI-11** (0.061 g, 0.189 mmol) and **SI-7** (0.055 g, 0.379 mmol) were used in General Procedure A to produce **SI-14** (0.043 g, 29%). $^{1}H$ NMR (500 MHz, $CDCl_3$) δ 5.39 (d, *J* = 5.9 Hz, 1H), 4.22–4.14 (m, 1H), 3.16 (q, *J* = 6.9 Hz, 1H), 2.99 (dd, *J* = 12.3, 4.1 Hz, 1H), 2.83 (dd, *J* = 12.2, 3.7 Hz, 1H), 1.47 (s, 9H), 1.45 (s, 9H), 1.44 (s, 8H), 1.20 (d, *J* = 7.0 Hz, 3H); $^{13}C$ NMR (126 MHz, $CDCl_3$) δ 174.89, 170.97, 57.57, 54.66, 49.14, 28.49, 28.21, 28.16, 19.13; HRMS (MALDI) Calculated for $C_{19}H_{37}N_2O_6$ 389.2646, found 389.2632 $[M+H]^+$.

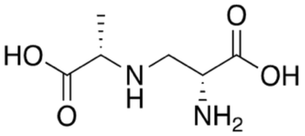

**SI-15**

**SI-14** was used to produce **SI-15** using General Procedure B. $^{1}H$ NMR (500 MHz, $D_2O$ + 0.1% $CH_3OH$) δ 4.10 (dd, *J* = 8.8, 5.5 Hz, 1H), 3.90–3.80 (m, 1H), 3.54 (ddt, *J* = 34.3, 12.8, 7.5 Hz, 2H), 1.56 (d, *J* = 7.2 Hz, 3H). $^{13}C$ NMR (126 MHz, $D_2O$ + 0.1% $CH_3OH$) δ 174.32, 170.97, 58.43, 49.60, 45.03, 14.87. HRMS (MALDI) Calculated for $C_6H_{13}N_2O_4$ 177.0870, found 177.0879 $[M+H]^+$.

Supplemental Information: **Integrated metabolomic and genomic insights into amino acid incorporation within the hybrid polyketide-alkaloid antibiotic TLN-05220**

### NMR and Compound Characterization

## SI-1:

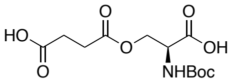


**SI-1**

500 MHz,
$^{1}H$ in $CD_3OD$

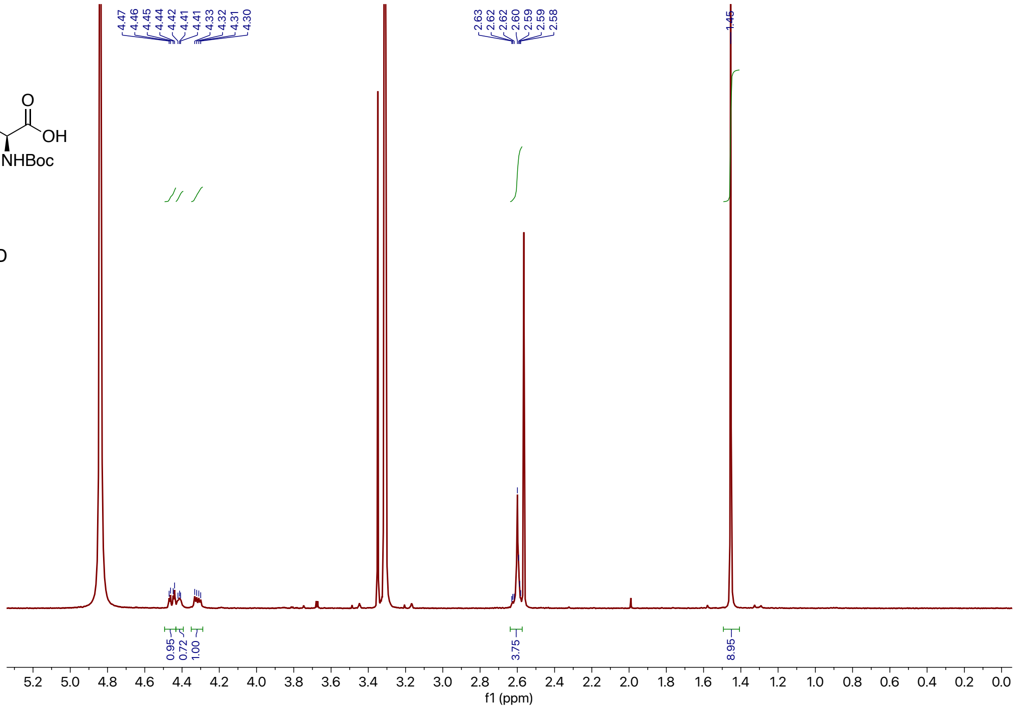

## SI-2:

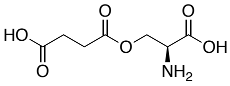


**SI-2**

500 MHz,
$^{1}H$ in $D_2O$
+ 0.1% MeOH

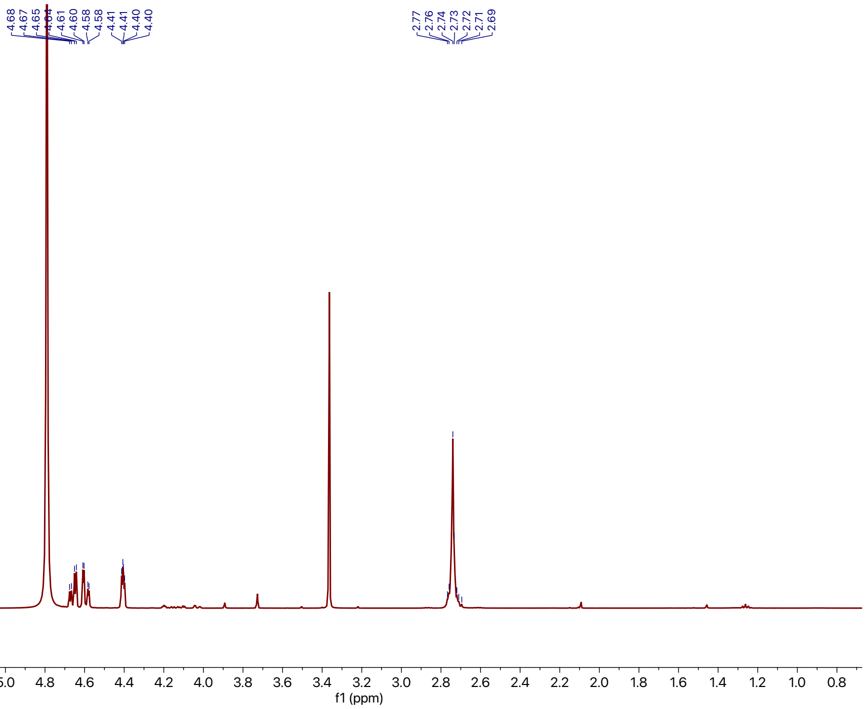


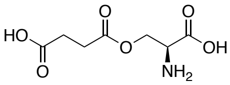


**SI-2**

125 MHz,
$^{13}C$ in $D_2O$
+ 0.1% MeOH

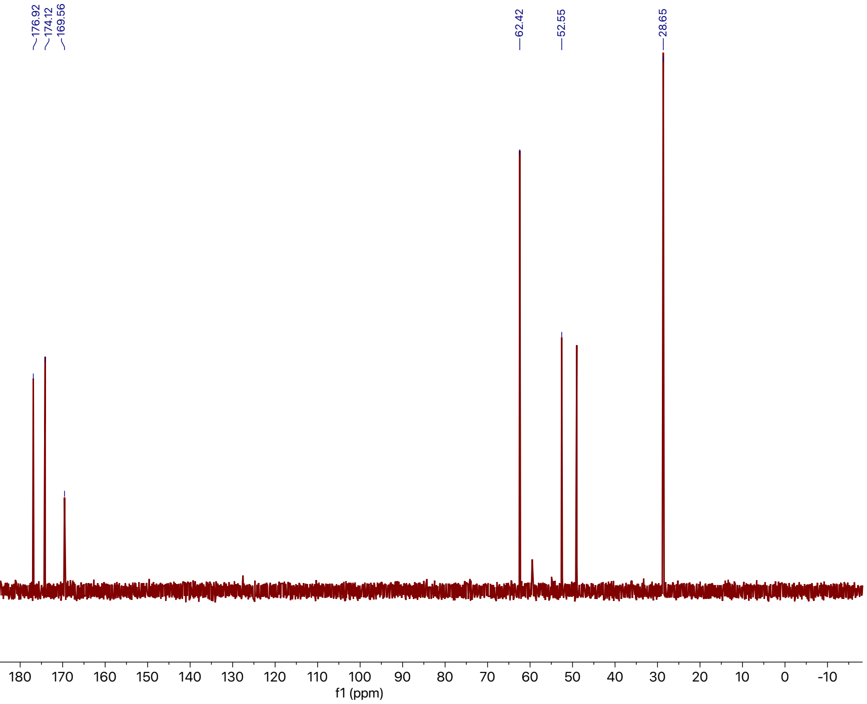

Supplemental Information: **Integrated metabolomic and genomic insights into amino acid incorporation within the hybrid polyketide-alkaloid antibiotic TLN-05220**

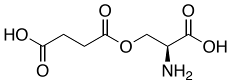

**SI-2**

500 MHz,
COSY in $D_2O$
+ 0.1% MeOH

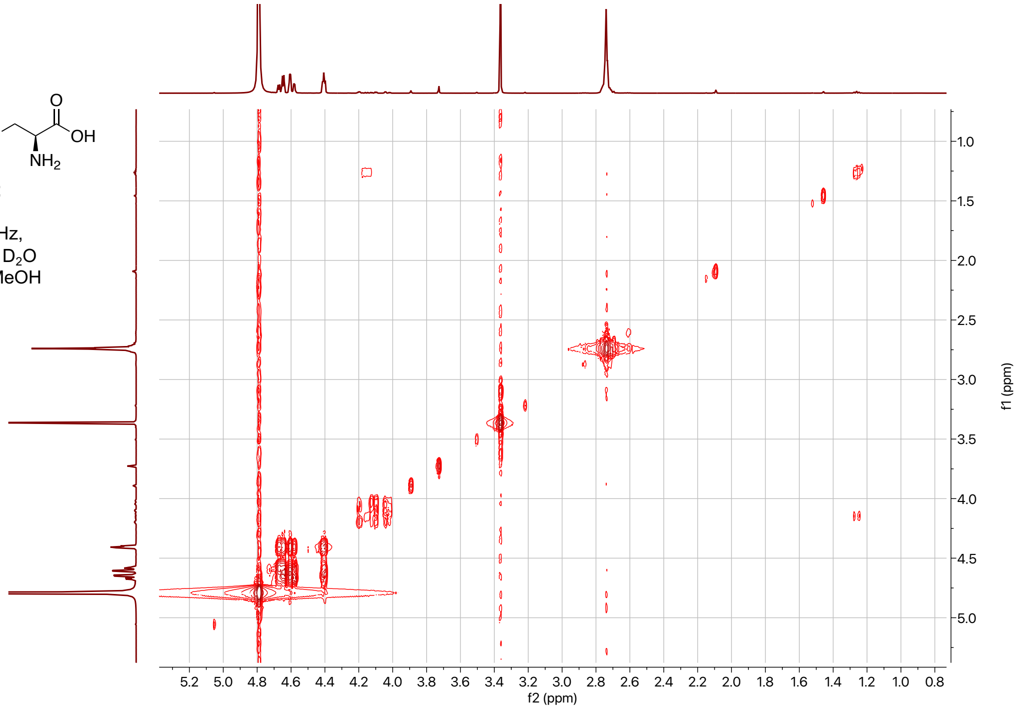


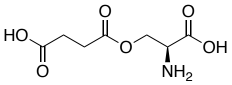

**SI-2**

500 MHz,
HSQC in $D_2O$
+ 0.1% MeOH

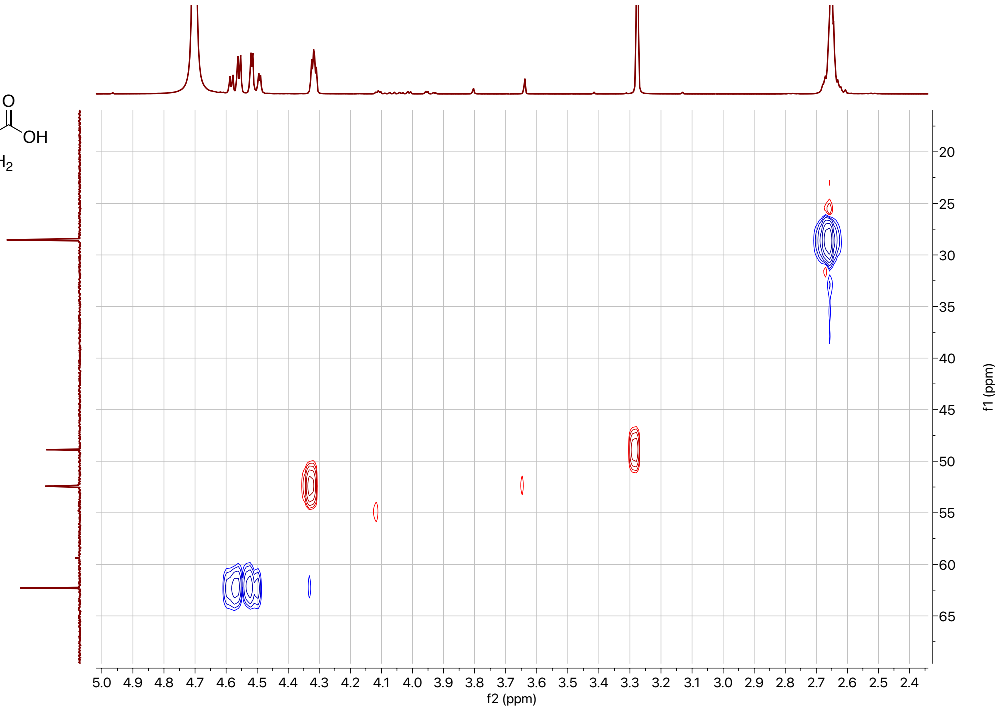

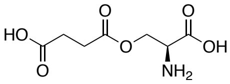


**SI-2**

500 MHz,
HMBC in $D_2O$
+ 0.1% MeOH

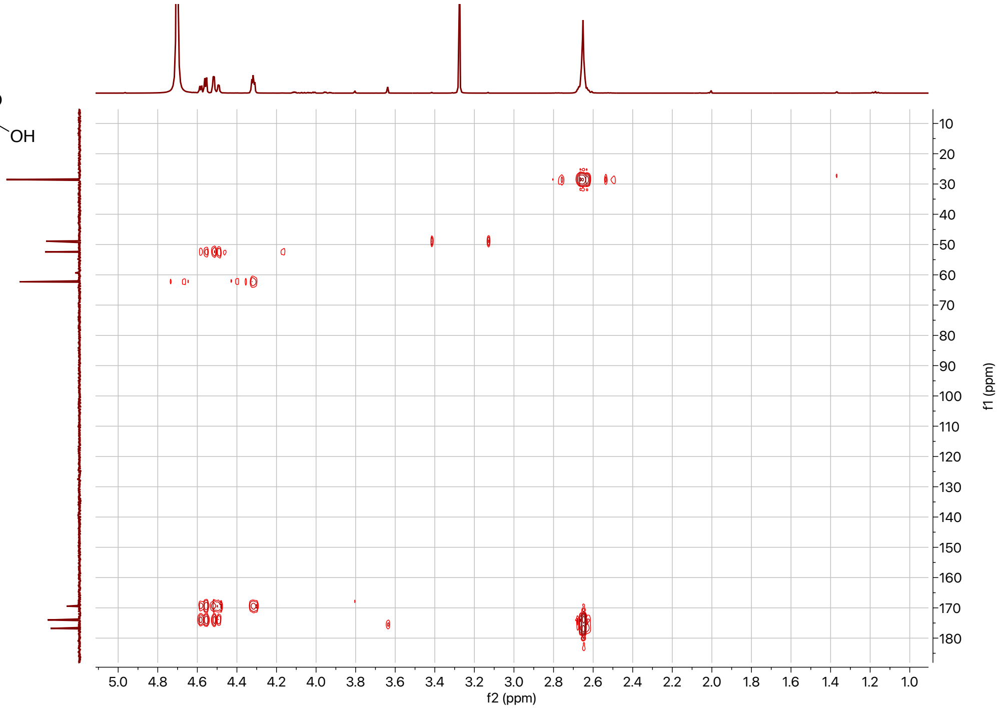

**SI-3:**

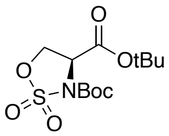


**SI-3**

500 MHz,
$^{1}H$ in $CDCl_3$

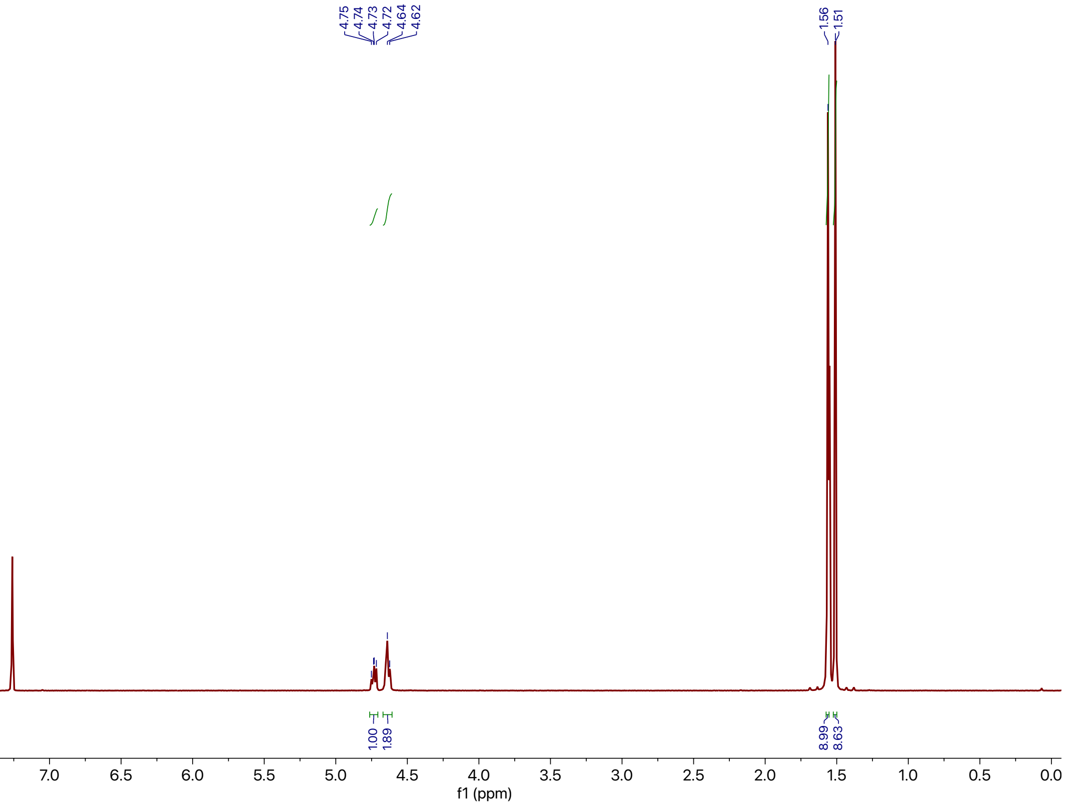

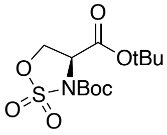


**SI-3**

125 MHz,
$^{13}C$ in $CDCl_3$

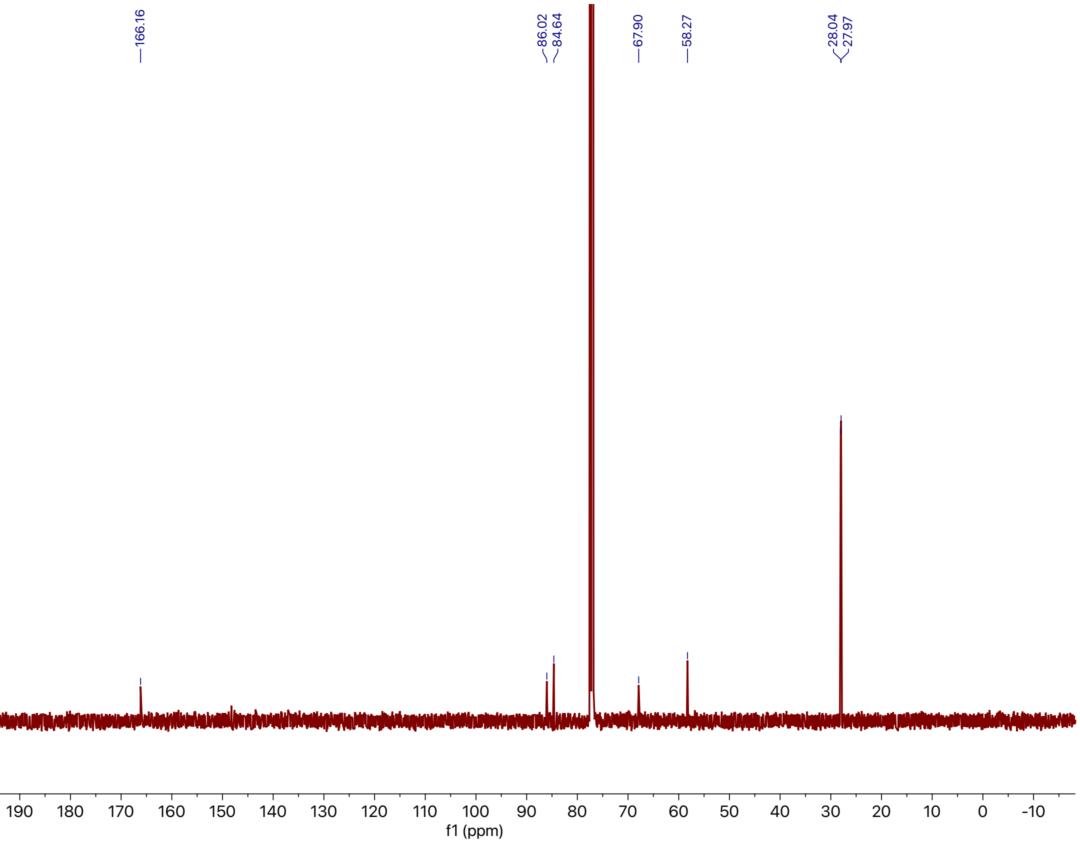


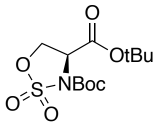


**SI-3**

500 MHz,
COSY in $CDCl_3$

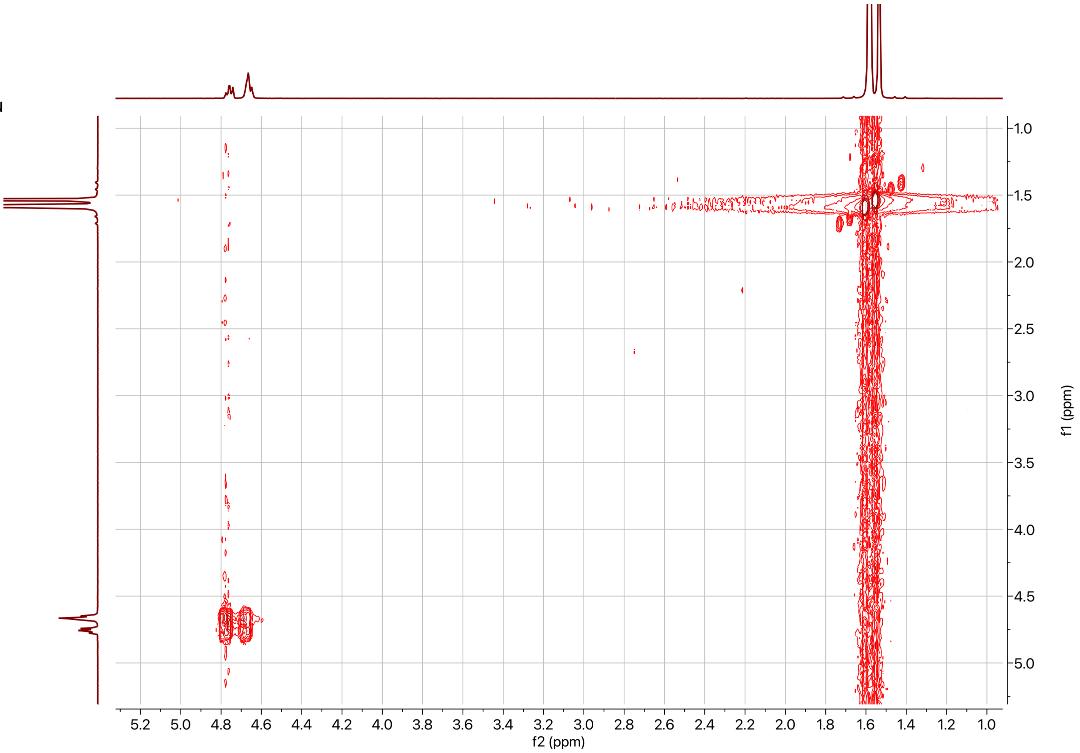

Supplemental Information: **Integrated metabolomic and genomic insights into amino acid incorporation within the hybrid polyketide-alkaloid antibiotic TLN-05220**

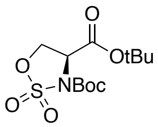


**SI-3**

500 MHz,
HSQC in $CDCl_3$

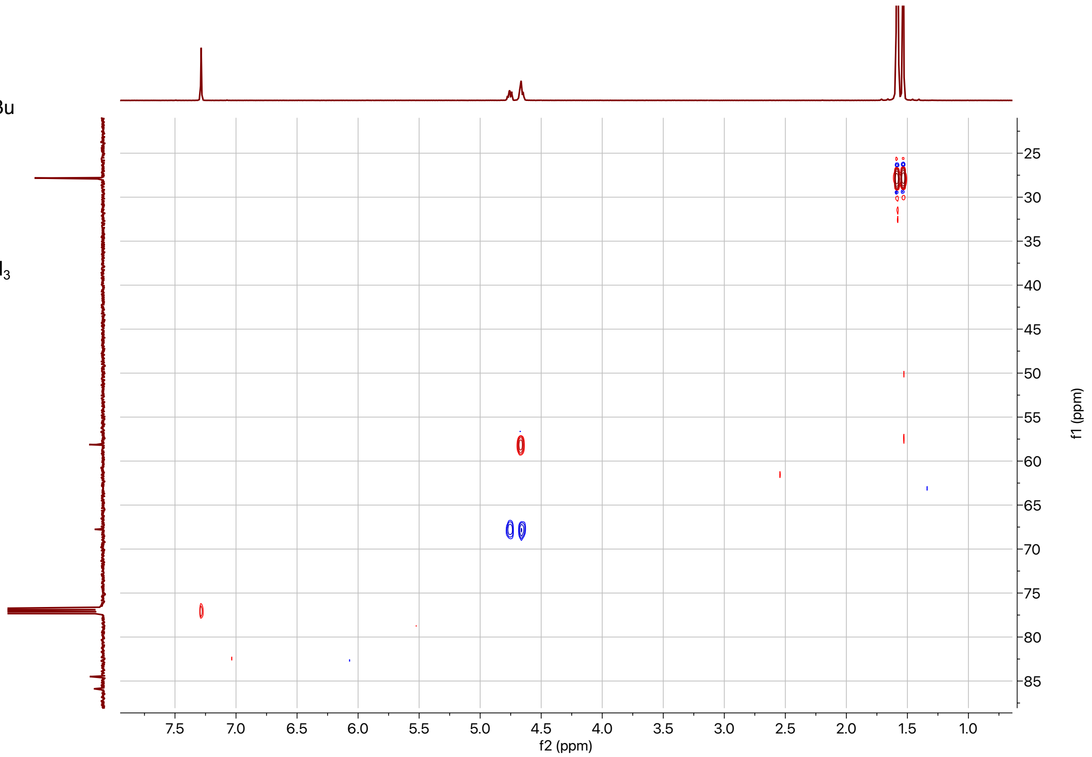


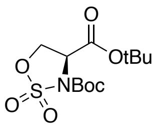


**SI-3**

500 MHz,
HMBC in $CDCl_3$

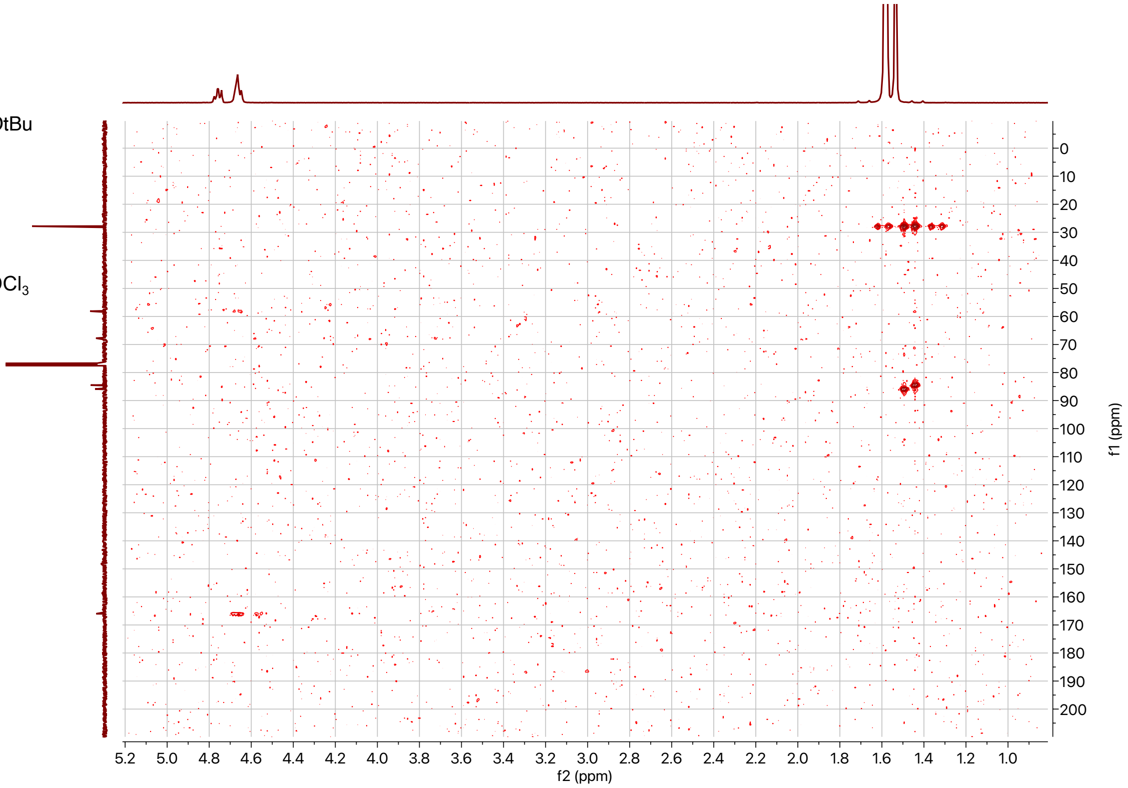

Supplemental Information: **Integrated metabolomic and genomic insights into amino acid incorporation within the hybrid polyketide-alkaloid antibiotic TLN-05220**

## SI-5:

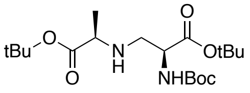


**SI-5**

500 MHz,
$^{1}$H in $CDCl_3$

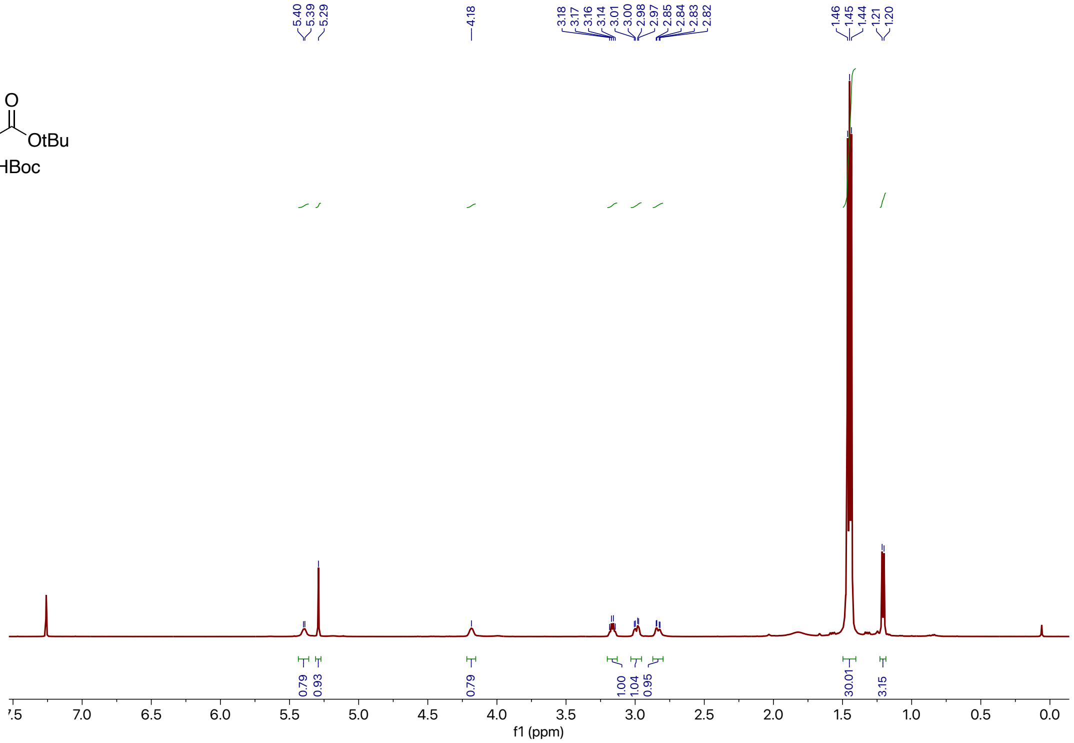


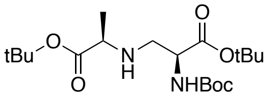


**SI-5**

125 MHz,
$^{13}$C in $CDCl_3$

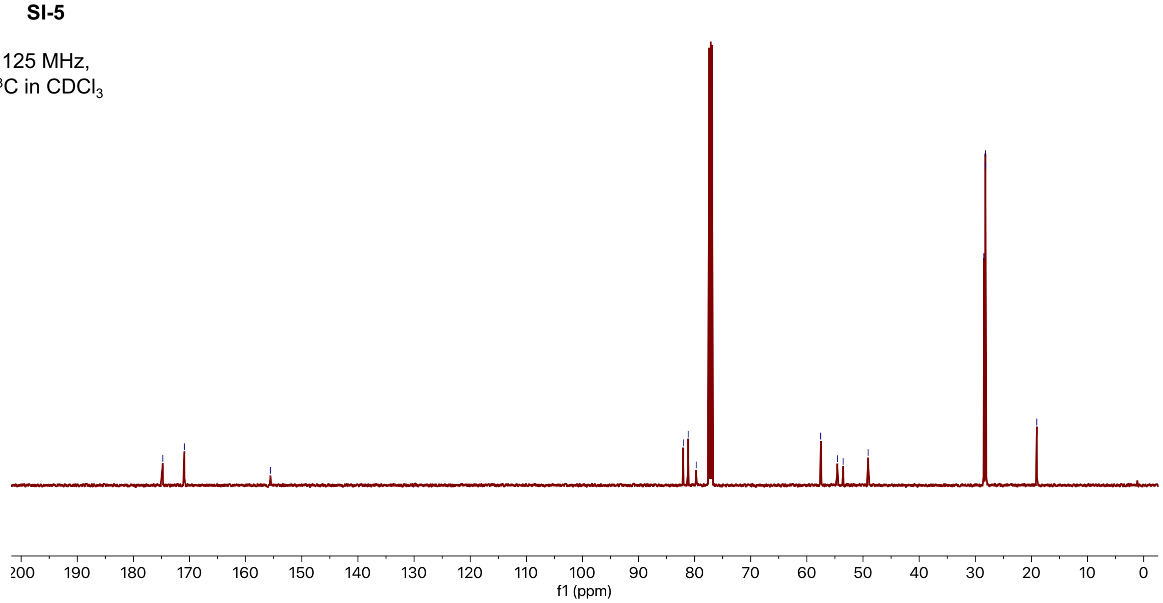

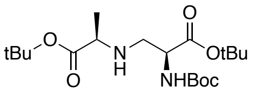


**SI-5**

500 MHz,
COSY in $CDCl_3$

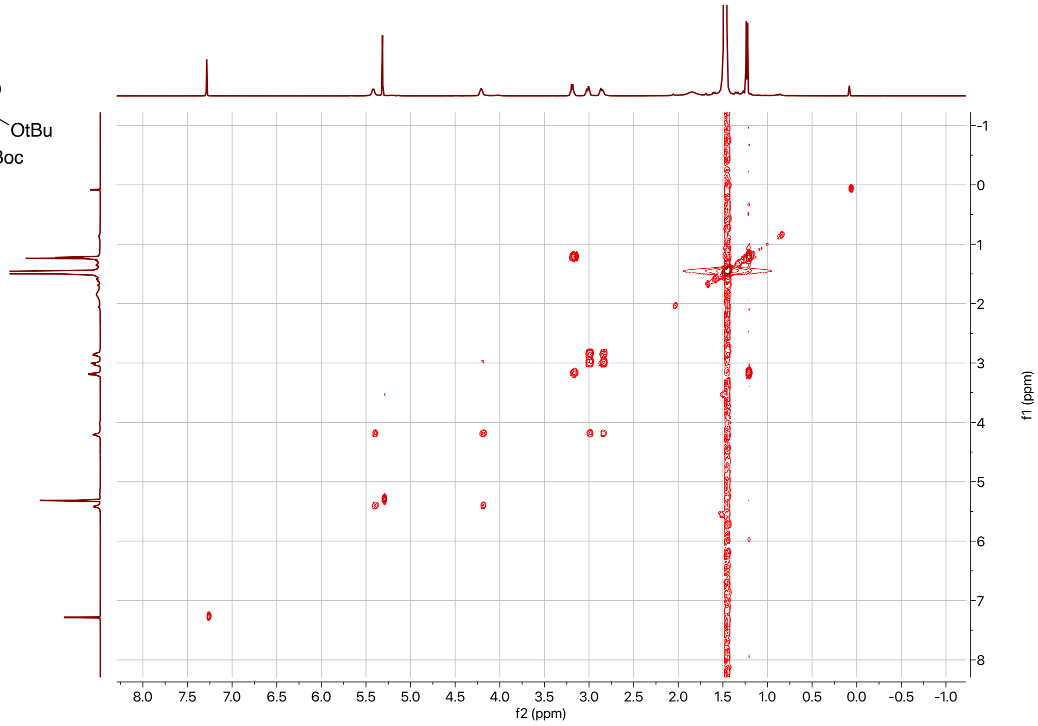


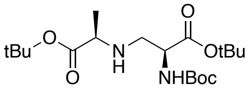


**SI-5**

500 MHz,
HSQC in $CDCl_3$

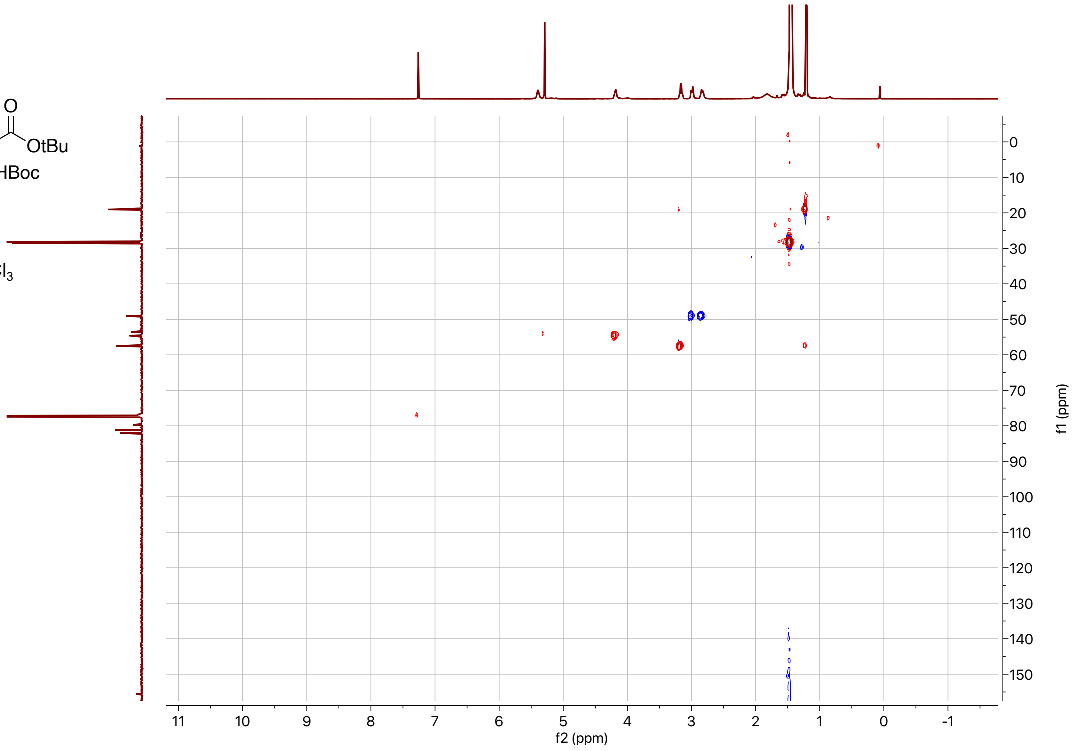

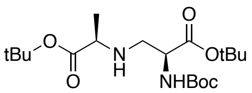


**SI-5**

500 MHz,
HMBC in $CDCl_3$

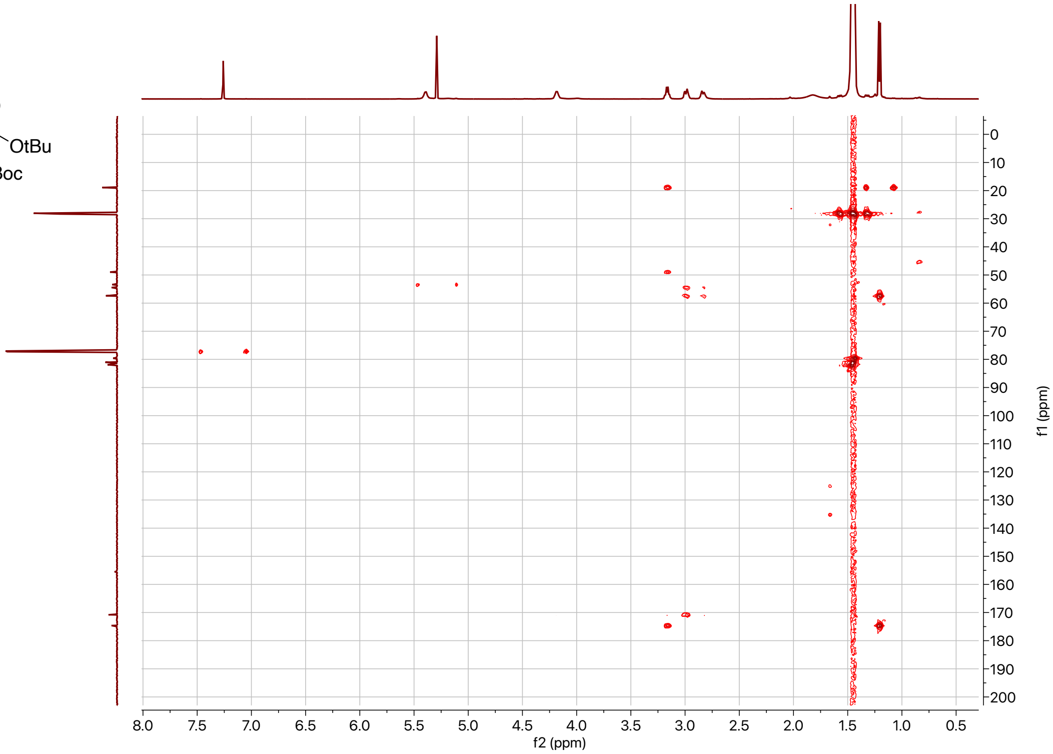

## SI-6:

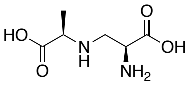


**SI-6**

500 MHz,
$^{1}H$ in $D_2O$
+ 0.1% MeOH

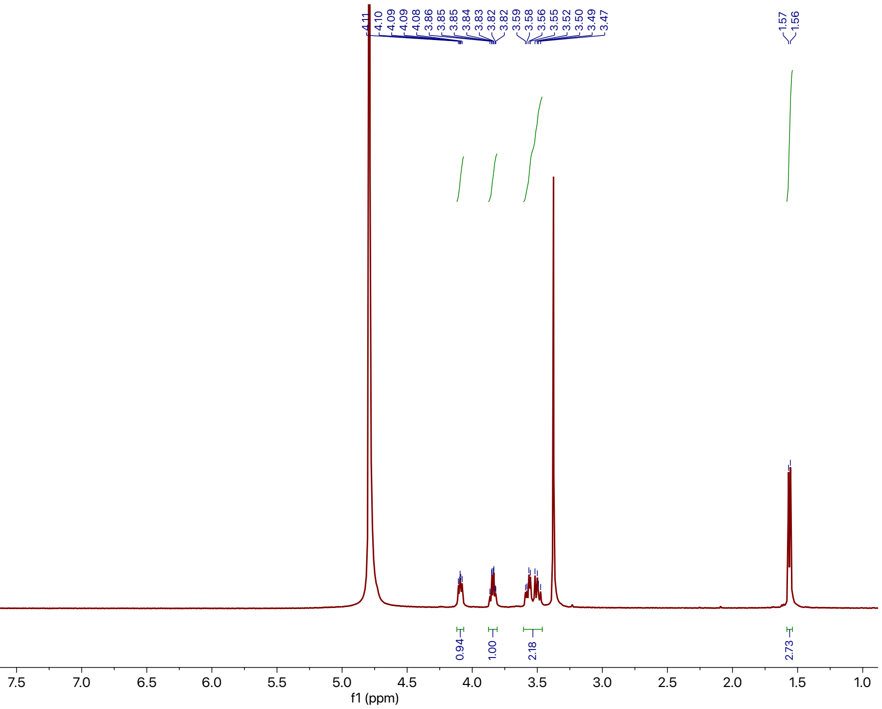


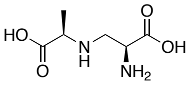


**SI-6**

125 MHz,
$^{1}H$ in $D_2O$
+ 0.1% MeOH

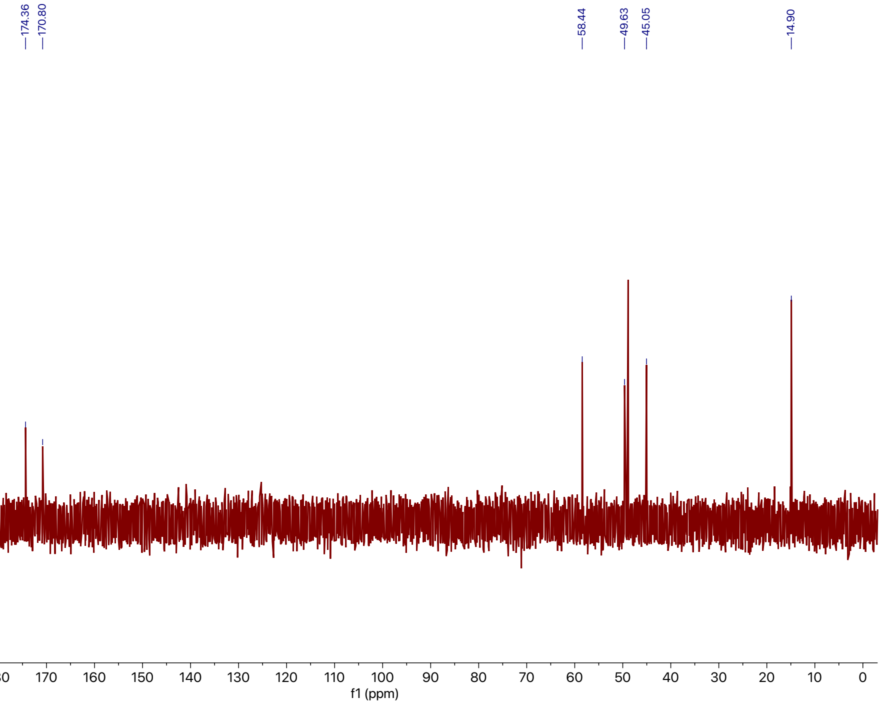

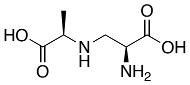

**SI-6**

500 MHz,
COSY in $D_2O$
+ 0.1% MeOH

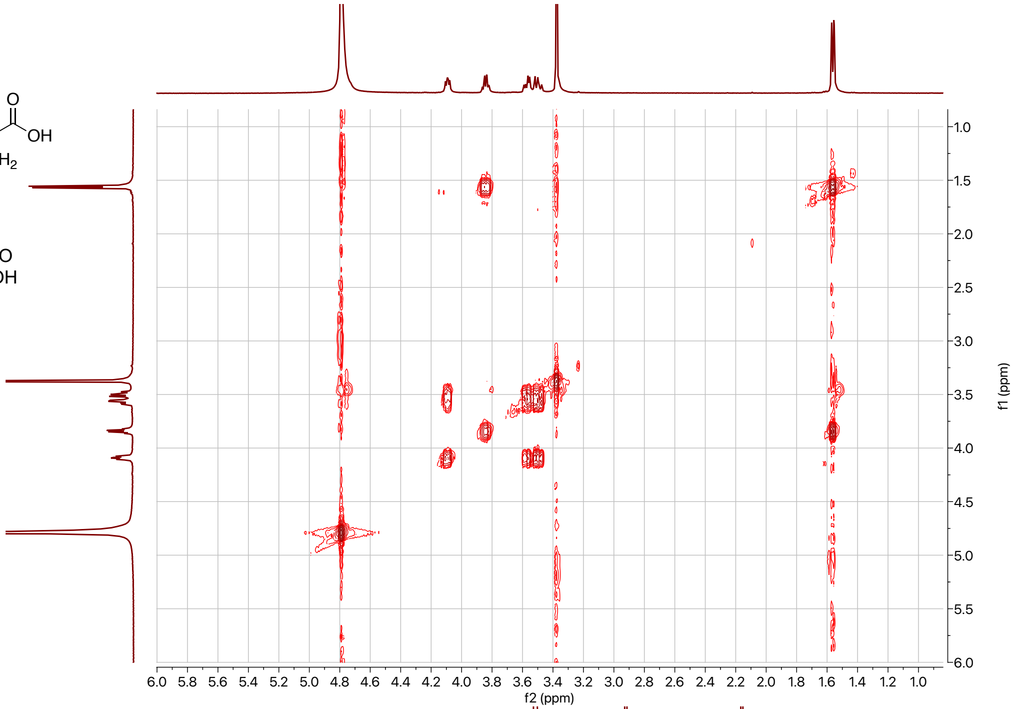

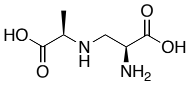

**SI-6**

500 MHz,
HSQC in $D_2O$
+ 0.1% MeOH

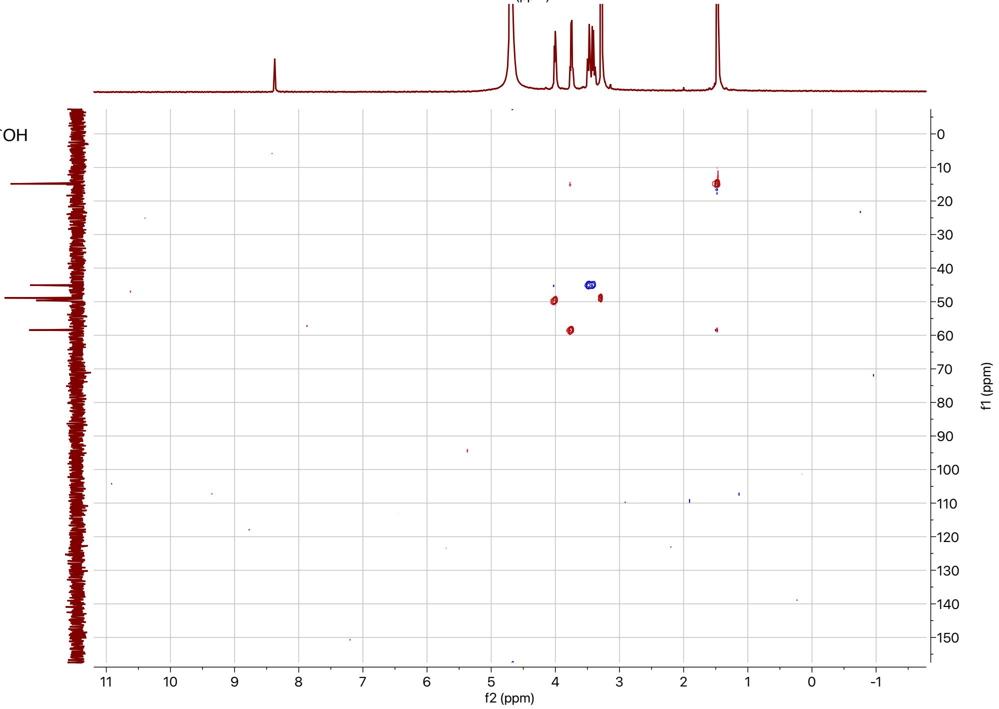

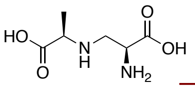

**SI-6**

500 MHz,
HMBC in $D_2O$
+ 0.1% MeOH

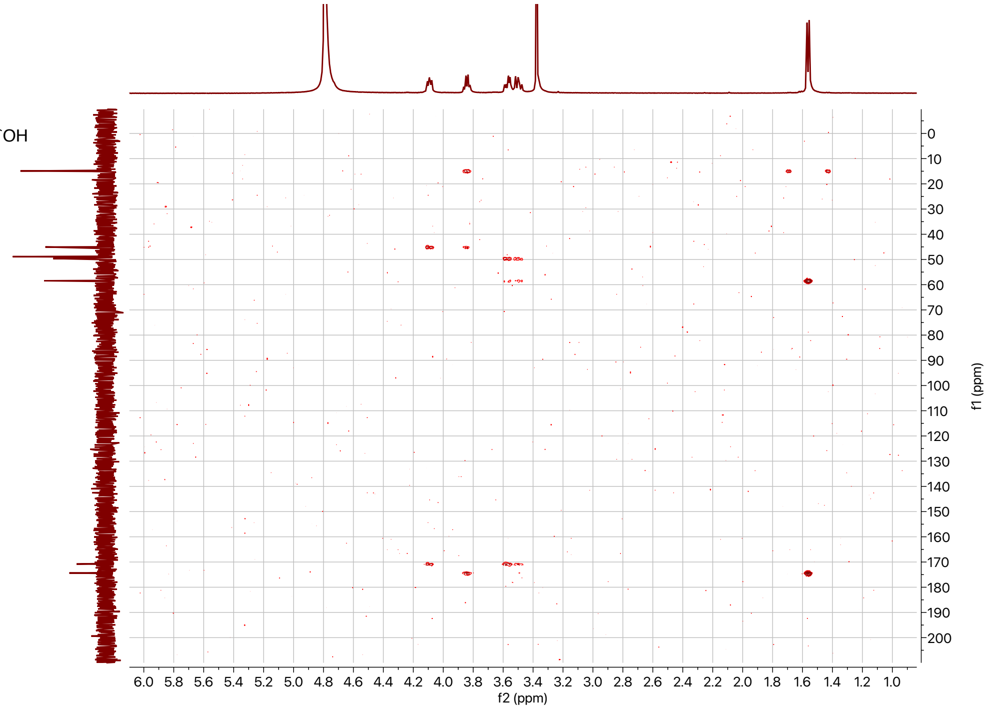

## SI-8:

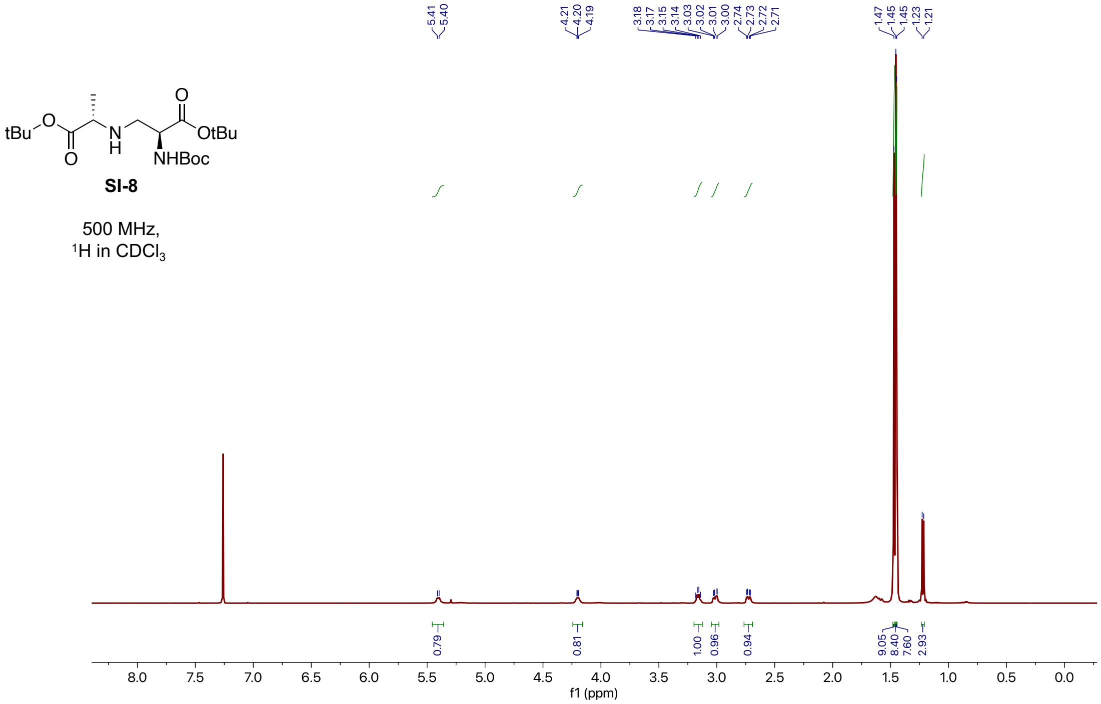


**SI-8**

500 MHz,
$^{1}H$ in $CDCl_3$

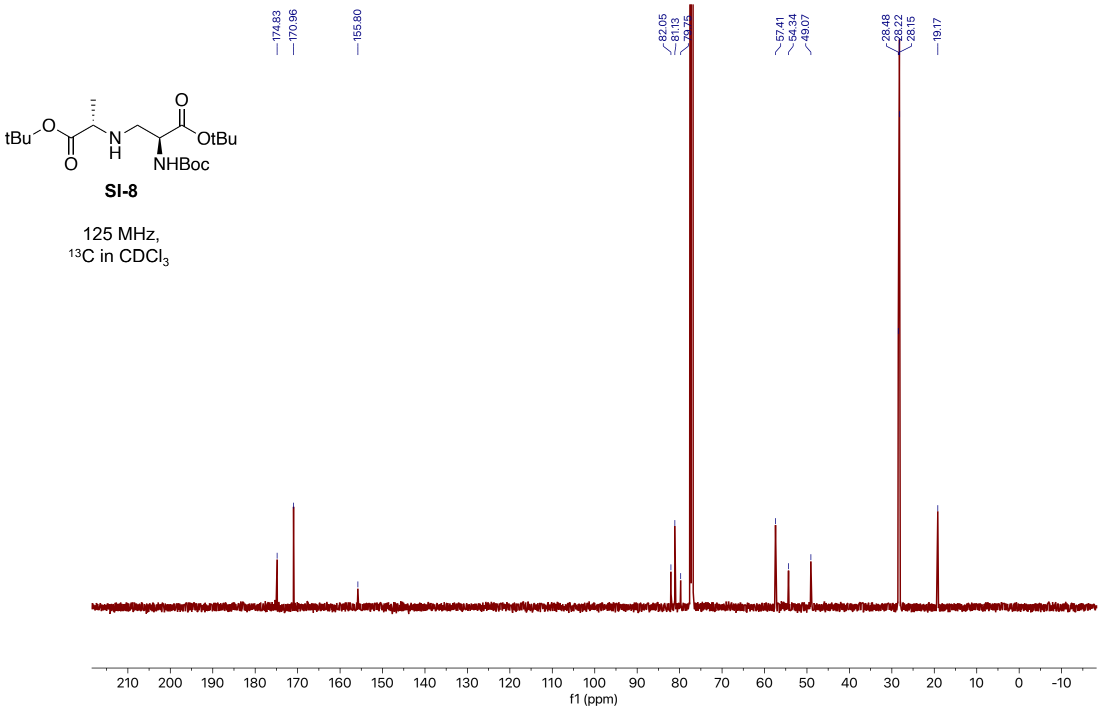


**SI-8**

125 MHz,
$^{13}C$ in $CDCl_3$

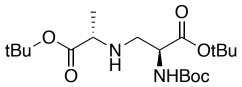


**SI-8**

500 MHz,
COSY in $CDCl_3$

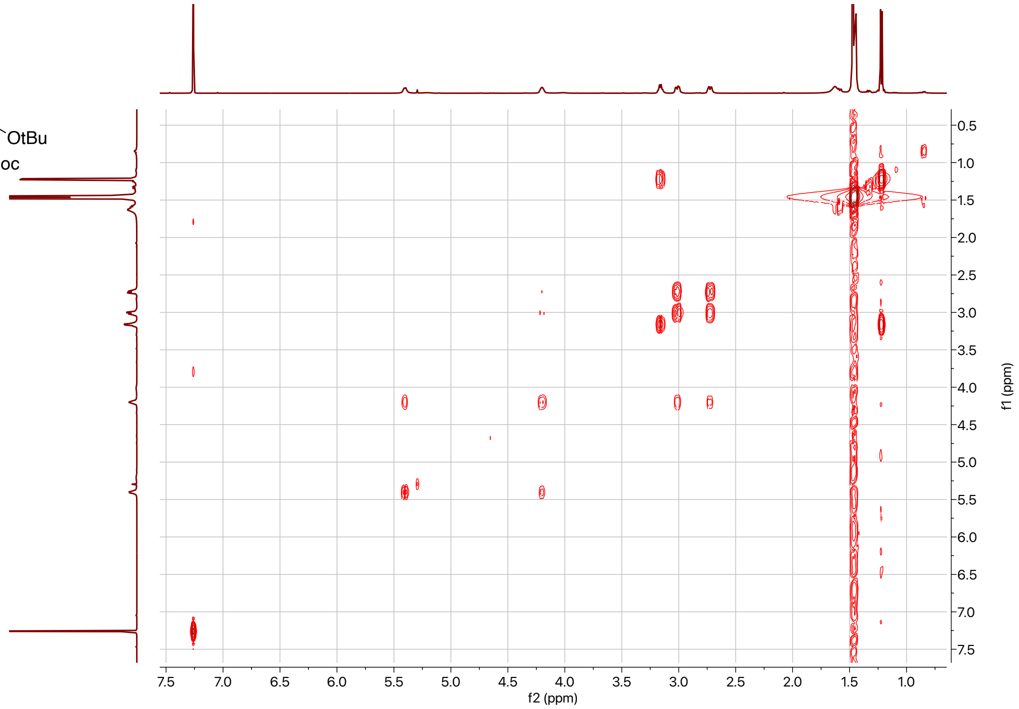


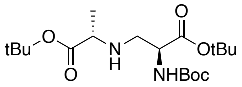


**SI-8**

500 MHz,
HSQC in $CDCl_3$

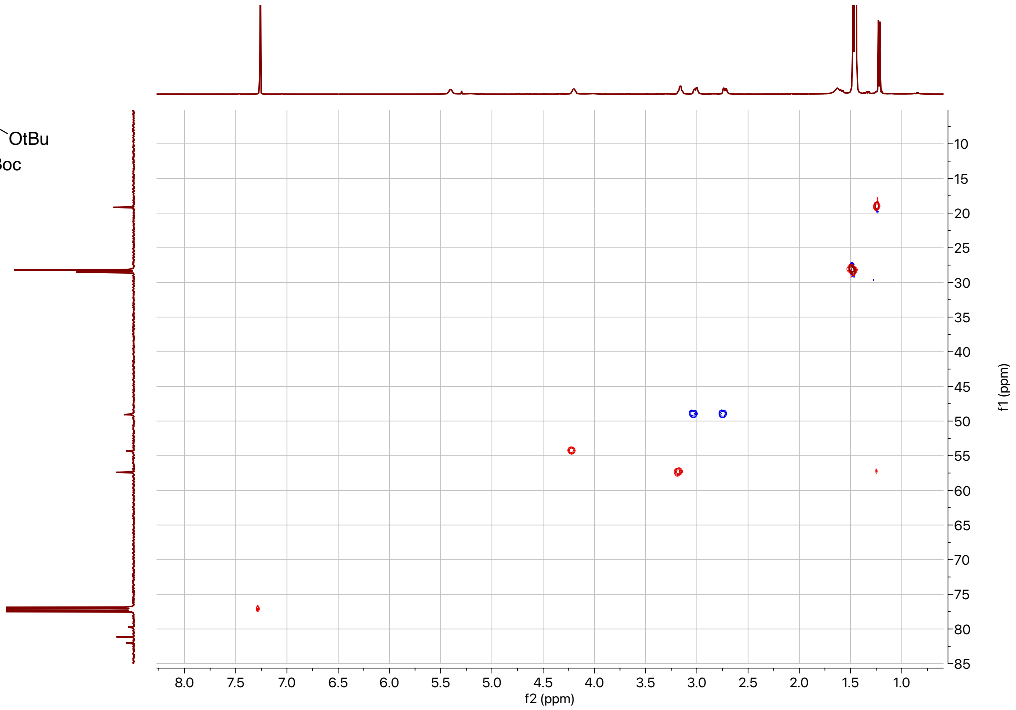

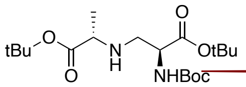


**SI-8**

500 MHz,
HMBC in $CDCl_3$

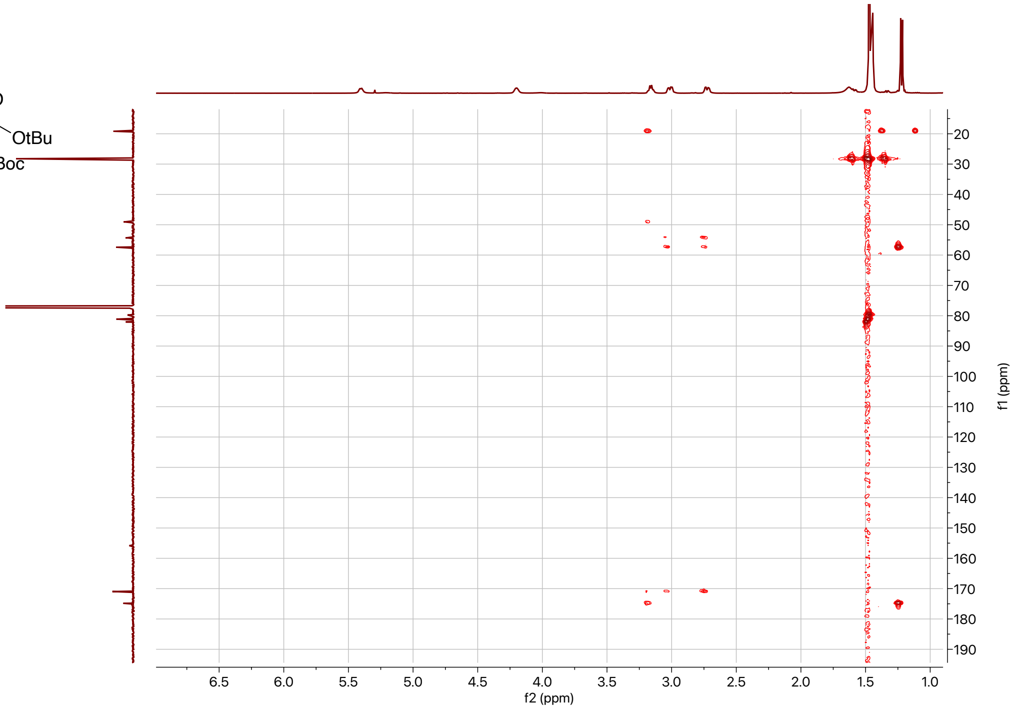

## SI-9:

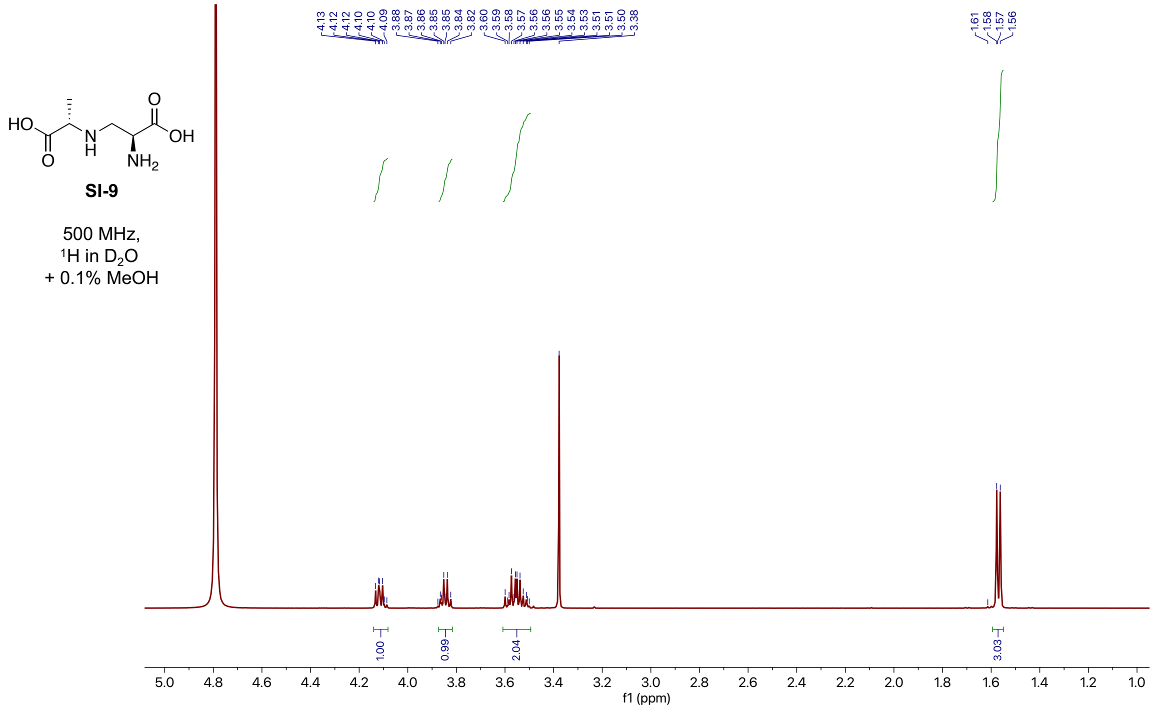
SI-9
500 MHz,
$^{1}H$ in $D_2O$
+ 0.1% MeOH
1.00
0.99
2.04
3.03
f1 (ppm)

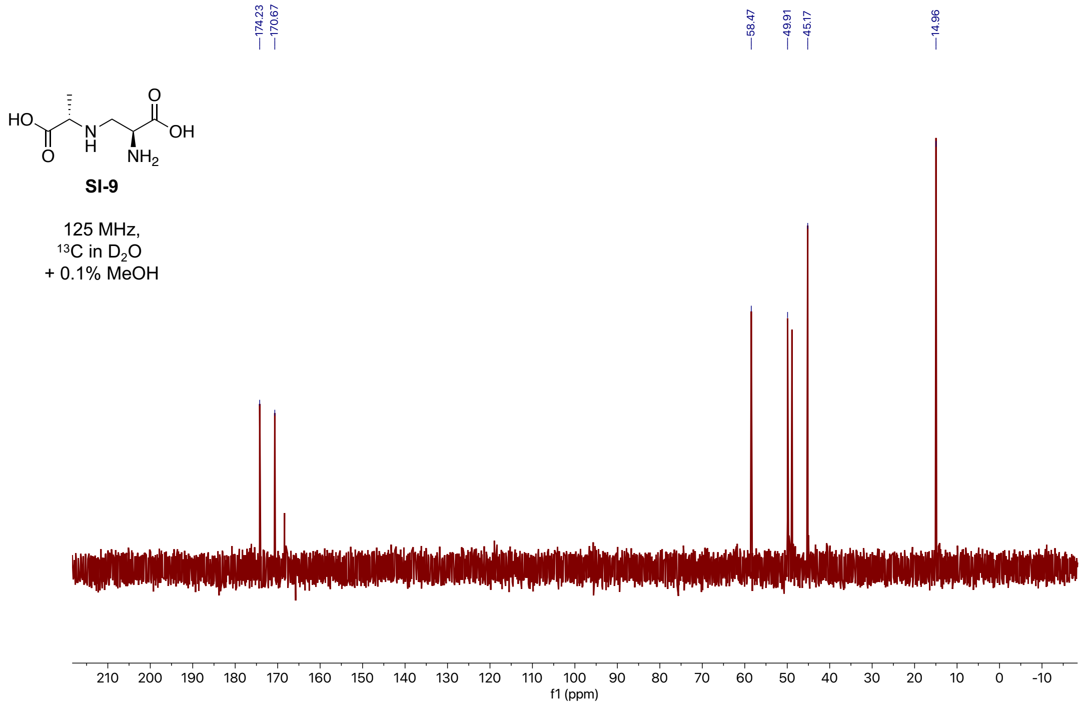
SI-9
125 MHz,
$^{13}C$ in $D_2O$
+ 0.1% MeOH
174.23
170.67
58.47
49.91
45.17
14.96
f1 (ppm)

Supplemental Information: **Integrated metabolomic and genomic insights into amino acid incorporation within the hybrid polyketide-alkaloid antibiotic TLN-05220**

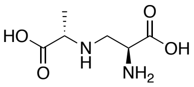

**SI-9**

500 MHz,
COSY in $D_2O$
+ 0.1% MeOH

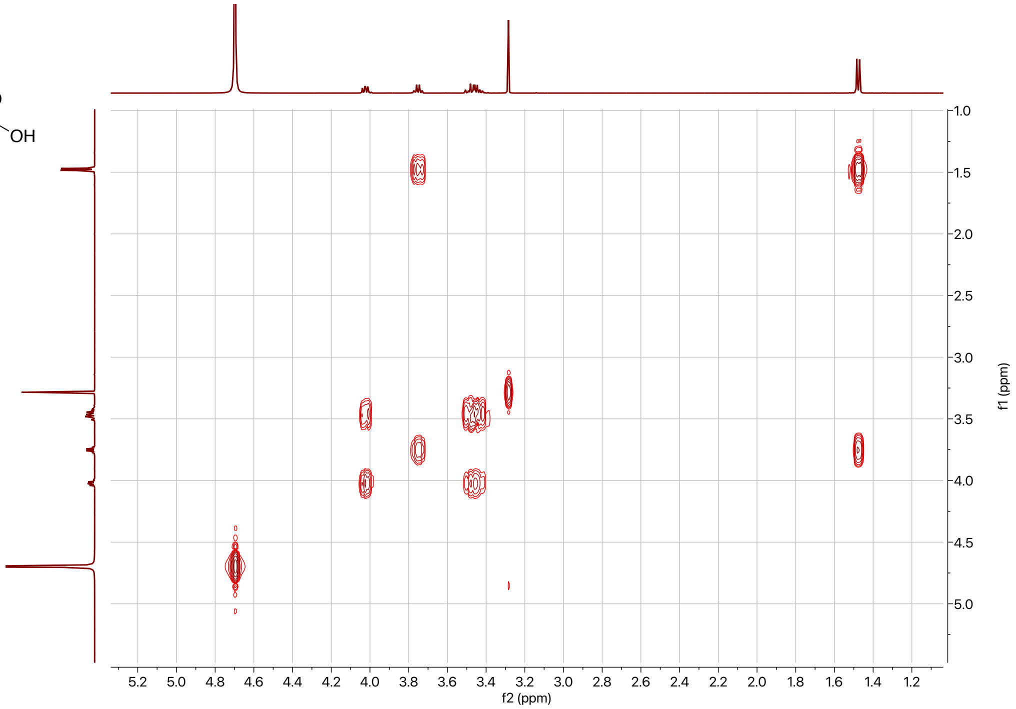


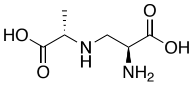

**SI-9**

500 MHz,
HSQC in $D_2O$
+ 0.1% MeOH

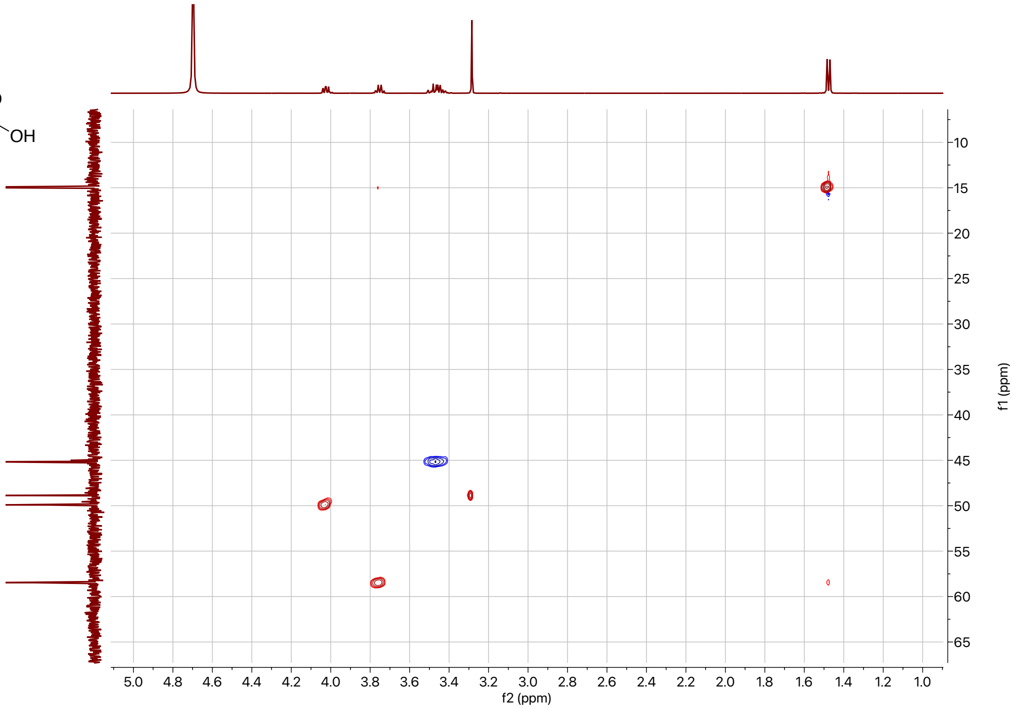

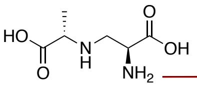

**SI-9**

500 MHz,
HMBC in $D_2O$
+ 0.1% MeOH

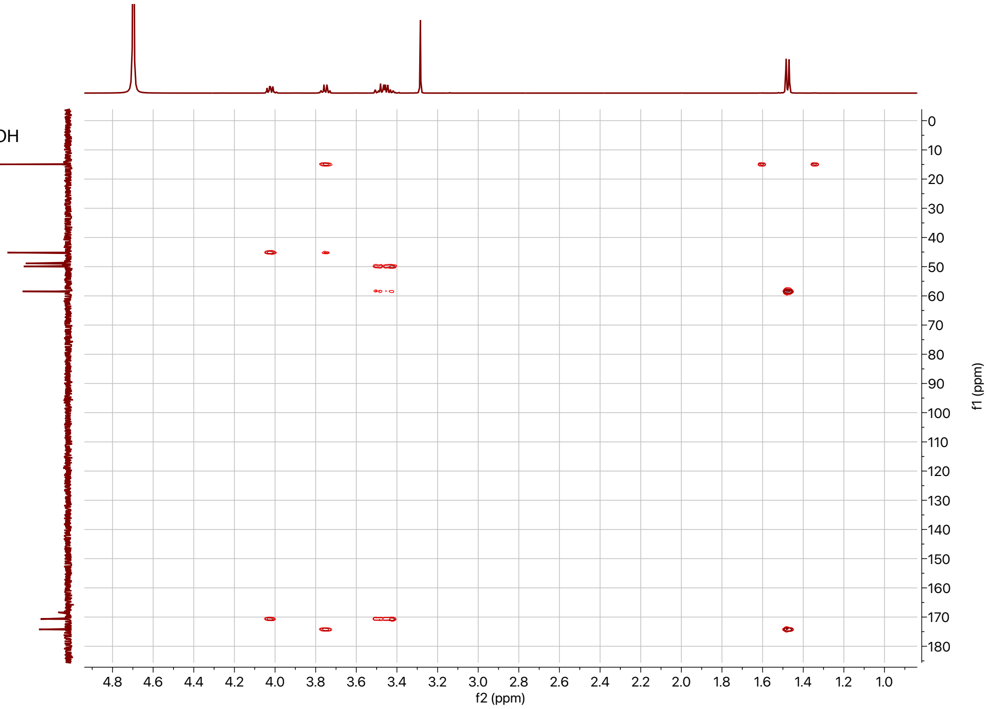

Supplemental Information: **Integrated metabolomic and genomic insights into amino acid incorporation within the hybrid polyketide-alkaloid antibiotic TLN-05220**

## SI-10:

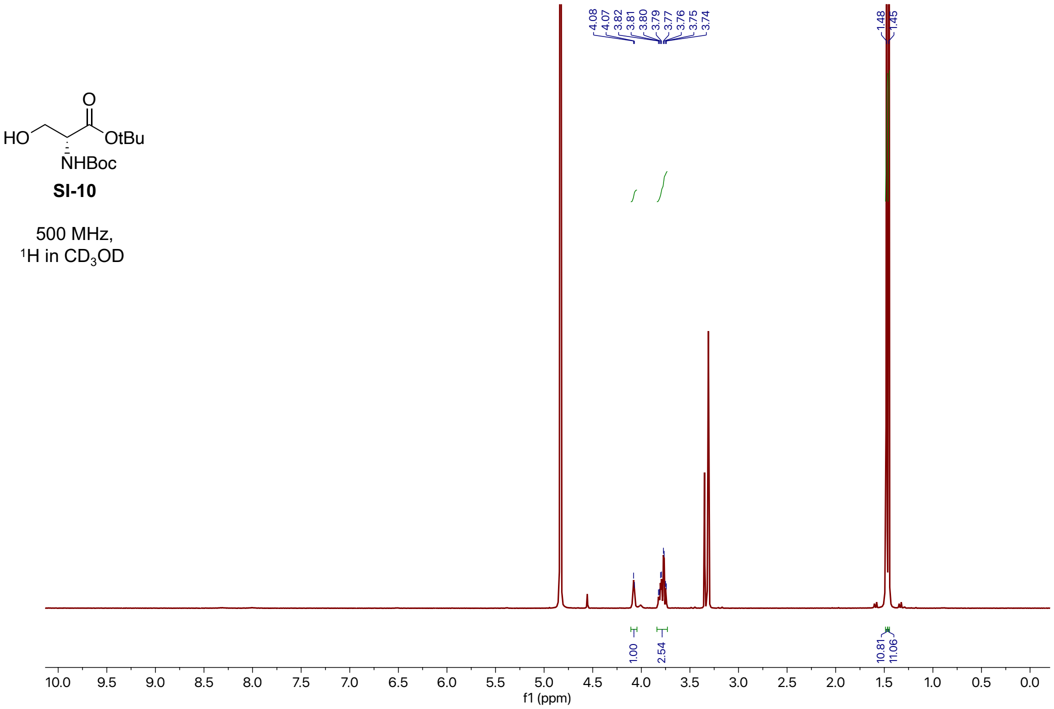


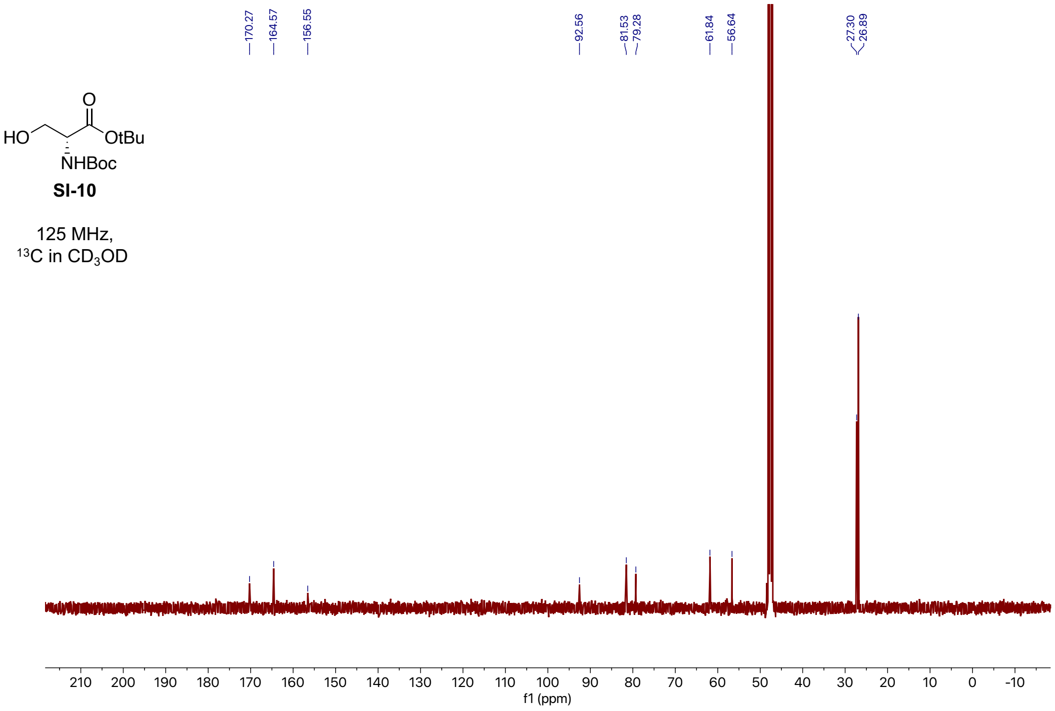

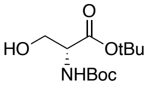


**SI-10**

500 MHz,
COSY in $CD_3OD$

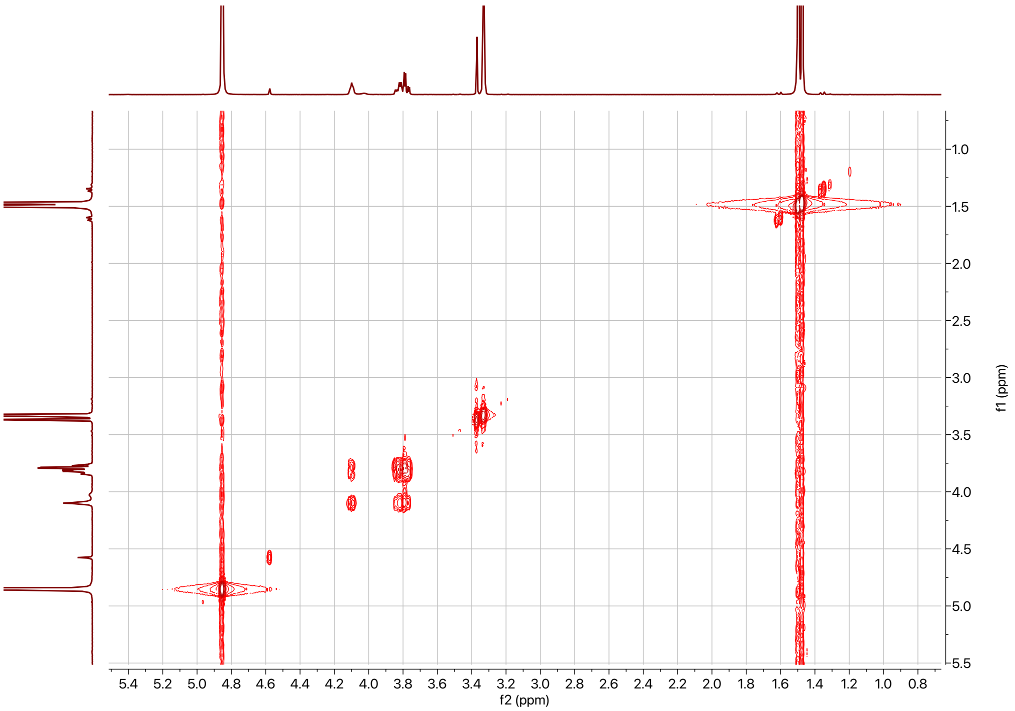


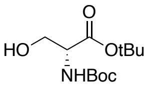


**SI-10**

500 MHz,
HSQC in $CD_3OD$

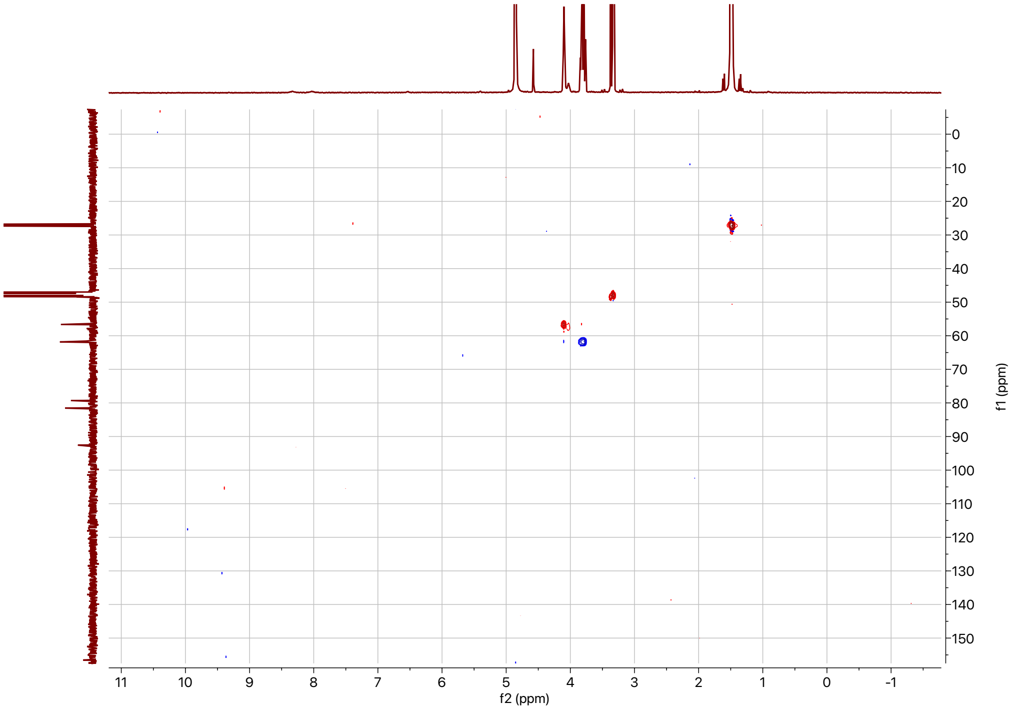

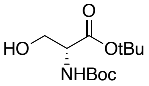


**SI-10**

500 MHz,
HMBC in $CD_3OD$

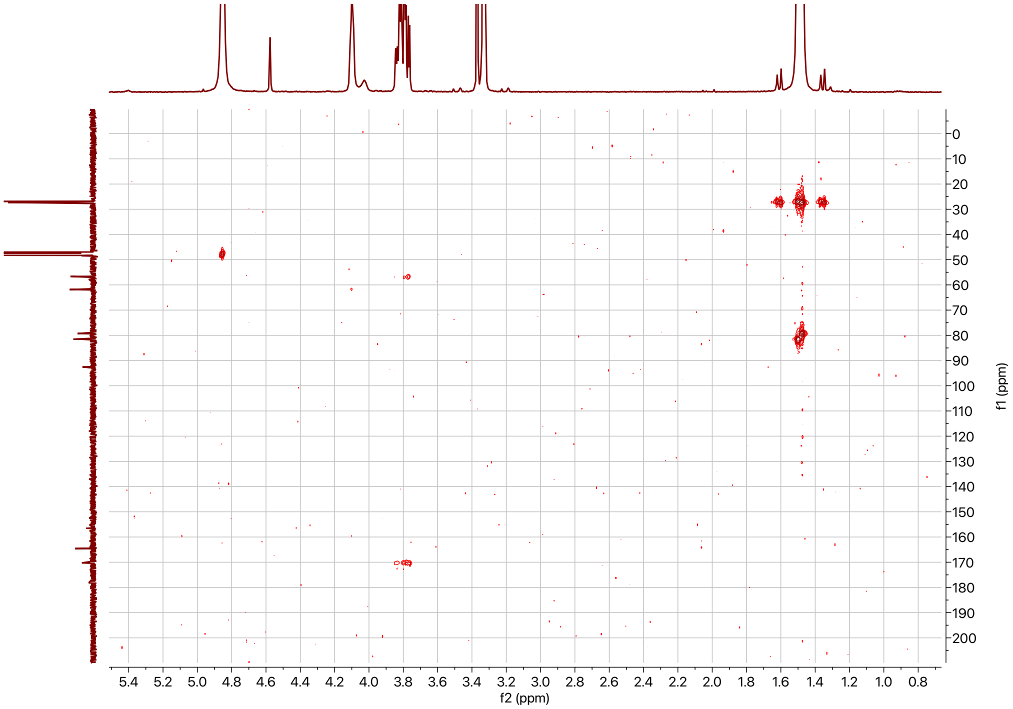

## SI-11:

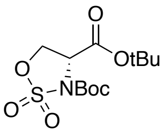


**SI-11**

500 MHz,
$^{1}H$ in $CDCl_3$

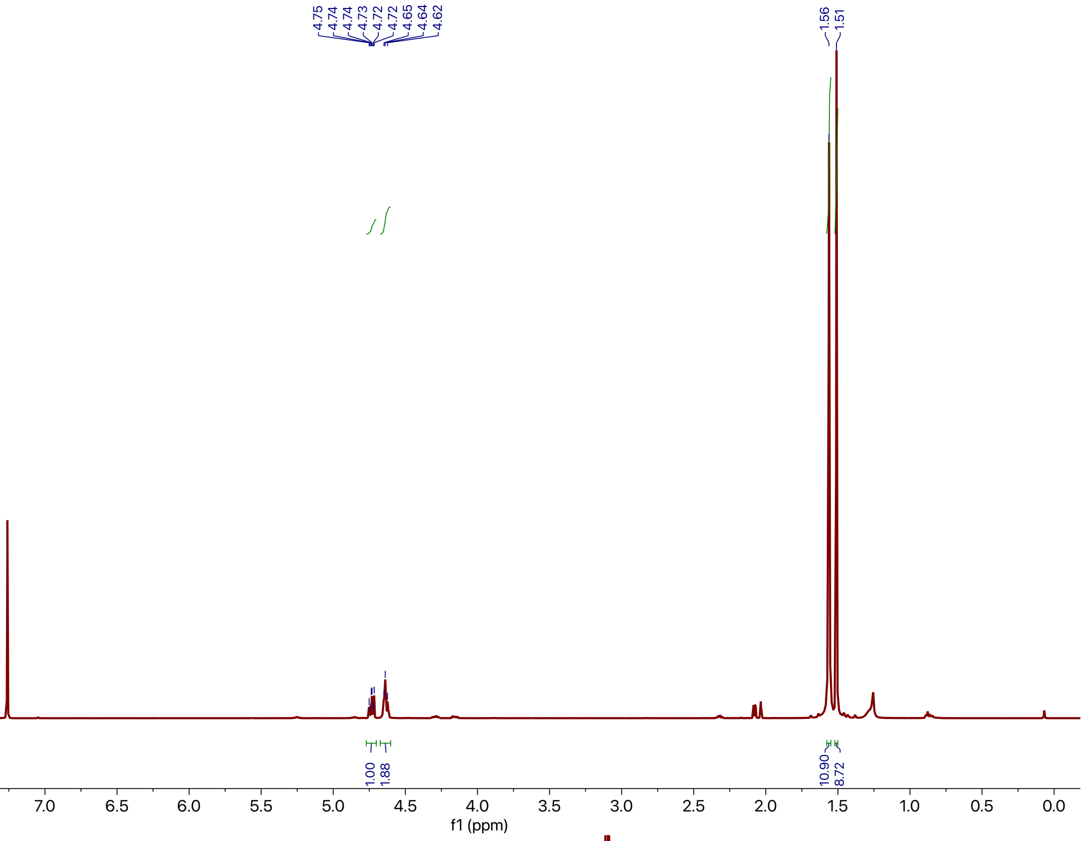


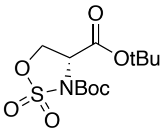


**SI-11**

125 MHz,
$^{13}C$ in $CDCl_3$

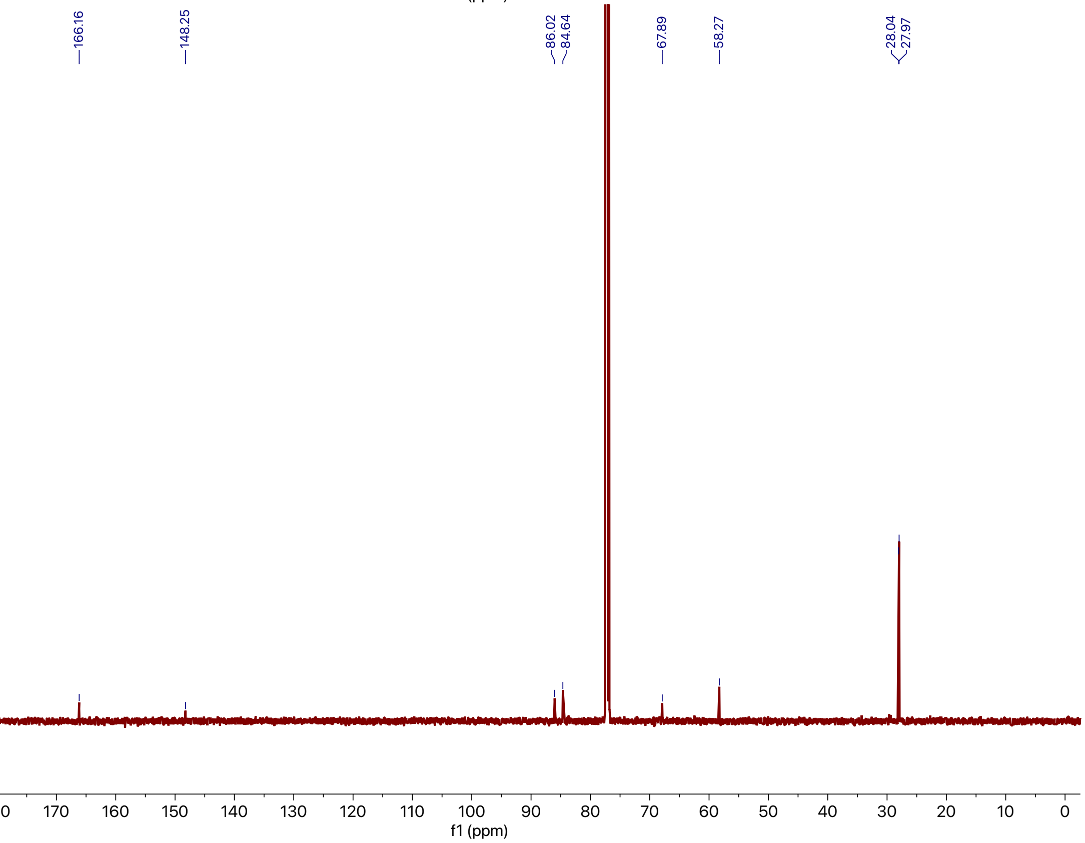

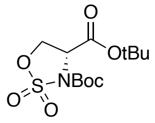


**SI-11**

500 MHz,
COSY in $CDCl_3$

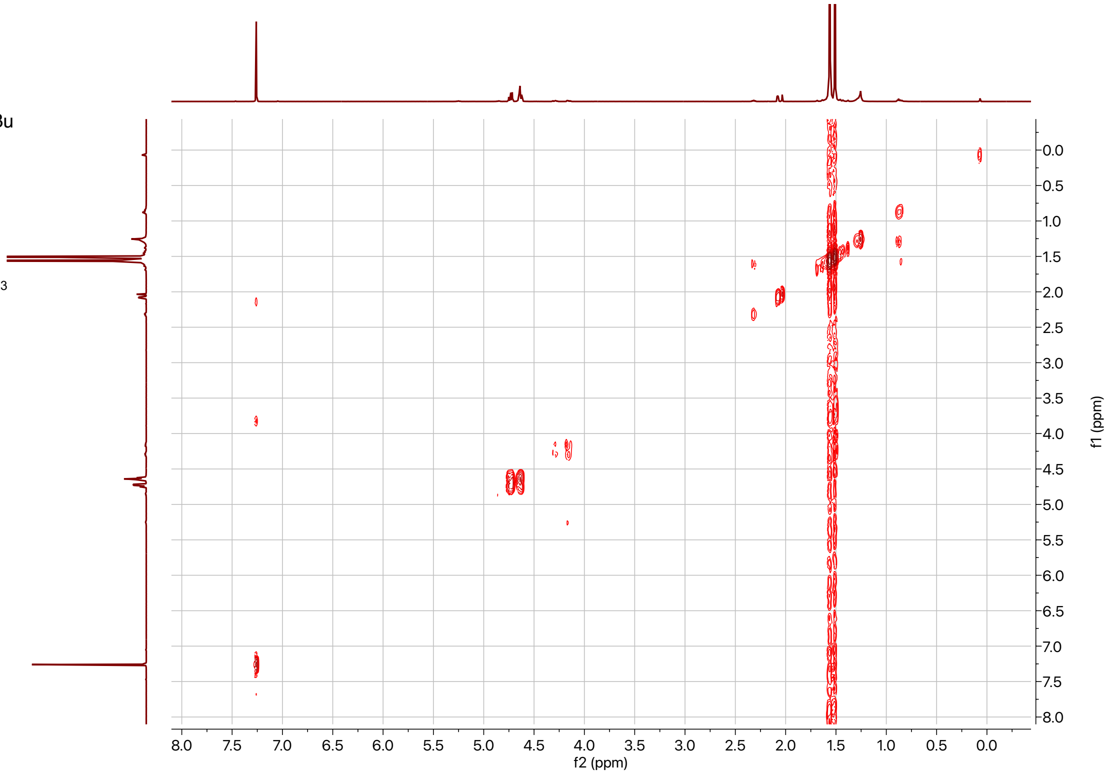


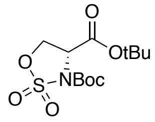


**SI-11**

500 MHz,
HSQC in $CDCl_3$

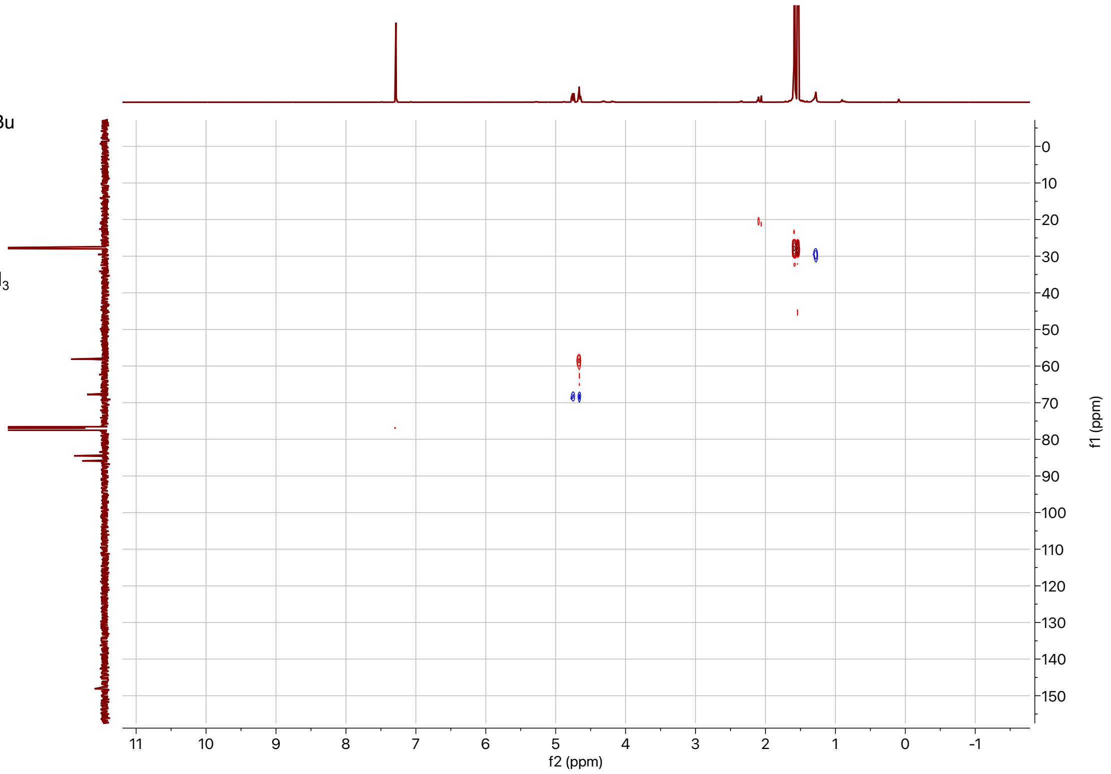

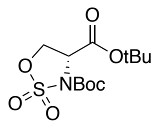

**SI-11**

500 MHz,
HMBC in $CDCl_3$

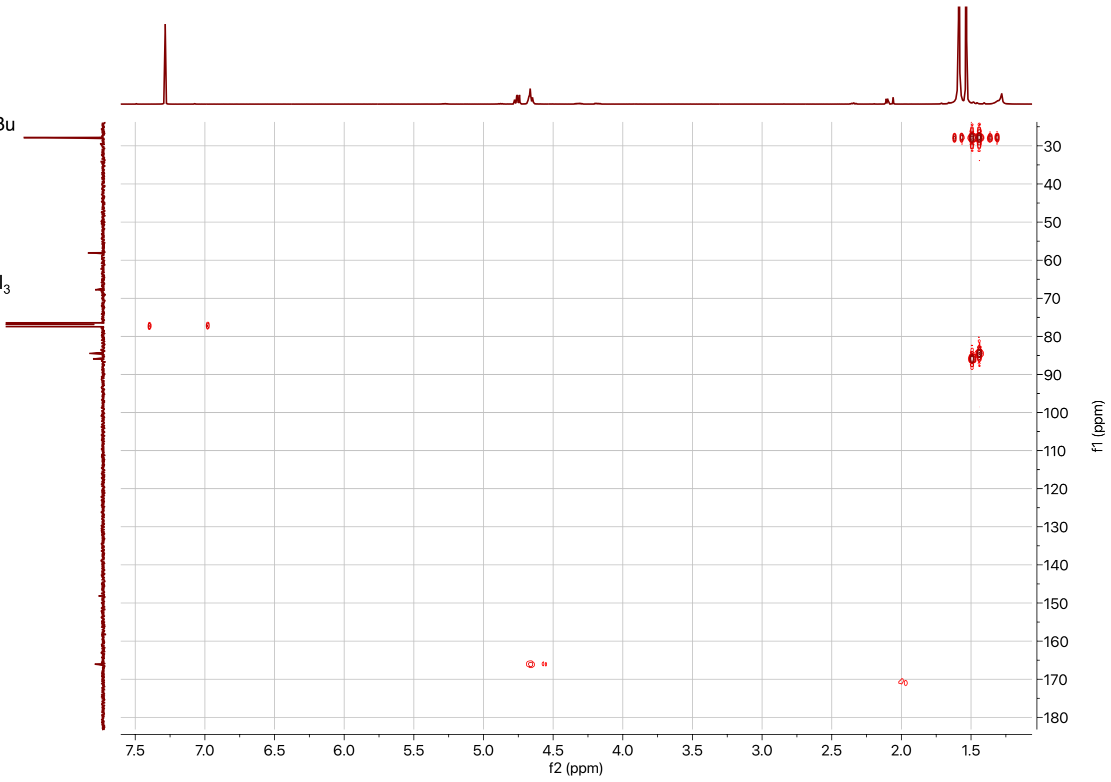

## SI-12:

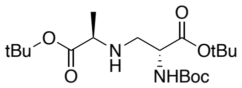


**SI-12**

500 MHz,
$^{1}H$ in $CDCl_3$

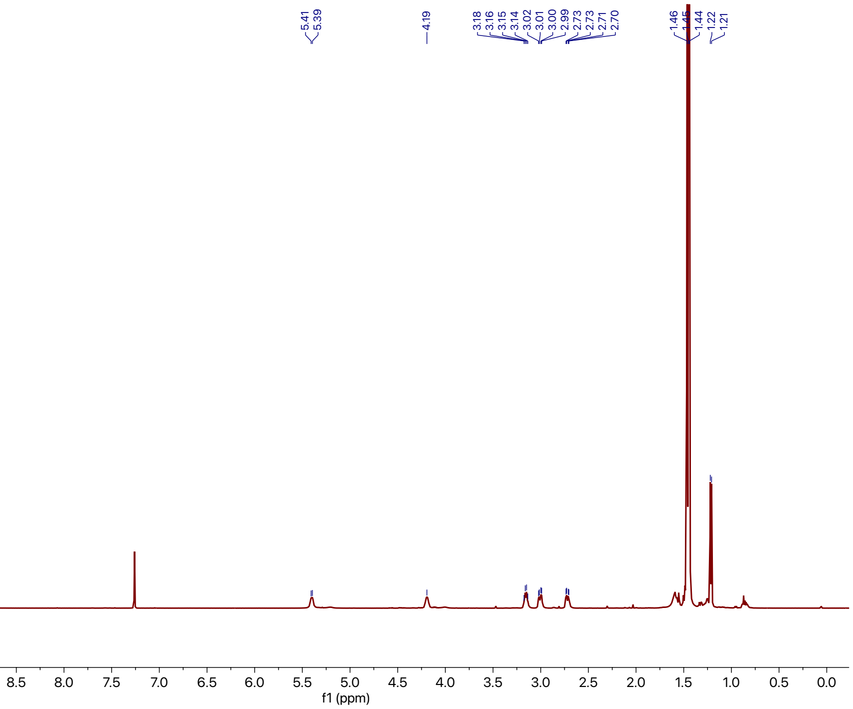


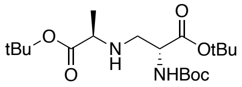


**SI-12**

125 MHz,
$^{13}C$ in $CDCl_3$

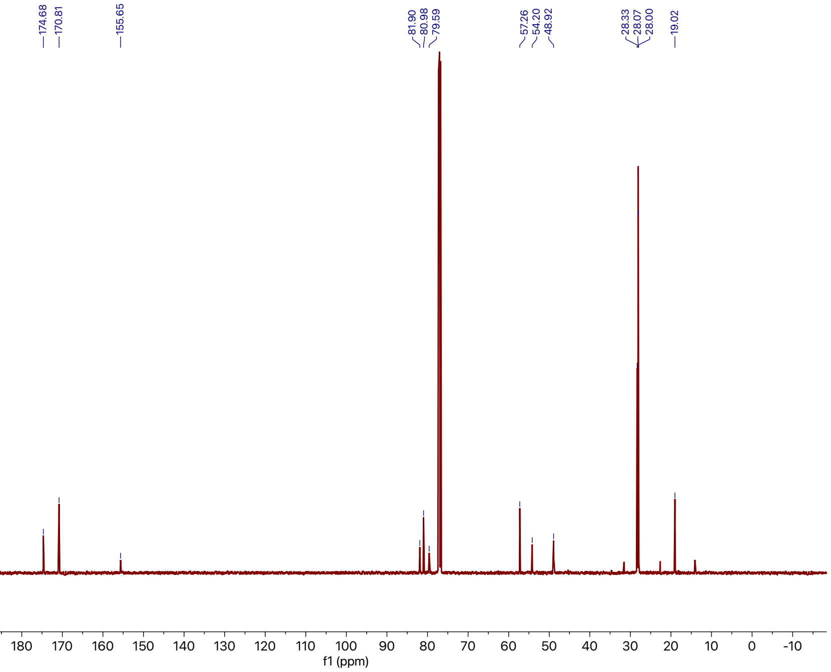

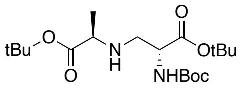

**SI-12**

500 MHz,
COSY in $CDCl_3$

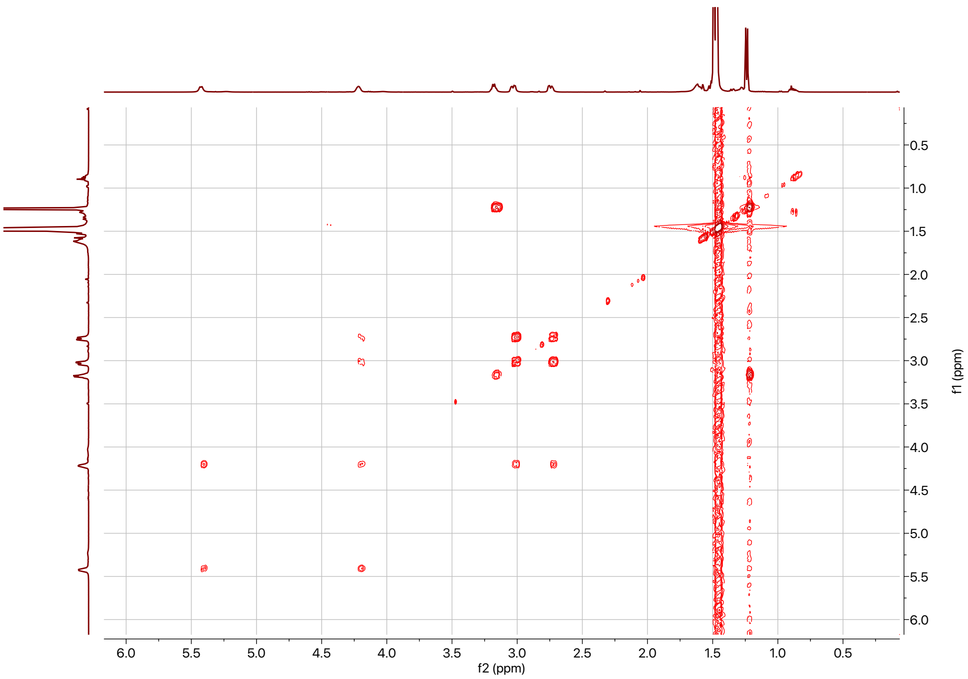


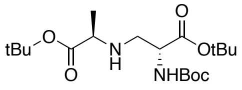

**SI-12**

500 MHz,
HSQC in $CDCl_3$

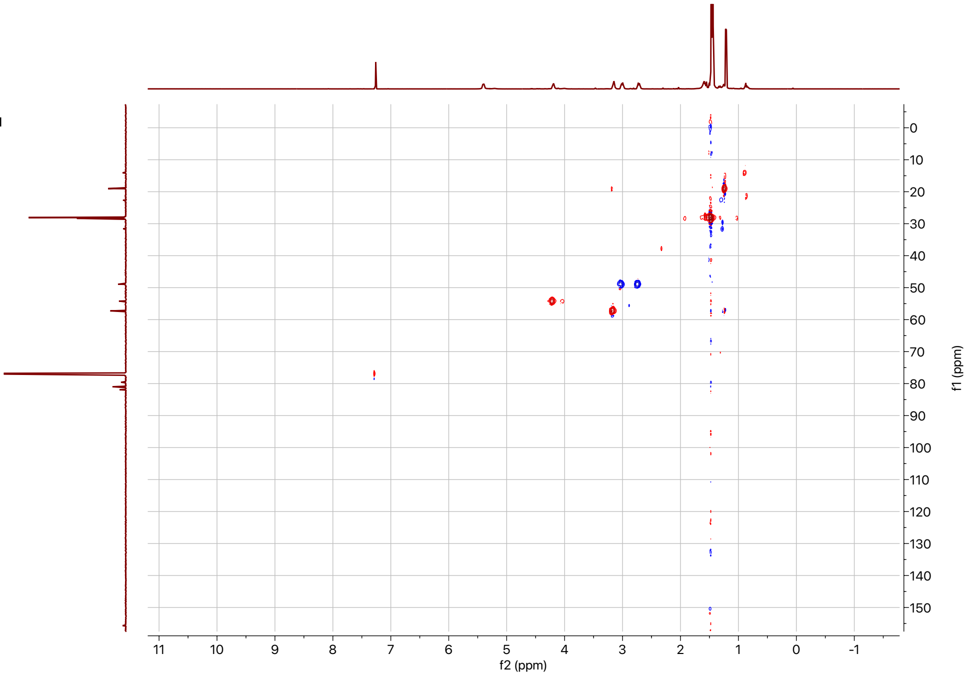

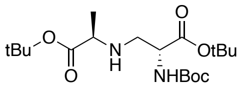


**SI-12**

500 MHz,
HMBC in $CDCl_3$

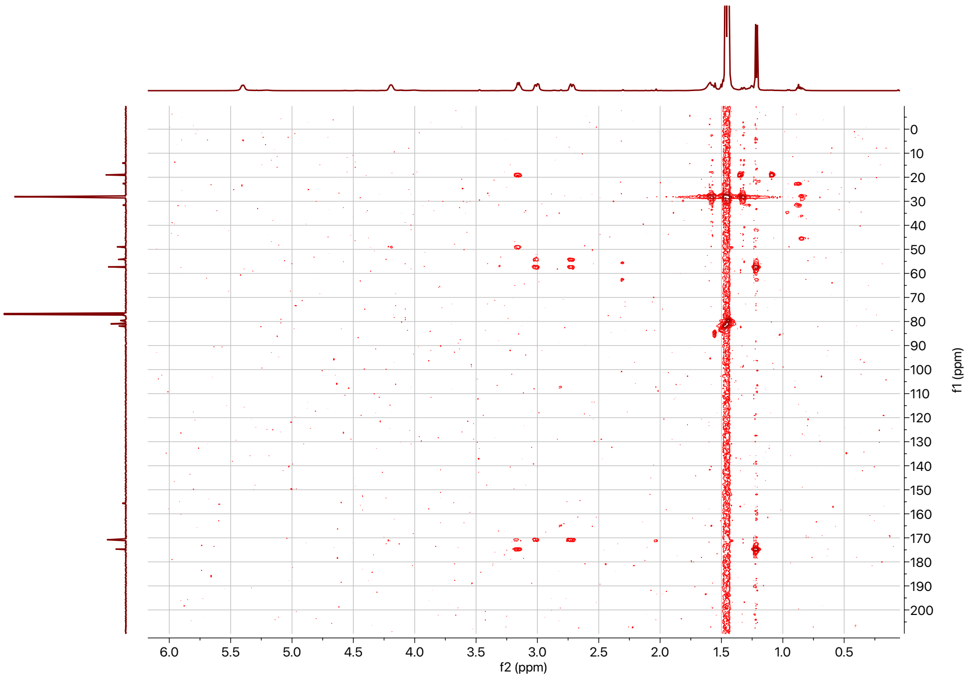

## SI-13:

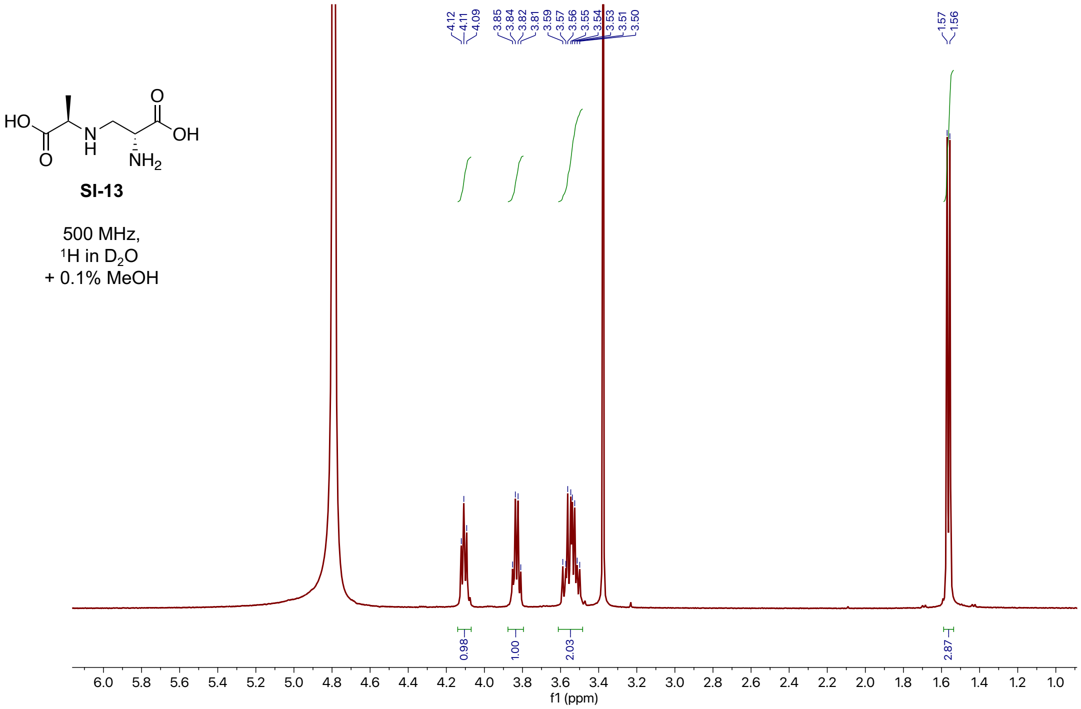


**SI-13**

500 MHz,
$^{1}H$ in $D_2O$
+ 0.1% MeOH

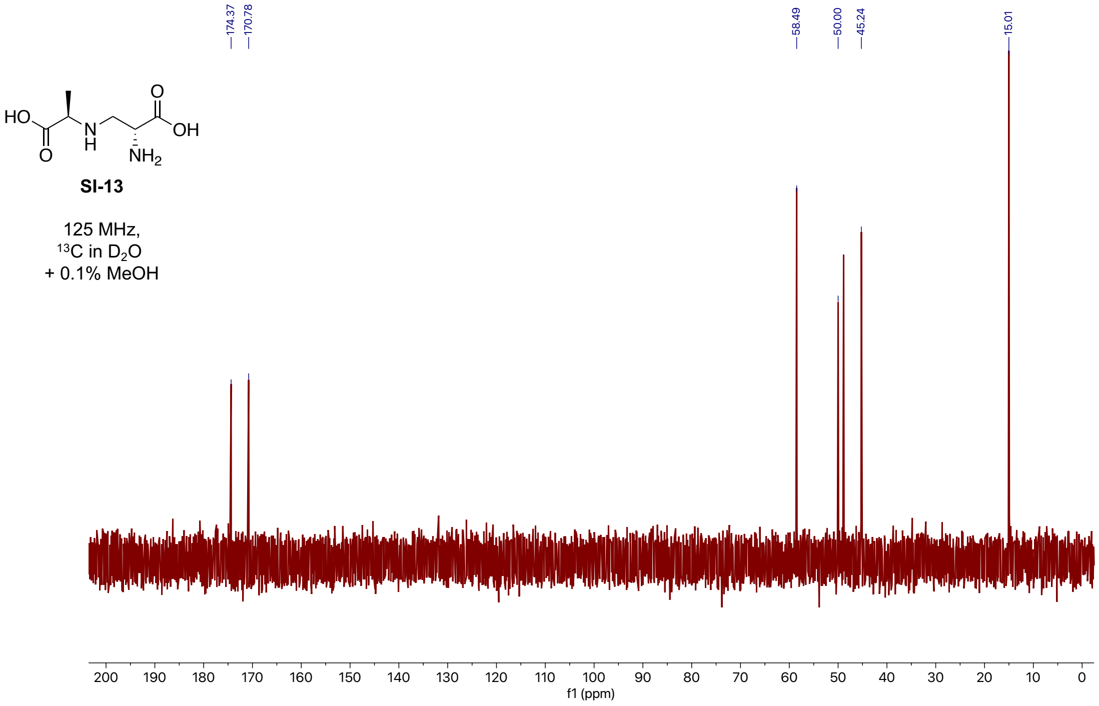


**SI-13**

125 MHz,
$^{13}C$ in $D_2O$
+ 0.1% MeOH

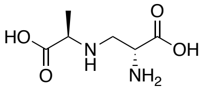


**SI-13**

500 MHz,
COSY in $D_2O$
+ 0.1% MeOH

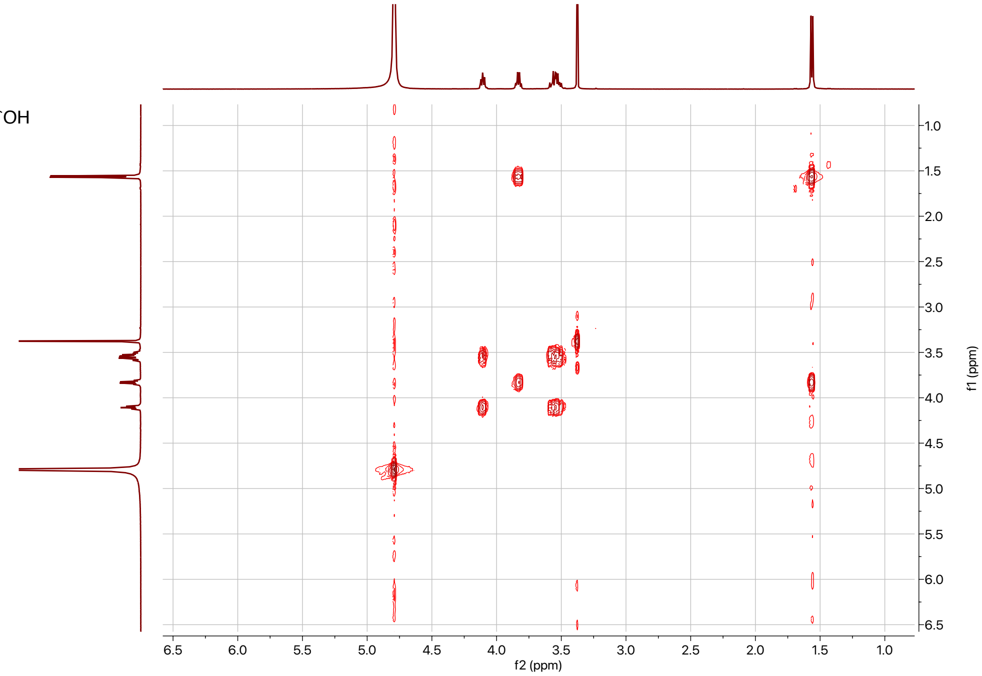


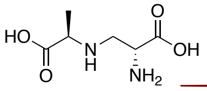


**SI-13**

500 MHz,
HSQC in $D_2O$
+ 0.1% MeOH

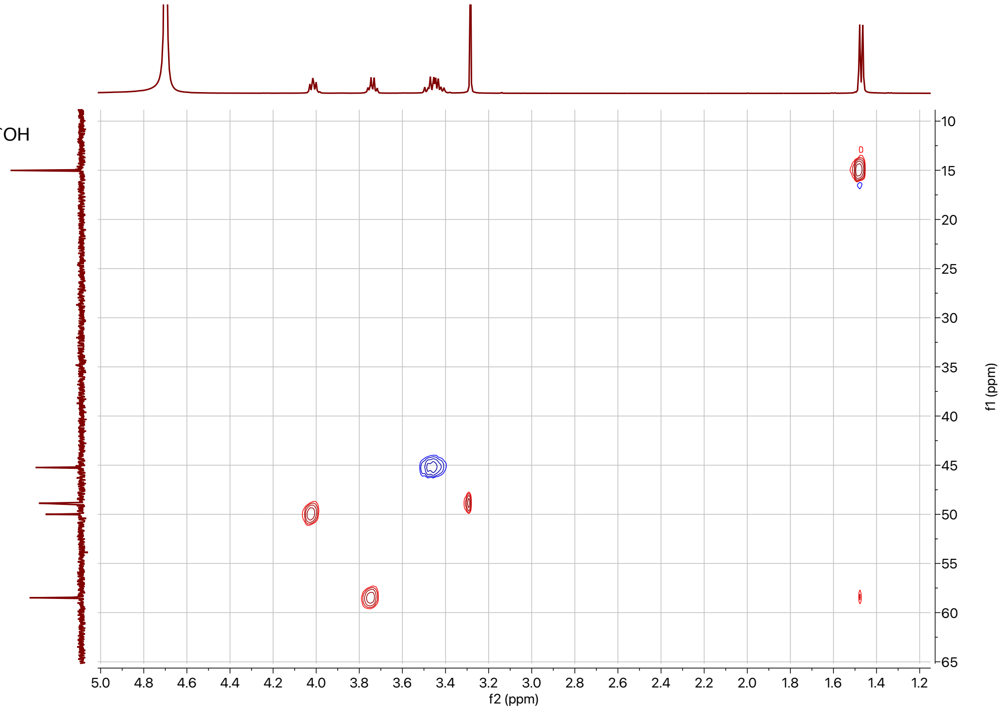

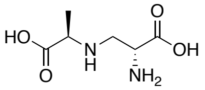


**SI-13**

500 MHz,
HMBC in $D_2O$
+ 0.1% MeOH

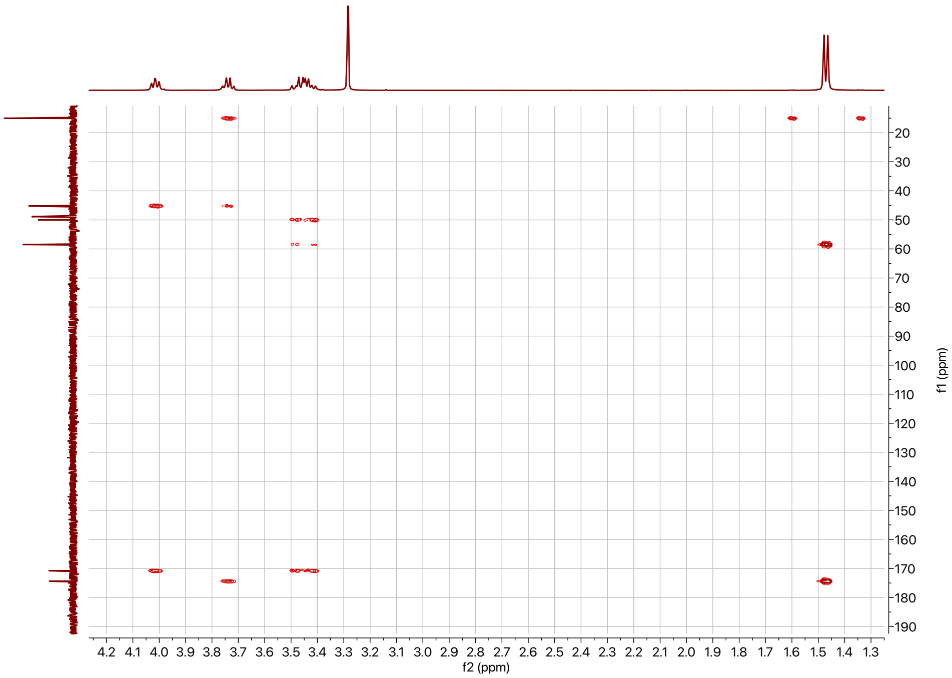

## SI-14:

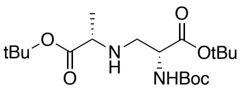


**SI-14**

500 MHz,
$^{1}H$ in $CDCl_3$

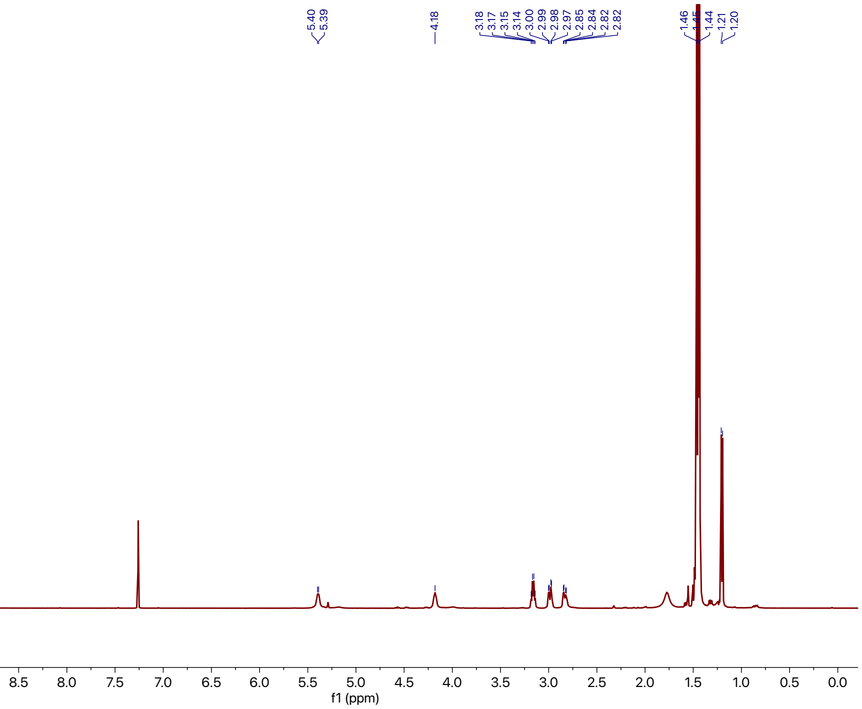


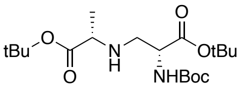


**SI-14**

500 MHz,
$^{13}C$ in $CDCl_3$

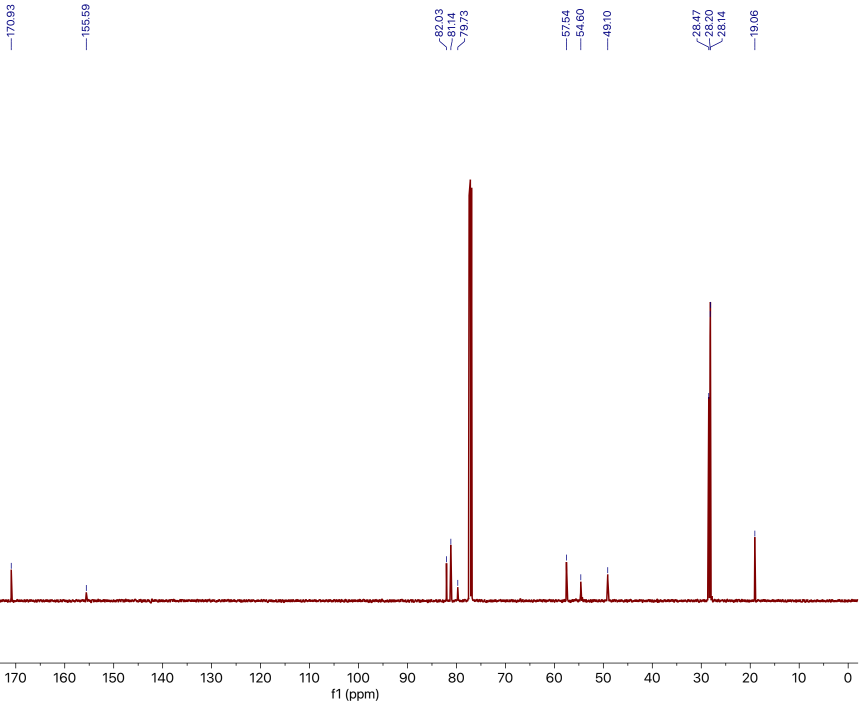

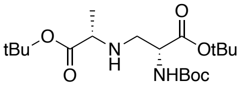


**SI-14**

500 MHz,
COSY in $CDCl_3$

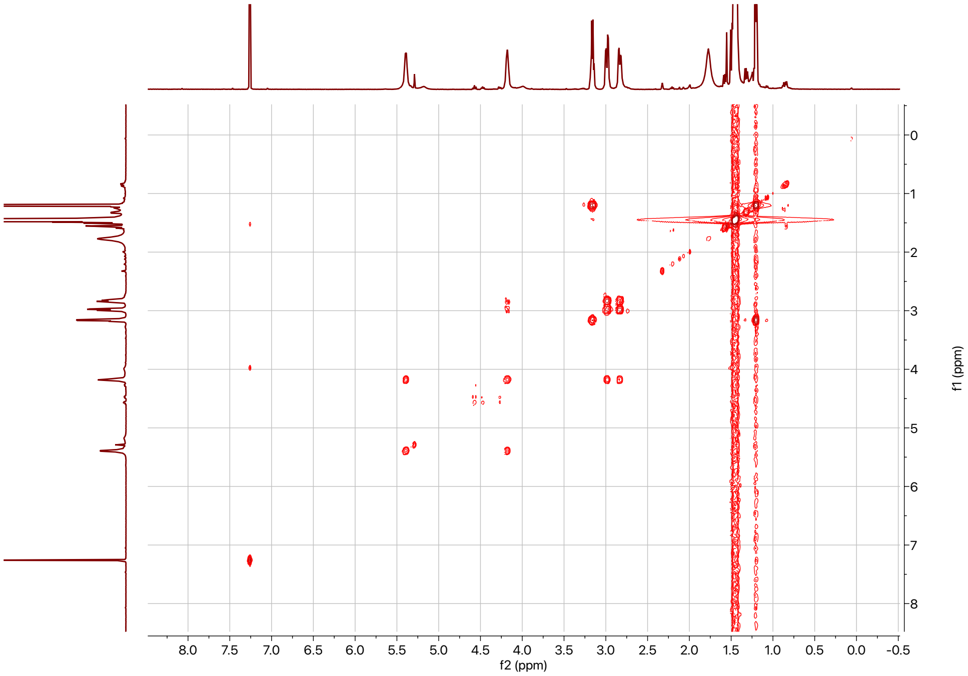


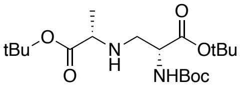


**SI-14**

500 MHz,
HSQC in $CDCl_3$

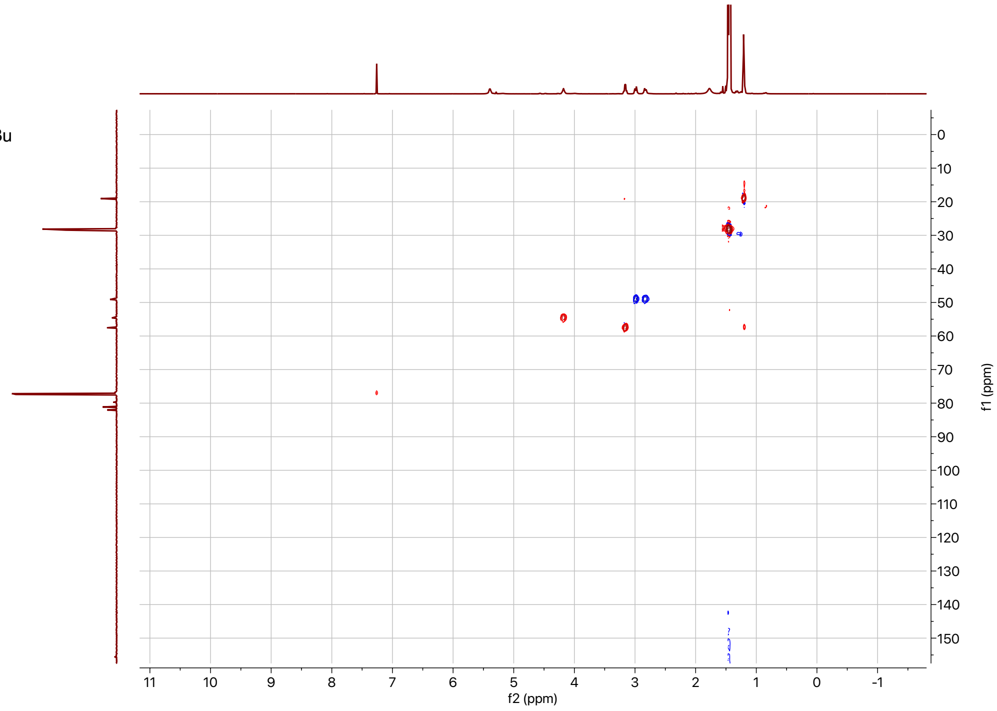

Supplemental Information: **Integrated metabolomic and genomic insights into amino acid incorporation within the hybrid polyketide-alkaloid antibiotic TLN-05220**

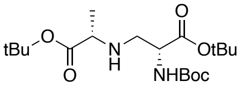

**SI-14**

500 MHz,
HMBC in $CDCl_3$

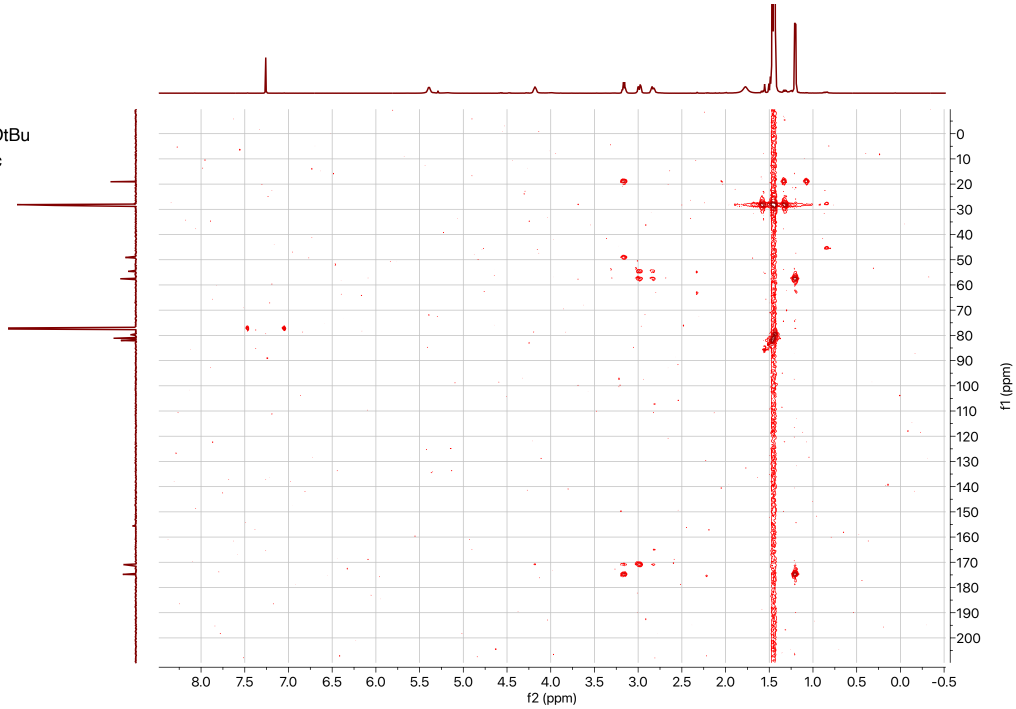

## SI-15:

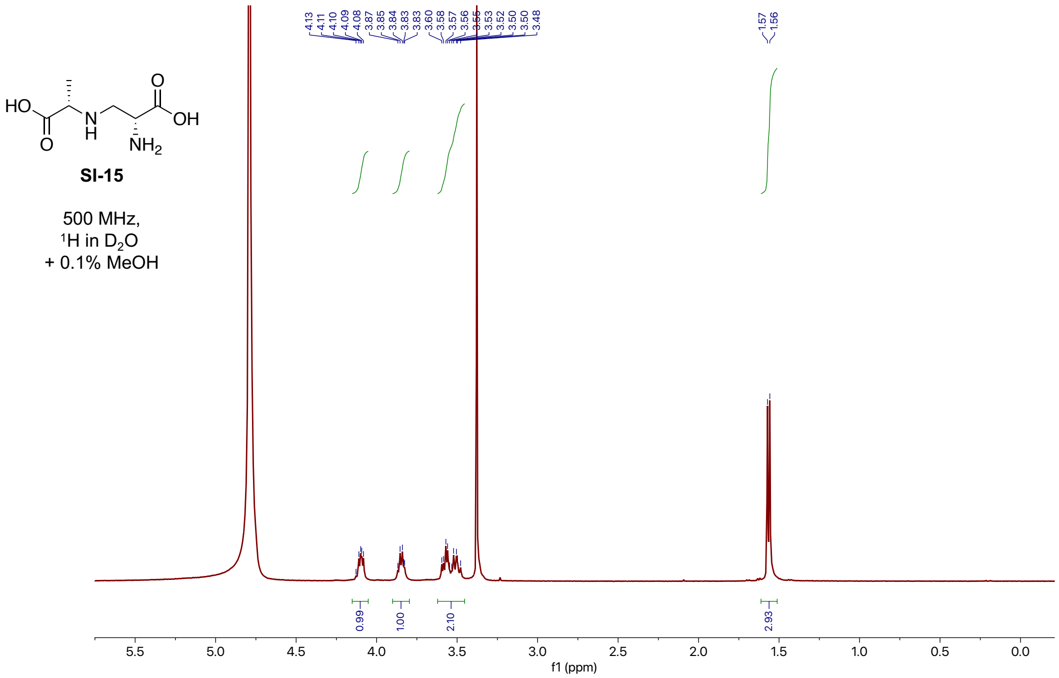
HO
O
N
H
O
OH
NH2
SI-15
500 MHz,
1H in D2O
+ 0.1% MeOH
4.13 4.11 4.10 4.09 4.08 3.87 3.85 3.84 3.83 3.83 3.60 3.58 3.57 3.56 3.55 3.53 3.52 3.50 3.50 3.48
1.57 1.56
0.99
1.00
2.10
2.93
5.5 5.0 4.5 4.0 3.5 3.0 2.5 2.0 1.5 1.0 0.5 0.0
f1 (ppm)

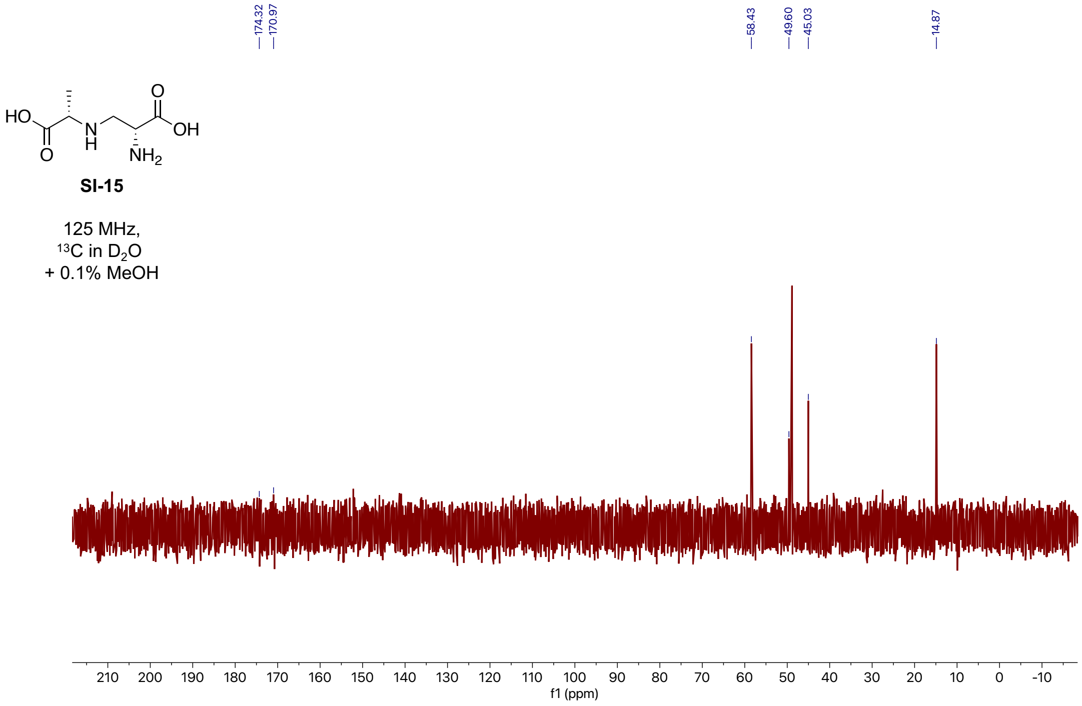
HO
O
N
H
O
OH
NH2
SI-15
125 MHz,
13C in D2O
+ 0.1% MeOH
174.32 170.97
58.43 49.60 45.03
14.87
210 200 190 180 170 160 150 140 130 120 110 100 90 80 70 60 50 40 30 20 10 0 -10
f1 (ppm)

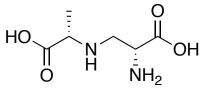


**SI-15**

500 MHz,
COSY in $D_2O$
+ 0.1% MeOH

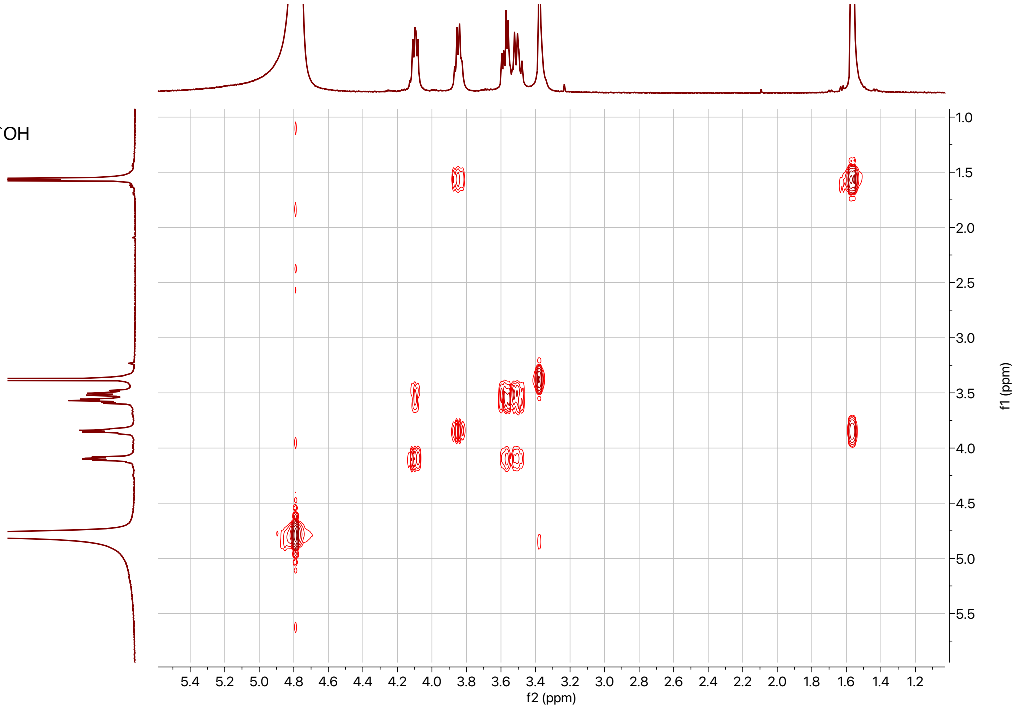


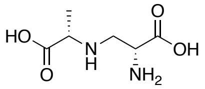


**SI-15**

500 MHz,
HSQC in $D_2O$
+ 0.1% MeOH

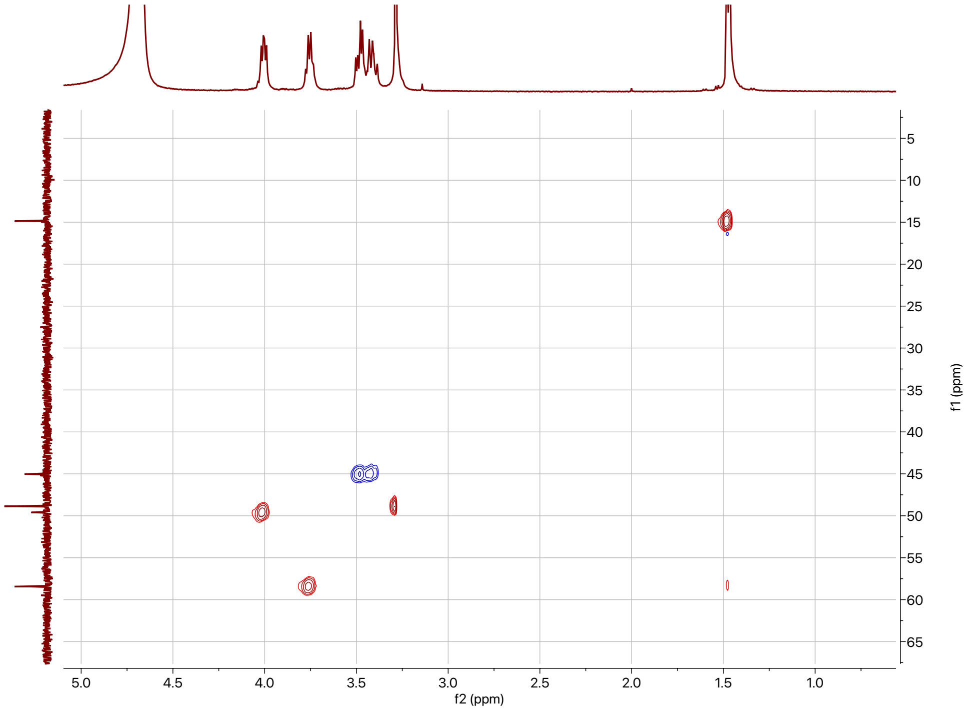

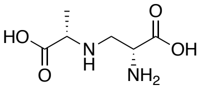

**SI-15**

500 MHz,
HMBC in $D_2O$
+ 0.1% MeOH

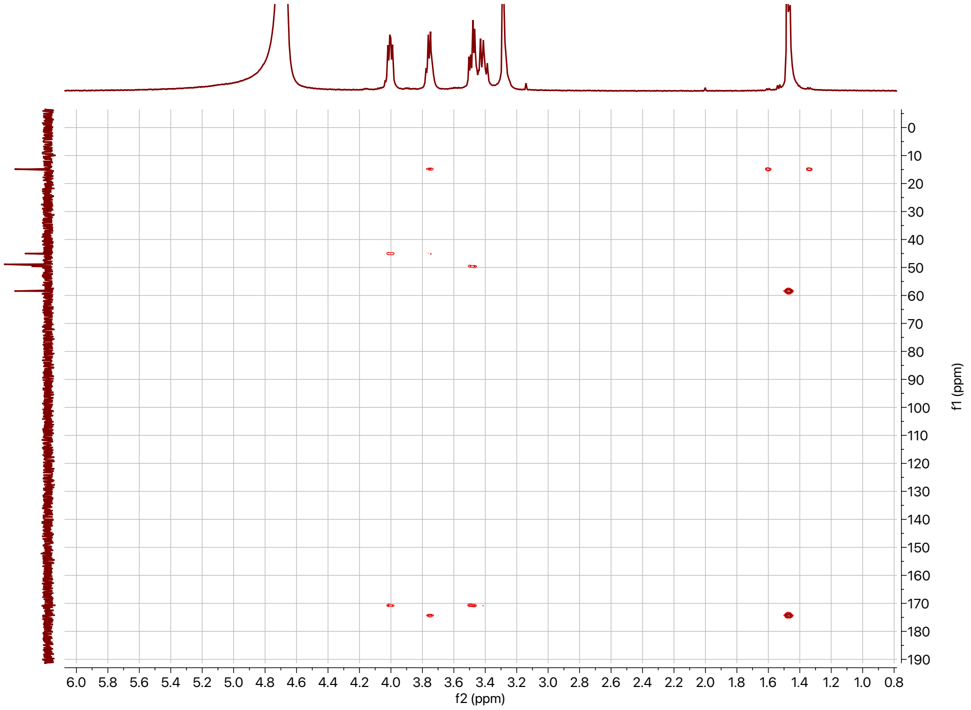

Enzymatic **SI-6**:

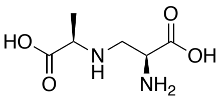


Enzymatic
**SI-6**

500 MHz,
$^1$H in $D_2O$
+ 0.1% MeOH

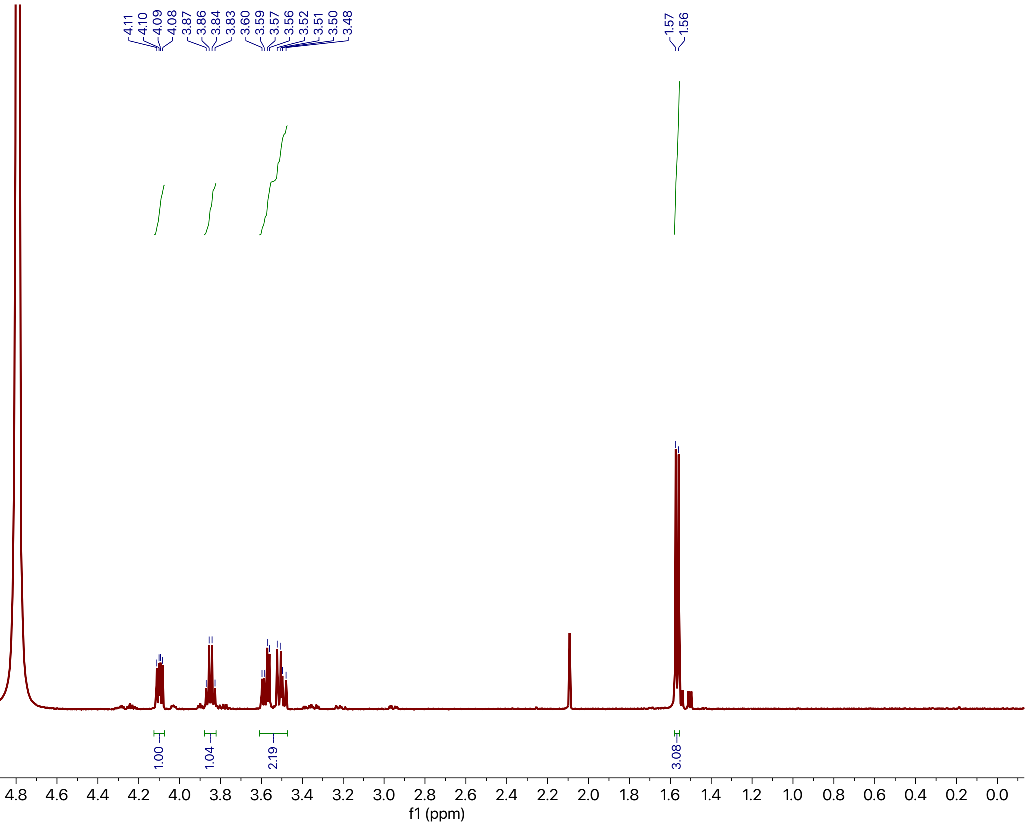


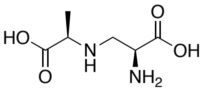


Enzymatic
**SI-6**

125 MHz,
$^{13}$C in $D_2O$
+ 0.1% MeOH

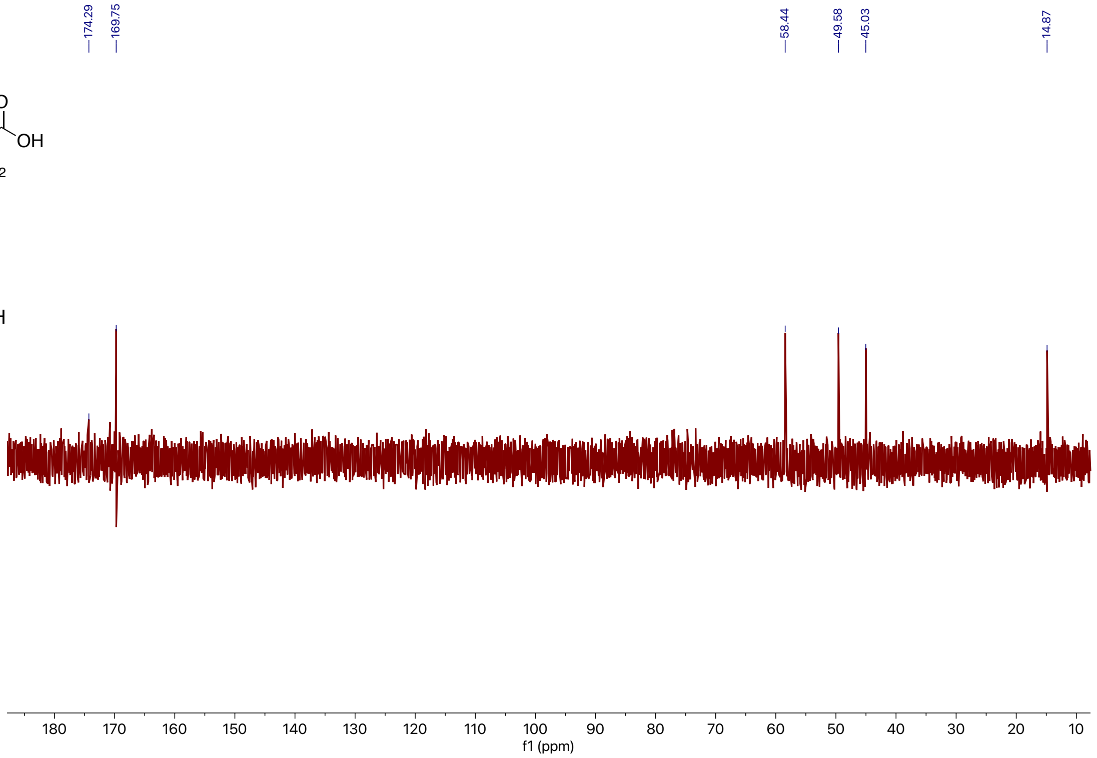

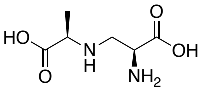


Enzymatic
**SI-6**

500 MHz,
COSY in $D_2O$
+ 0.1% MeOH

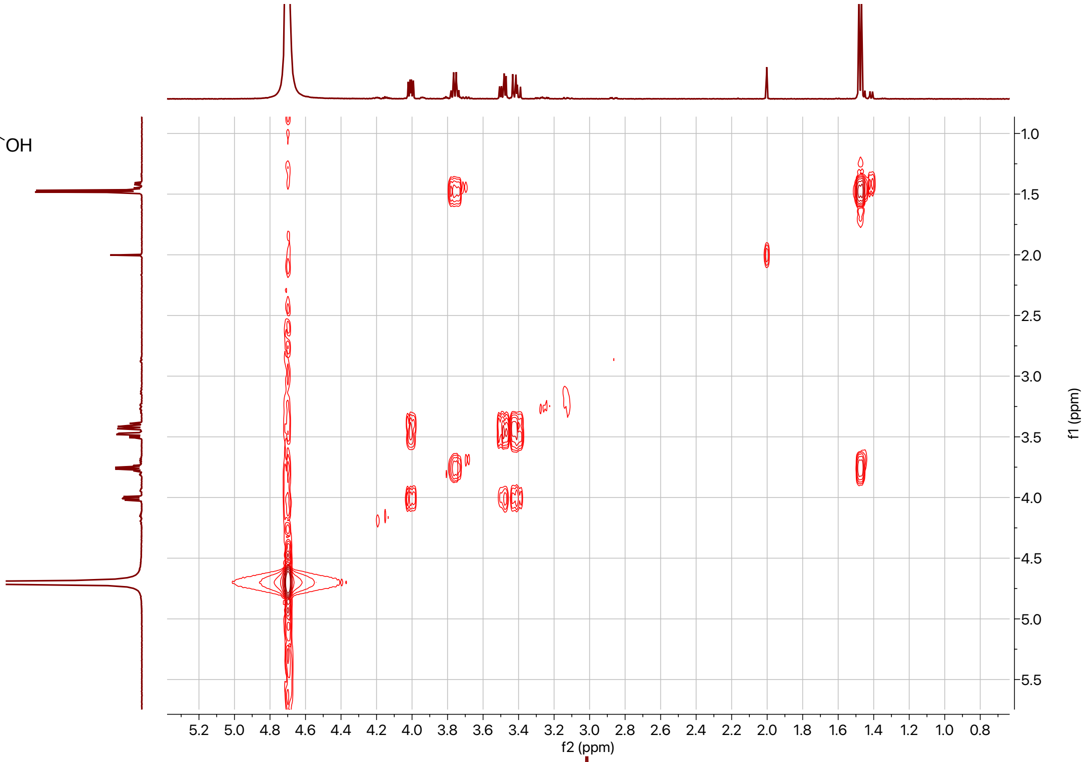


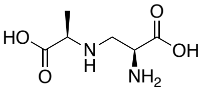


Enzymatic
**SI-6**

500 MHz,
HSQC in $D_2O$
+ 0.1% MeOH

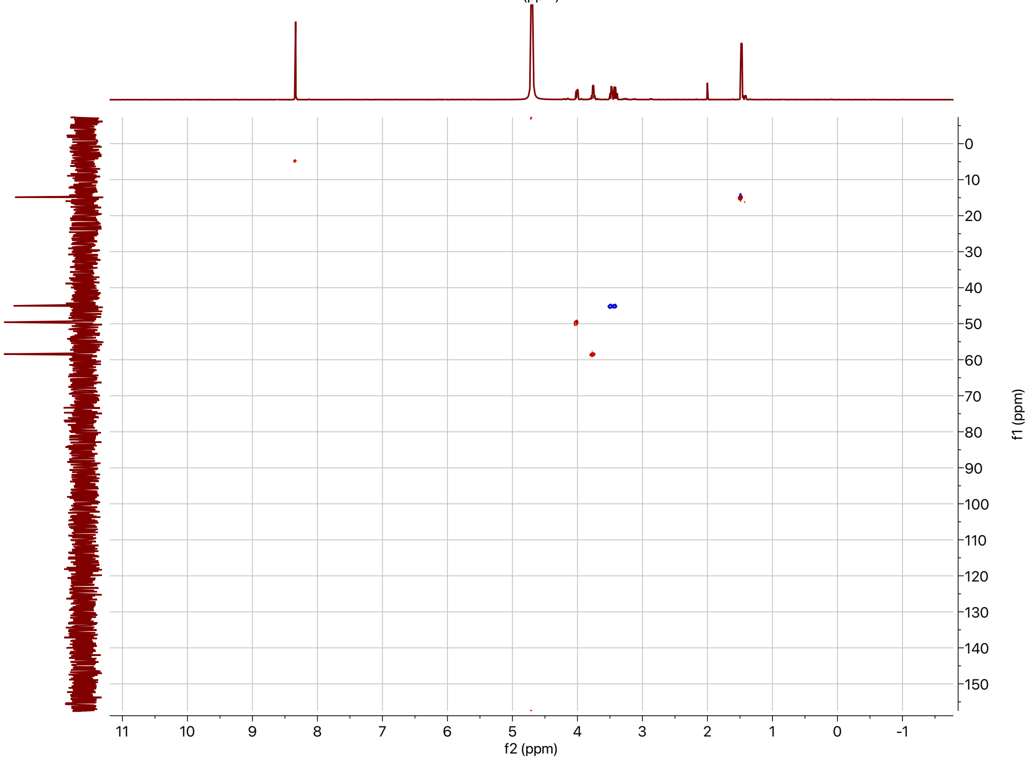

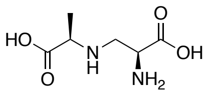

Enzymatic
**SI-6**

500 MHz,
HMBC in $D_2O$
+ 0.1% MeOH

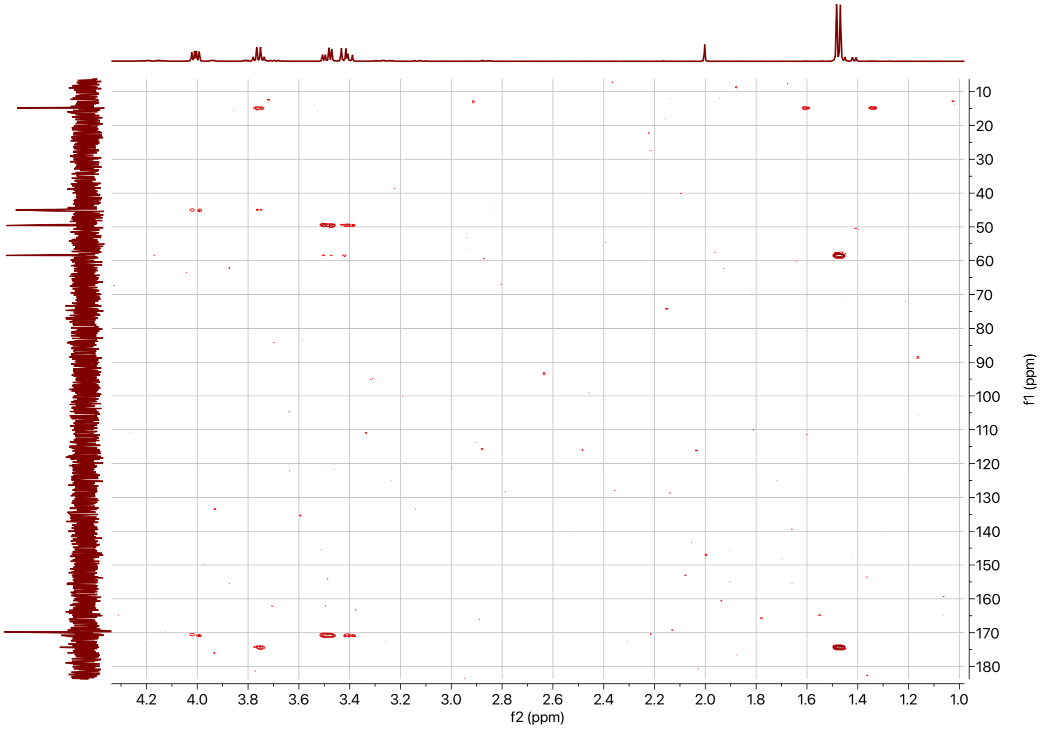

## SI-16:

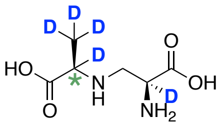


**SI-16**

500 MHz,
$^{1}H$ in $D_2O$
+ 0.1% MeOH

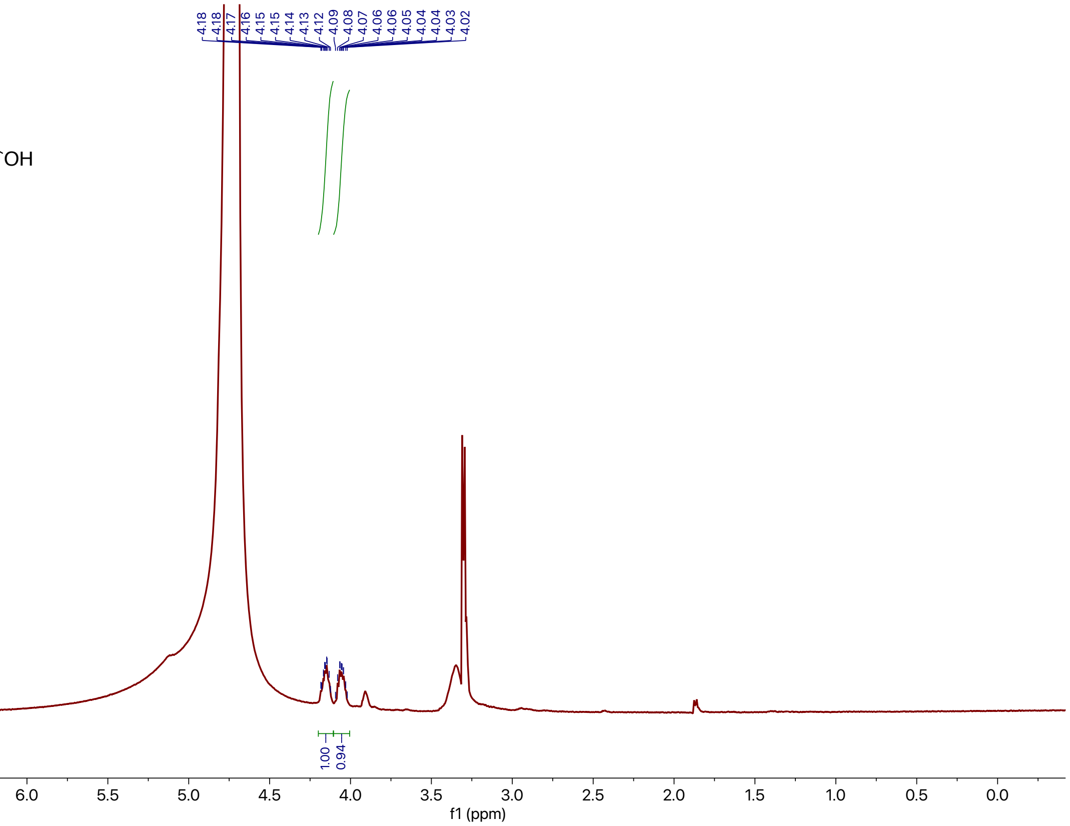


## SI-17:

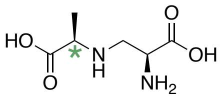


(mixture)
**SI-17**

500 MHz,
$^{1}H$ in $D_2O$
+ 0.1% MeOH

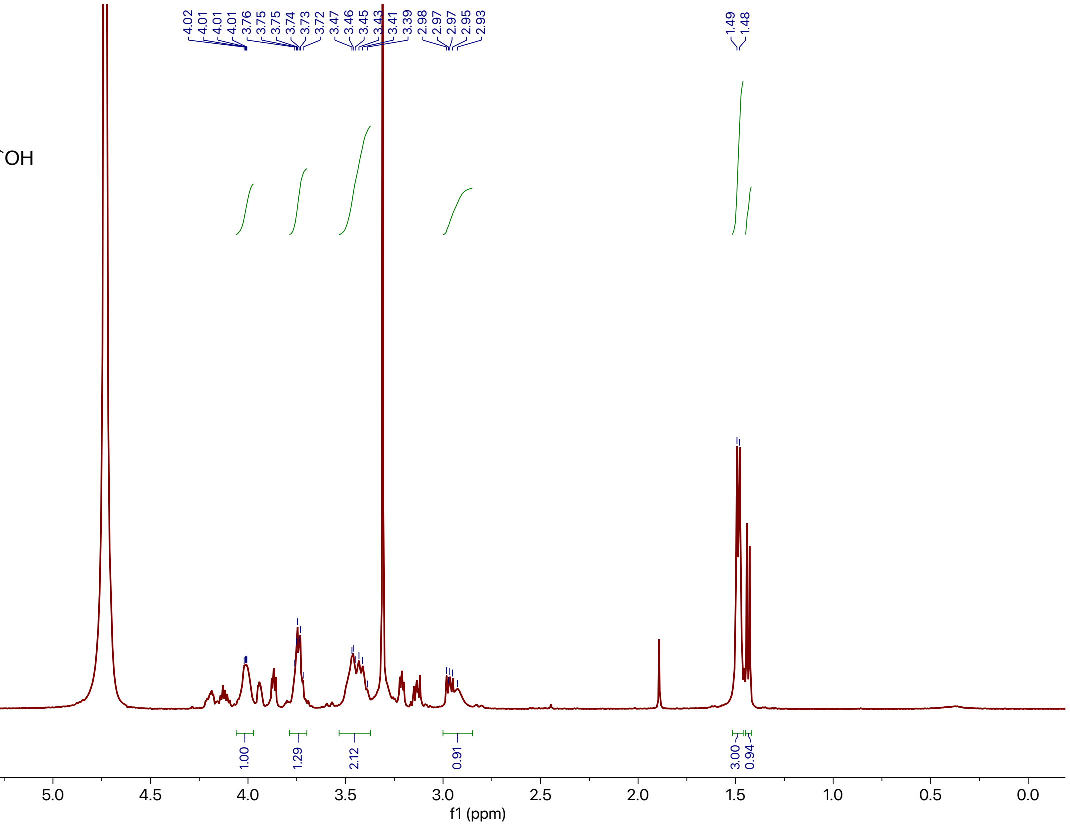

### Supplemental References

(1) Luedemann, G.; Brodsky, B. Taxonomy of gentamicin-producing *Micromonospora*. *Antimicrob. Agents Chemother.* **1996**, *161*, 116–124.
(2) Cordoza, J. L.; Chen, P. Y.-T.; Blaustein, L. R.; Lima, S. T.; Fiore, M. F.; Chekan, J. R.; Moore, B. S.; McKinnie, S. M. K. Mechanistic and Structural Insights into a Divergent PLP-Dependent l-Enduracididine Cyclase from a Toxic Cyanobacterium. *ACS Catal.* **2023**, *13* (14), 9817–9828. https://doi.org/10.1021/acscatal.3c01294.
(3) Guo, Y.; Frisvad, J. C.; Larsen, T. O. Review of Oxepine-Pyrimidinone-Ketopiperazine Type Nonribosomal Peptides. *Metabolites* **2020**, *10* (6), 246. https://doi.org/10.3390/metabo10060246.
(4) Lautru, S.; Gondry, M.; Genet, R.; Pernodet, J.-L. The Albonoursin Gene Cluster of *S. noursei*. *Chem. Biol.* **2002**, *9* (12), 1355–1364. https://doi.org/10.1016/S1074-5521(02)00285-5.
(5) Belin, P.; Moutiez, M.; Lautru, S.; Sequin, J.; Pernodel, J.-L.; Gonry, M. The nonribosomal synthesis of diketopiperazines in tRNA-dependent cyclodipeptide synthase pathways. *Nat. Prod. Rep.* **2012**, *29* (9), 961. https://doi.org/10.1039/c2np20010d.
(6) Yu, X.; Liu, F.; Zou, Y.; Tang, M.-C.; Hang, L.; Houk, K. N.; Tang, Y. Biosynthesis of Strained Piperazine Alkaloids: Uncovering the Concise Pathway of Herquline A. *J. Am. Chem. Soc.* **2016**, *138* (41), 13529–13532. https://doi.org/10.1021/jacs.6b09464.
(7) Zhu, L.-L.; Yang, Q.; Wang, D.-G.; Niu, L.; Pan, Z.; Li, S.; Li, Y.-Z.; Zhang, W.; Wu, C. Deciphering the Biosynthesis and Physiological Function of 5-Methylated Pyrazinones Produced by Myxobacteria. *ACS Cent. Sci.* **2024**, *10* (3), 555–568. https://doi.org/10.1021/acscentsci.3c01363.
(8) Tietze, A.; Shi, Y.; Kronenwerth, M.; Bode, H. B. Nonribosomal Peptides Produced by Minimal and Engineered Synthetases with Terminal Reductase Domains. *ChemBioChem* **2020**, *21* (19), 2750–2754. https://doi.org/10.1002/cbic.202000176.
(9) Zdouc, M. M.; Blin, K.; Louwen, N. L. L.; Navarro, J.; Loureiro, C.; Bader, C. D.; Bailey, C. B.; Barra, L.; Booth, T. J.; Bozhüyük, K. A. J.; et al. MIBiG 4.0: advancing biosynthetic gene cluster curation through global collaboration. *Nucleic Acids Res.* **2025**, *53* (D1), D678–D690. https://doi.org/10.1093/nar/gkae1115.
(10) Banskota, A. H.; Aouidate, M.; Sørensen, D.; Ibrahim, A.; Piraee, M.; Zazopoulos, E.; Alarco, A. M.; Gourdeau, H.; Mellon, C.; Farnet, C. M.; et al. TLN-05220, TLN-05223, new Echinosporamicin-type antibiotics, and proposed revision of the structure of bravomicins. *J. Antibiot. (Tokyo)* **2009**, *62* (10), 565–570. https://doi.org/10.1038/ja.2009.77.
(11) Blin, K.; Shaw, S.; Vader, L.; Szenei, J.; Reitz, Z. L.; Augustijn, H. E.; Cediel-Becerra, J. D. D.; de Crécy-Lagard, V.; Koetsier, R. A.; Williams, S. E.; et al. antiSMASH 8.0: extended gene cluster detection capabilities and analyses of chemistry, enzymology, and regulation. *Nucleic Acids Res.* **2025**, *53* (W1), W32–W38. https://doi.org/10.1093/nar/gkaf334.
(12) Gilchrist, C. L. M.; Chooi, Y.-H. clinker & clustermap.js: automatic generation of gene cluster comparison figures. *Bioinformatics* **2021**, *37* (16), 2473–2475. https://doi.org/10.1093/bioinformatics/btab007.
(13) Skinnider, M. A.; Johnston, C. W.; Gunabalasingam, M.; Merwin, N. J.; Kieliszek, A. M.; MacLellan, R. J.; Li, H.; Ranieri, M. R. M.; Webster, A. L. H.; Cao, M. P. T.; et al. Comprehensive prediction of secondary metabolite structure and biological activity from microbial genome sequences. *Nat. Commun.* **2020**, *11* (1), 6058. https://doi.org/10.1038/s41467-020-19986-1.
(14) Del Vecchio, F.; Petkovic, H.; Kendrew, S. G.; Low, L.; Wilkinson, B.; Lill, R.; Cortés, J.; Rudd, B. A. M.; Staunton, J.; Leadlay, P. F. Active-site residue, domain and module swaps in modular polyketide synthases. *J. Ind. Microbiol. Biotechnol.* **2003**, *30* (8), 489–494. https://doi.org/10.1007/s10295-003-0062-0.
(15) Chisuga, T.; Nagai, A.; Miyanaga, A.; Goto, E.; Kishikawa, K.; Kudo, F.; Eguchi, T. Structural Insight into the Reaction Mechanism of Ketosynthase-Like Decarboxylase in a Loading Module of Modular Polyketide Synthases. *ACS Chem. Biol.* **2022**, *17* (1), 198–206. https://doi.org/10.1021/acschembio.1c00856.
(16) Moore, B. S.; Hertweck, C. Biosynthesis and attachment of novel bacterial polyketide synthase starter units. *Nat. Prod. Rep.* **2002**, *19* (1), 70–99. https://doi.org/10.1039/b003939j.

(17) Villebro, R.; Shaw, S.; Blin, K.; Weber, T. Sequence-based classification of type II polyketide synthase biosynthetic gene clusters for antiSMASH. *J. Ind. Microbiol. Biotechnol.* **2019**, *46* (3–4), 469–475. https://doi.org/10.1007/s10295-018-02131-9.
(18) Bililign, T.; Hyun, C.-G.; Williams, J. S.; Czisny, A. M.; Thorson, J. S. The Hedamycin Locus Implicates a Novel Aromatic PKS Priming Mechanism. *Chem. Biol.* **2004**, *11* (7), 959–969. https://doi.org/10.1016/j.chembiol.2004.04.016.
(19) Fitzgerald, J. T.; Charkoudian, L. K.; Watts, K. R.; Khosla, C. Analysis and refactoring of the A-74528 biosynthetic pathway. *J. Am. Chem. Soc.* **2013**, *135* (10), 3752–3755. https://doi.org/10.1021/ja311579s.
(20) Zaleta-Rivera, K.; Charkoudian, L. K.; Ridley, C. P.; Khosla, C. Cloning, sequencing, heterologous expression, and mechanistic analysis of A-74528 biosynthesis. *J. Am. Chem. Soc.* **2010**, *132* (26), 9122–9128. https://doi.org/10.1021/ja102519v.
(21) Das, A.; Khosla, C. In Vivo and In Vitro Analysis of the Hedamycin Polyketide Synthase. *Chem. Biol.* **2009**, *16* (11), 1197–1207. https://doi.org/10.1016/j.chembiol.2009.11.005.
(22) Lackner, G.; Schenk, A.; Xu, Z.; Reinhardt, K.; Yunt, Z. S.; Piel, J.; Hertweck, C. Biosynthesis of Pentangular Polyphenols: Deductions from the Benastatin and Griseorhodin Pathways. *J. Am. Chem. Soc.* **2007**, *129* (30), 9306–9312. https://doi.org/10.1021/ja0718624.
(23) Zhang, W.; Wang, L.; Kong, L.; Wang, T.; Chu, Y.; Deng, Z.; You, D. Unveiling the post-PKS redox tailoring steps in biosynthesis of the type II polyketide antitumor antibiotic xantholipin. *Chem. Biol.* **2012**, *19* (3), 422–432. https://doi.org/10.1016/j.chembiol.2012.01.016.
(24) Zhan, J.; Watanabe, K.; Tang, Y. Synergistic actions of a monooxygenase and cyclases in aromatic polyketide biosynthesis. *ChemBioChem* **2008**, *9*, 1710–1715. https://doi.org/10.1002/cbic.200800178.
(25) Caldara-Festin, G.; Jackson, D. R.; Barajas, J. F.; Valentic, T. R.; Patel, A. B.; Aguilar, S.; Nguyen, M.; Vo, M.; Khanna, A.; Sasaki, E.; et al. Structural and functional analysis of two di-domain aromatase/cyclases from type II polyketide synthases. *Proc. Natl. Acad. Sci.* **2015**, *112* (50). https://doi.org/10.1073/pnas.1512976112.
(26) Ogasawara, Y.; Yackley, B. J.; Greenberg, J. A.; Rogelj, S.; Melancon, C. E. Expanding our understanding of sequence-function relationships of Type II polyketide biosynthetic gene clusters: Bioinformatics-guided identification of frankiamicin a from *Frankia* sp. EAN1pec. *PLoS ONE* **2015**, *10* (4), 1–25. https://doi.org/10.1371/journal.pone.0121505.
(27) Napan, K.; Zhang, S.; Morgan, W.; Anderson, T.; Takemoto, J. Y.; Zhan, J. Synergistic Actions of Tailoring Enzymes in Pradimicin Biosynthesis. *ChemBioChem* **2014**, *15* (15), 2289–2296. https://doi.org/10.1002/cbic.201402306.
(28) Chen, Y.; Wendt-Pienkoski, E.; Rajski, S. R.; Shen, B. In vivo investigation of the roles of FdmM and FdmM1 in fredericamycin biosynthesis unveiling a new family of oxygenases. *J. Biol. Chem.* **2009**, *284* (37), 24735–24743. https://doi.org/10.1074/jbc.M109.014191.
(29) Kudo, F.; Yonezawa, T.; Komatsubara, A.; Mizoue, K.; Eguchi, T. Cloning of the biosynthetic gene cluster for naphthoxanthene antibiotic FD-594 from *Streptomyces* sp. TA-0256. *J. Antibiot. (Tokyo)* **2011**, *64* (1), 123–132. https://doi.org/10.1038/ja.2010.145.
(30) McAlpine, J. B.; Banskota, A. H.; Charan, R. D.; Schlingmann, G.; Zazopoulos, E.; Piraee, M.; Janso, J.; Bernan, V. S.; Aouidate, M.; Farnet, C. M.; et al. Biosynthesis of Diazepinomicin/ECO-4601, a *Micromonospora* Secondary Metabolite with a Novel Ring System. *J. Nat. Prod.* **2008**, *71* (9), 1585–1590. https://doi.org/10.1021/np800376n.
(31) Zhan, J.; Qiao, K.; Tang, Y. Investigation of Tailoring Modifications in Pradimicin Biosynthesis. *ChemBioChem* **2009**, *10* (9), 1447–1452. https://doi.org/10.1002/cbic.200900082.
(32) Banskota, A. H.; Aouidate, M.; Sørensen, D.; Ibrahim, A.; Piraee, M.; Zazopoulos, E.; Alarco, A. M.; Gourdeau, H.; Mellon, C.; Farnet, C. M.; et al. TLN-05220, TLN-05223, new Echinosporamicin-type antibiotics, and proposed revision of the structure of bravomicins. *J. Antibiot. (Tokyo)* **2009**, *62* (10), 565–570. https://doi.org/10.1038/ja.2009.77.
(33) Kim, B. C.; Lee, J. M.; Ahn, J. S.; Kim, B. S. Cloning, sequencing, and characterization of the pradimicin biosynthetic gene cluster of *Actinomadura hibisca* P157-2. *J. Microbiol. Biotechnol.* **2007**, *17*, 830–839.

(34) Kang, H. S.; Brady, S. F. Arixanthomycins A-C: Phylogeny-guided discovery of biologically active eDNA-derived pentangular polyphenols. *ACS Chem. Biol.* **2014**, *9* (6), 1267–1272. https://doi.org/10.1021/cb500141b.
(35) Monciardini, P.; Bernasconi, A.; Iorio, M.; Brunati, C.; Sosio, M.; Campochiaro, L.; Landini, P.; Maffioli, S. I.; Donadio, S. Antibacterial Aromatic Polyketides Incorporating the Unusual Amino Acid Enduracididine. *J. Nat. Prod.* **2019**, *82* (1), 35–44. https://doi.org/10.1021/acs.jnatprod.8b00354.
(36) Wendt-Pienkowski, E.; Huang, Y.; Zhang, J.; Li, B.; Jiang, H.; Kwon, H.; Hutchinson, C. R.; Shen, B. Cloning, sequencing, analysis, and heterologous expression of the fredericamycin biosynthetic gene cluster from *Streptomyces griseus*. *J. Am. Chem. Soc.* **2005**, *127* (3), 16442–16452. https://doi.org/10.1021/ja054376u.
(37) Zhang, W.; Wang, L.; Kong, L.; Wang, T.; Chu, Y.; Deng, Z.; You, D. Unveiling the post-PKS redox tailoring steps in biosynthesis of the type II polyketide antitumor antibiotic xantholipin. *Chem. Biol.* **2012**, *19* (3), 422–432. https://doi.org/10.1016/j.chembiol.2012.01.016.
(38) Lopez, P.; Hornung, A.; Welzel, K.; Unsin, C.; Wohlleben, W.; Weber, T.; Pelzer, S. Isolation of the lysolipin gene cluster of *Streptomyces tendae* Tu 4042. *Gene* **2010**, *461* (1–2), 5–14. https://doi.org/10.1016/j.gene.2010.03.016.
(39) Hu, X.; Sun, W.; Li, S.; Li, L.; Yu, L.; Liu, H.; You, X.; Jiang, B.; Wu, L. Cervinomycins C1-4 with cytotoxic and antibacterial activity from *Streptomyces* sp. CPCC 204980. *J. Antibiot. (Tokyo)* **2020**, *73* (12), 812–817. https://doi.org/10.1038/s41429-020-0342-1.
(40) Kersten, R. D.; Ziemert, N.; Gonzalez, D. J.; Duggan, B. M.; Nizet, V.; Dorrestein, P. C.; Moore, B. S. Glycogenomics as a mass spectrometry-guided genome-mining method for microbial glycosylated molecules. *Proc. Natl. Acad. Sci.* **2013**, *110* (47). https://doi.org/10.1073/pnas.1315492110.
(41) Kang, H.-S.; Brady, S. F. Mining Soil Metagenomes to Better Understand the Evolution of Natural Product Structural Diversity: Pentangular Polyphenols as a Case Study. *J. Am. Chem. Soc.* **2014**, *136* (52), 18111–18119. https://doi.org/10.1021/ja510606j.
(42) Xu, Z.; Schenk, A.; Hertweck, C. Molecular Analysis of the Benastatin Biosynthetic Pathway and Genetic Engineering of Altered Fatty Acid−Polyketide Hybrids. *J. Am. Chem. Soc.* **2007**, *129* (18), 6022–6030. https://doi.org/10.1021/ja069045b.
(43) Guo, Q.; Wu, D.; Gao, L.; Bai, Y.; Liu, Y.; Guo, N.; Du, X.; Yang, J.; Wang, X.; Lei, X. Identification of the AMA Synthase from the Aspergillomarasmine A Biosynthesis and Evaluation of Its Biocatalytic Potential. *ACS Catal.* **2020**, *10* (11), 6291–6298. https://doi.org/10.1021/acscatal.0c01187.
(44) Buller, A. R.; Brinkmann-Chen, S.; Romney, D. K.; Herger, M.; Murciano-Calles, J.; Arnold, F. H. Directed evolution of the tryptophan synthase β-subunit for stand-alone function recapitulates allosteric activation. *Proc. Natl. Acad. Sci.* **2015**, *112* (47), 14599–14604. https://doi.org/10.1073/pnas.1516401112.
(45) Chen, Y.; Wendt-Pienkowski, E.; Ju, J.; Lin, S.; Rajski, S. R.; Shen, B. Characterization of FdmV as an amide synthetase for fredericamycin A biosynthesis in *Streptomyces griseus* ATCC 43944. *J. Biol. Chem.* **2010**, *285* (50), 38853–38860. https://doi.org/10.1074/jbc.M110.147744.
(46) Database resources of the National Center for Biotechnology Information. *Nucleic Acids Res.* **2014**, *42* (D1), D7–D17. https://doi.org/10.1093/nar/gkt1146.
(47) Zhu, X.; Siitonen, V.; Melançon Iii, C. E.; Metsä-Ketelä, M. Biosynthesis of Diverse Type II Polyketide Core Structures in *Streptomyces coelicolor* M1152. *ACS Synth. Biol.* **2021**, *10* (2), 243–251. https://doi.org/10.1021/acssynbio.0c00482.
(48) Camacho, C.; Coulouris, G.; Avagyan, V.; Ma, N.; Papadopoulos, J.; Bealer, K.; Madden, T. L. BLAST+: architecture and applications. *BMC Bioinformatics* **2009**, *10* (1), 421. https://doi.org/10.1186/1471-2105-10-421.
(49) Blum, M.; Andreeva, A.; Florentino, L. C.; Chuguransky, S. R.; Grego, T.; Hobbs, E.; Pinto, B. L.; Orr, A.; Paysan-Lafosse, T.; Ponamareva, I.; et al. InterPro: the protein sequence classification resource in 2025. *Nucleic Acids Res.* **2025**, *53* (D1), D444–D456. https://doi.org/10.1093/nar/gkae1082.
(50) Bastard, K.; Perret, A.; Mariage, A.; Bessonnet, T.; Pinet-Turpault, A.; Petit, J.-L.; Darii, E.; Bazire, P.; Vergne-Vaxelaire, C.; Brewee, C.; et al. Parallel evolution of non-homologous isofunctional enzymes in methionine biosynthesis. *Nat. Chem. Biol.* **2017**, *13* (8), 858–866. https://doi.org/10.1038/nchembio.2397.
(51) Hsu, S.-H.; Zhang, S.; Huang, S.-C.; Wu, T.-K.; Xu, Z.; Chang, C.-Y. Characterization of Enzymes Catalyzing the Formation of the Nonproteinogenic Amino Acid l-Dap in Capreomycin Biosynthesis. *Biochemistry* **2021**, *60* (1), 77–84. https://doi.org/10.1021/acs.biochem.0c00808.

(52) Albu, S. A.; Koteva, K.; King, A. M.; Al-Karmi, S.; Wright, G. D.; Capretta, A. Total Synthesis of Aspergillomarasmine A and Related Compounds: A Sulfamidate Approach Enables Exploration of Structure–Activity Relationships. *Angew. Chem. Int. Ed.* **2016**, *55* (42), 13259–13262. https://doi.org/10.1002/anie.201606657.

(53) Koshizuka, M.; Makino, K.; Shimada, N. Diboronic Acid Anhydride-Catalyzed Direct Peptide Bond Formation Enabled by Hydroxy-Directed Dehydrative Condensation. *Org. Lett.* **2020**, *22* (21), 8658–8664. https://doi.org/10.1021/acs.orglett.0c03252.
